# Disease-specific tau filaments assemble via polymorphic intermediates

**DOI:** 10.1101/2023.07.24.550295

**Authors:** Sofia Lövestam, David Li, Jane L. Wagstaff, Abhay Kotecha, Dari Kimanius, Stephen H. McLaughlin, Alexey G. Murzin, Stefan M.V. Freund, Michel Goedert, Sjors H.W. Scheres

**Author notes:** These authors jointly supervised this project.

## Abstract

Intermediate species in the assembly of amyloid filaments are believed to play a central role in neurodegenerative diseases and may constitute important targets for therapeutic intervention. However, structural information about intermediate species has been scarce and the molecular mechanisms by which amyloids assemble remain largely unknown. Here, we use time-resolved electron cryo-microscopy (cryo-EM) to study the *in vitro* assembly of recombinant truncated tau (amino acids 297-391) into paired helical filaments of Alzheimer’s disease or into filaments of chronic traumatic encephalopathy. We report the formation of a shared first intermediate amyloid (FIA), with an ordered core comprising amino acids 302-316. Nuclear magnetic resonance indicates that the same amino acids adopt rigid, β-strand-like conformations in monomeric tau. At later time points, the FIAs disappear and we observe many different intermediate amyloid filaments, with structures that depend on the reaction conditions. At the end of both reactions, most intermediate amyloids disappear and filaments with the same ordered cores as those from human brains remain. Our results provide structural insights into the processes of primary and secondary nucleation of amyloid assembly, with implications for the design of novel therapies.

## Introduction

The assembly of amyloid-β, tau, α-synuclein and TDP-43 into amyloid filaments defines most cases of human neurodegenerative disease ^1^. The hypothesis that the formation of amyloid filaments causes disease is supported by the observation that mutations in the genes that encode these proteins or increase their production give rise to inherited forms of disease ^2^. Moreover, cryo-EM structures of amyloid filaments from human brains have revealed that distinct folds of tau, α-synuclein and TDP-43 define different diseases, suggesting that specific mechanisms of amyloid formation may underlie these diseases ^3–11^. Nevertheless, the molecular mechanisms by which amyloids cause neurodegeneration remain unknown.

It has been suggested that intermediate species, on-pathway to the formation of mature filaments, are main drivers of amyloid toxicity ^12^. Both non-filamentous species, so-called oligomers, and filamentous intermediates, known as protofibrils, have been proposed to play a role. Intermediate species of amyloid assembly are thus an important target for therapeutic intervention. Lecanemab, the first approved drug for Alzheimer’s disease with a measurable reduction of cognitive decline ^13^, is a humanised mouse monoclonal antibody that was raised against what were thought to be protofibrils of synthetic Aβ40 peptide with the Arctic mutation ^14^.

Despite the interest in intermediate species of amyloid formation, little is known about their structures. Due to their transient nature, most experimental data on oligomers and protofibrils comes from *in vitro* assembly reactions with recombinant proteins, including amyloid-β ^15, 16^, tau ^17^ and α-synuclein ^18^. Most *in vitro* reactions yield filaments with ordered cores that are different in structure from human brain filaments. Tau is the only known exception. Amino acids 297-391 (using the numbering of the longest human brain tau isoform) constitute the proteolytically stable core of paired helical filaments (PHFs) from the brains of individuals with Alzheimer’s disease ^19^. The tau(297-391) construct, upon shaking in phosphate buffer with magnesium chloride, forms PHFs with ordered cores that are identical to those from human brains ^7, 20^. The use of sodium chloride instead of magnesium chloride ^20^ leads to the formation of filaments with ordered cores that are identical to those extracted from the brains of individuals with chronic traumatic encephalopathy (CTE) (Falcon et al., 2019).

Here, we used time-resolved cryo-EM to characterise the filamentous intermediates that form during the *in vitro* assembly of tau into PHFs or CTE filaments. We report the formation of a common first intermediate amyloid (FIA) in both reactions, and the presence of multiple, polymorphic filamentous intermediates, with structures that depend on the reaction conditions, at later time points. Our results provide new insights into primary and secondary nucleation of tau amyloid formation that challenge existing theories and provide novel avenues for therapeutic design.

## Results

### Tau monomers adopt extended conformations in solution

We expressed and purified recombinant human tau(297-391) (**Materials and Methods**). Analytical ultracentrifugation indicated that at a concentration of 6 mg/ml, purified tau was monomeric in solution, with flexible conformations (**Extended Data Figure 1a**). Solution-state nuclear magnetic resonance (NMR) confirmed the presence of disordered tau monomers and suggested that amino acids 305-314 and 336-345 have an increased tendency to adopt extended β-strand-like conformations. Similar observations have also been reported for full-length 4R tau ^21^ and for a 4R tau construct comprising amino acids 244-372 (K18) or its 3R version (K19) ^22, 23^. For a more detailed analysis of the dynamic landscape of the conformational ensemble of tau(297-391), we performed Interpretation of Motions by a Projection onto an Array of Correlation Times (IMPACT) analysis of backbone relaxation measurements at different field strengths ^24^. IMPACT analysis indicated that motions in tau(297-391) monomers are best approximated by 5 correlation times ranging from 36ps to 36ns. In particular, 3 regions (amino acids 305-317, 343-349 and 377-381) contribute to slow segmental motion associated with the increased tendency to adopt an extended structure, which was most evident for amino acids 305-317 (**Figure 1; Extended Data Figure 1b-l**).

**Figure 1:**
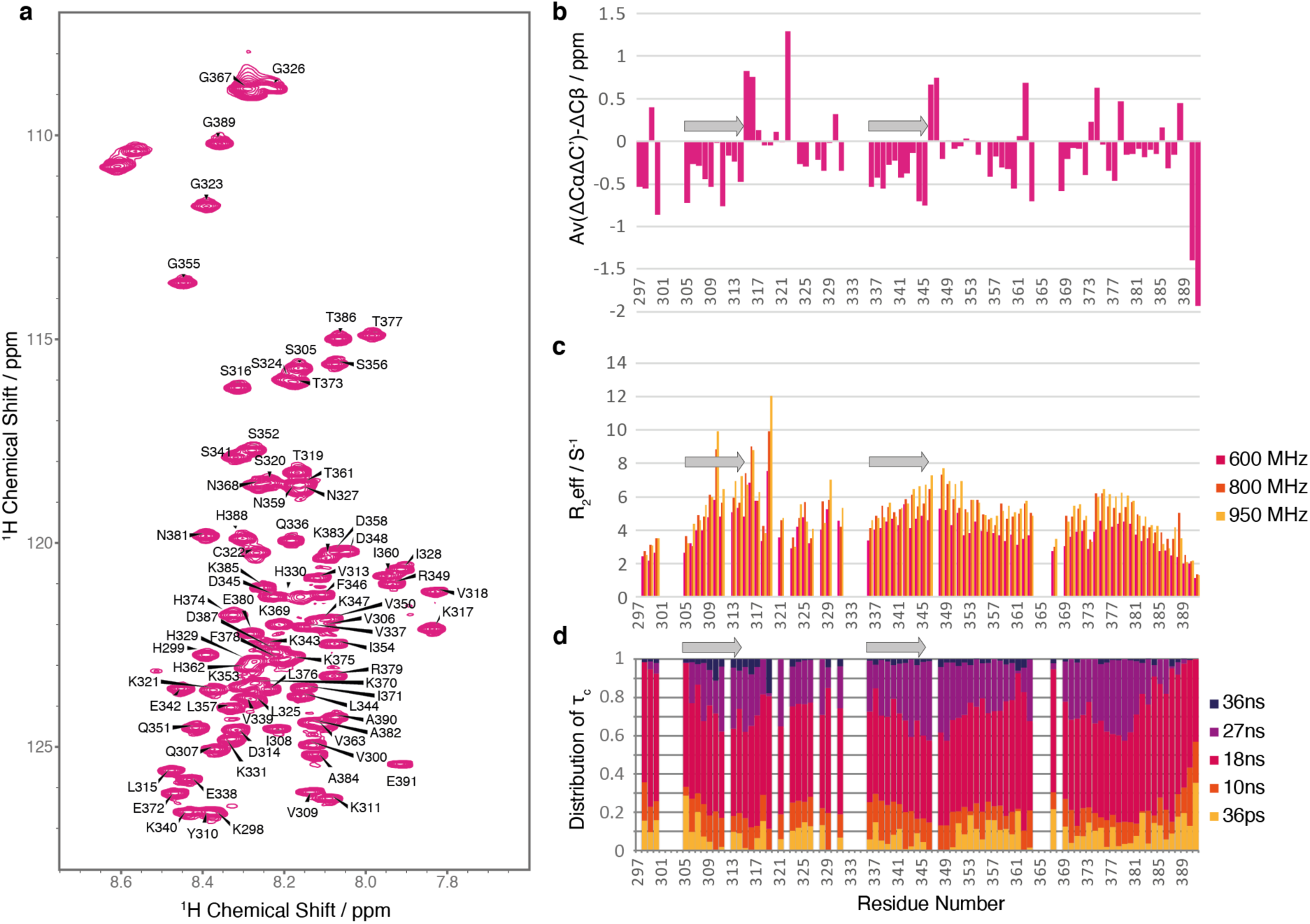
Solution-state NMR on tau monomers. **a.** Assigned 600 MHz ^15^N-^1^H heteronuclear single quantum coherence (HSQC) spectrum of human tau(297-391). **b.** Secondary shift analysis of the backbone Cα, Cβ and C’ chemical shifts. Stretches of amino acids with negative values, as seen for residues 305-314 and 336-345, indicate a propensity to adopt an extended β-strand-like conformation. **c.** Exchange-free transverse relaxation (R2eff) rates collected at 600 (magenta), 800 (red) and 950 MHz (orange). Higher rates are indicative of increased rigidity on the ms-μs timescale. **d.** IMPACT analysis of tau motions on timescales ranging from 36 ps (yellow) to 36 ns (purple). The diagram illustrates the distribution of internal backbone motions (τ) at 5 frequencies. Backbone dynamics of amino acids 305-317, 343-349 and 377-381 display motions at slower frequencies, indicative of segmental motion associated with conformational restrictions. This is most pronounced for amino acids 305-317, with marked contributions from the slowest timescale of motion (36 ns).

### Abundant tau filaments form prior to the increase in ThT fluorescence

We then initiated multiple replicates of 2 assembly reactions. The first reaction was performed in the presence of magnesium chloride for forming PHFs, whereas the second contained sodium chloride for forming CTE filaments. To a subset of reactions, we added 1.5 μM thioflavin T (ThT) to monitor fluorescence continuously. For reactions without ThT, we took aliquots at various time points for cryo-EM structure determination. Because each cryo-EM sample uses 3 μl of the 40 μl reaction, and because not all cryo-EM grids are suitable for data acquisition, we collected cryo-EM data sets from 5 and 6 replicates for each of the PHF and CTE reactions, respectively. Further replicates were used for quantification of pelletable tau by ultracentrifugation and for off-line ThT monitoring, as these required the entire reaction volumes. Multiple different protein preparations were used among the replicates.

For both PHF and CTE reactions, continuous ThT fluorescence monitoring showed a typical sigmoidal curve that has been associated with a nucleation-polymerisation model of amyloid formation (**Figure 2a; Extended Data Figure 2**) ^25^. For approximately the first 240 minutes (min) after starting the assembly reactions, ThT fluorescence remained low, but it increased sharply between 240 and 480 min, after which it plateaued. Off-line ThT measurements were in accordance with continuous monitoring, indicating that the presence of ThT in the reaction mixture did not alter the kinetics.

**Figure 2:**
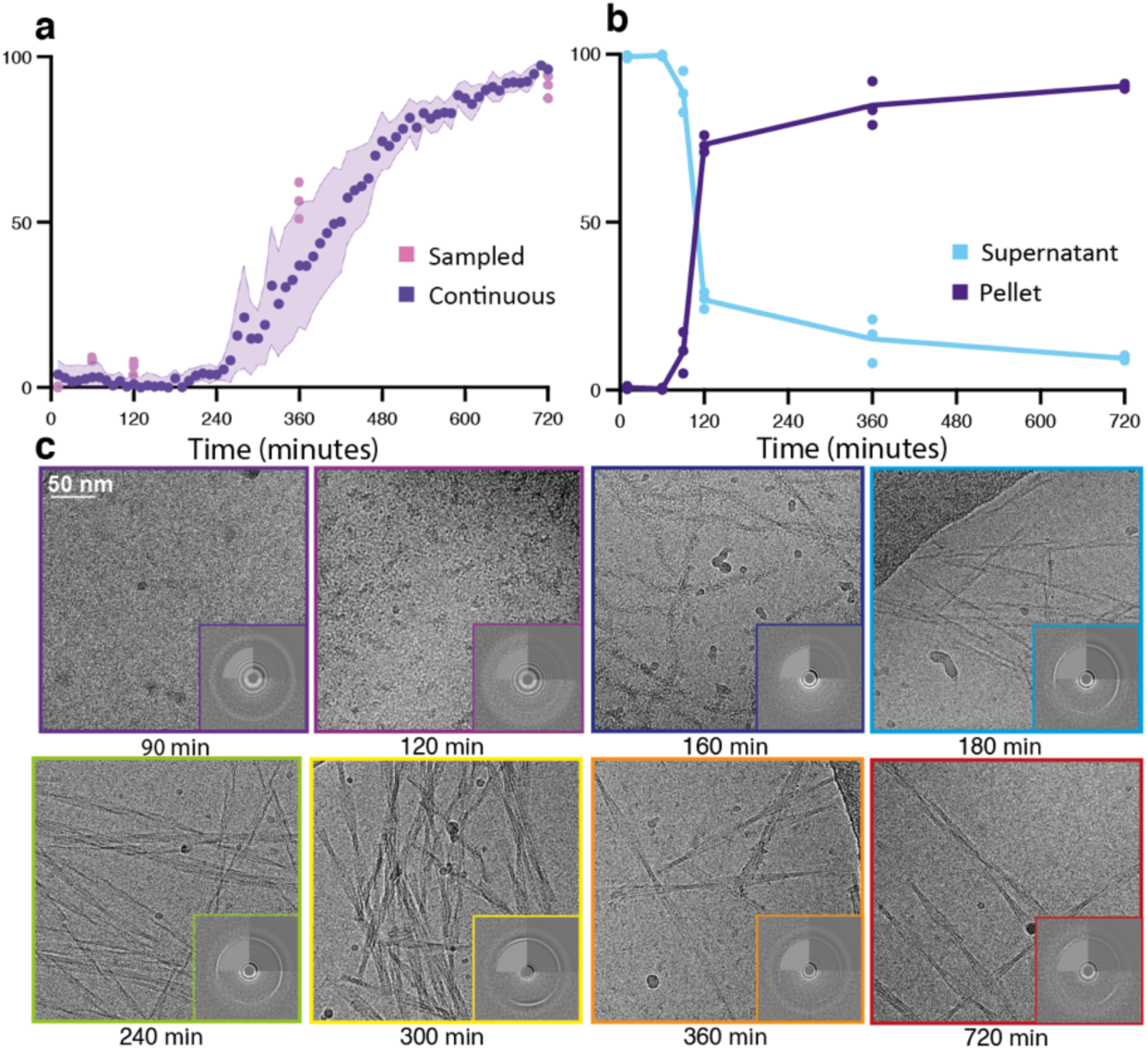
Time-resolved cryo-EM. **a.** ThT fluorescence profile of the PHF reaction. Purple circles indicate the average of 3 replicates; purple shading indicates the standard deviation among replicates; pink circles represent off-line ThT measurements. **b.** The amount of tau in the pellet and in the supernatant (in % of the total amount of tau) after centrifugation for 15 min at 400,000 g, quantified by SDS-PAGE. **c.** Cryo-electron micrographs at various time points in the PHF reaction. Insets show the power spectrum of the electron micrographs, with the water ring at 3.6 Å and/or the 4.75 Å signal that is representative for β-sheet structures.

The samples that were used for off-line ThT fluorescence measurements were also used for quantification of pelletable tau by ultracentrifugation. Because abundant amyloid filaments remained in the supernatants of ultracentrifugation runs at 100,000-130,000 g ^26, 27^, we centrifuged the samples at 400,000 g for 15 min at 20 °C, to quantify the amount of soluble versus pelletable tau by SDS-PAGE (**Supplementary Information**). Until 60 min, almost all tau remained soluble. However, 70-80% of tau was already pelletable at 120 min, and the amount of pelletable tau plateaued at 80-90% at 720 min (**Figure 2b**).

Cryo-EM imaging confirmed the presence of amyloid filaments from 120 min (**Figure 2c**). Images of samples taken at 30, 60 or 90 min were devoid of filaments and did not show evidence of β-sheets (**Extended Data Figure 3a**). However, at 120 min, many filaments were visible in both PHF and CTE reactions, and power spectra showed a strong 4.7 Å signal, indicative of abundant β-sheets. These initial tau filaments have a fuzzy, beads-on-string-like appearance, with a short cross-over distance of 13.5 nm. They range in size from just one or two cross-overs to filaments longer than the field of view (∼300 nm). At later time points, numerous types of amyloid filaments could be distinguished (**Figure 2c**). Using helical reconstruction in RELION ^28^, we solved 163 cryo-EM structures from the PHF and CTE reactions. We built atomic models for 44 different structures with resolutions ranging from 1.7-3.8 Å (**Extended Data Figure 4-5**, **Supplementary Information**).

### The same early filamentous intermediate forms in PHF and CTE reactions

Cryo-EM structure determination revealed the presence of the same filament at 120 min in the PHF and CTE reactions (**Figure 3**). Because we observed no evidence of earlier filaments, and because this filament adopted a cross-β packing characteristic of amyloids, we termed it the first intermediate amyloid (FIA). Although filamentous, the FIA does not generate fluorescence with ThT. The FIA adopts a pseudo-2_1_ helical symmetry and has an atypically large, left-handed twist of -6.3°.

**Figure 3:**
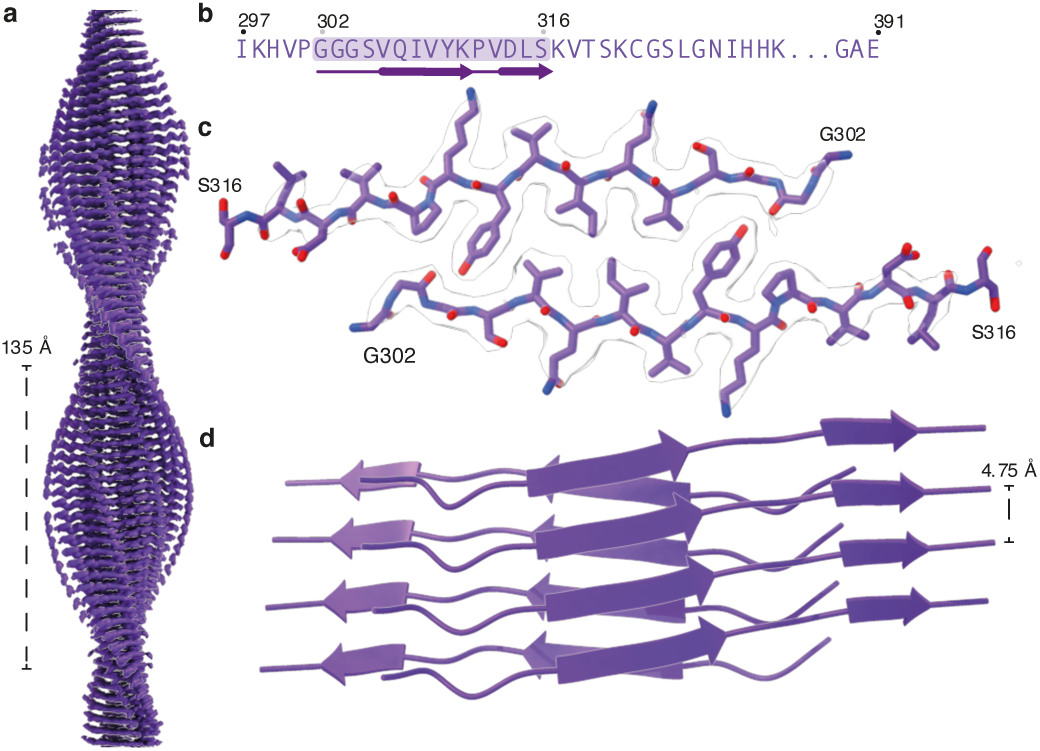
Structure of the FIA. **a.** Side view of the cryo-EM reconstruction, with the cross-over distance indicated. **b.** Amino acid sequence of the ordered core (highlighted in purple). **c.** Top view of the cryo-EM density (in transparent white) and the atomic model. **d.** Side view of the atomic model in cartoon representation.

The ordered core of the FIA is the smallest of any amyloid observed by cryo-EM helical reconstruction to date, comprising amino acids _302_GGGSVQIVYKPVDLS_316_ from two anti-parallel tau molecules, with a predominantly hydrophobic interface. At its centre, the side chains of valine 306 and isoleucine 308 from opposite protofilaments pack against each other and are flanked by the side chain of tyrosine 310, the hydroxyl groups of which form hydrogen bonds to the backbone groups of glycine 303 and serine 305. The protofilament interface is structurally similar to the β-sandwich that was first observed between parallel β-sheets in globular proteins with a β-helix fold ^29^. Also, the twist in the opposite β-sheets of these β-helices is similar to that in the β-sheets of the FIA. In other amyloid filaments, including PHFs and tau filaments of CTE, the twist in β-sheets is typically much smaller. The twisted β-sheets of the FIA also represent a major difference with crystal structures of the _306_VQIVYK_311_ peptide alone ^30, 31^, where β-sheets in the crystal packed against each other without twist (**Extended Data Figure 6**).

The FIA only exists for a short time. At 120 min, 100% of the filaments that yield interpretable 2D class averages are FIAs, but they are no longer observed at 160 min (**Extended Data Figure 3a**). At 140 and 160 min in the PHF reaction, multiple different types of filaments give rise to uninterpretable 2D class averages, many of which lack the helical twist required to solve their structures (**Extended Data Figure 3b)**. At 160 min in the CTE reaction, we were able to solve 9 structures (**Extended Data Figure 3c**).

### Later filamentous intermediates in the PHF reaction are polymorphic

From 180 min, we observed multiple types of filaments in the PHF reaction (**Figure 4a**). Most filaments at 180 min were made of two identical protofilaments with an ordered core that comprised amino acids 305-380, similar to the extent of the ordered core of PHFs ^7^. As is the case of the Alzheimer fold, these protofilaments formed a turn of a β-helix at amino acids 337-356. However, whereas the Alzheimer fold is C-shaped, the ordered cores of most protofilaments at 180 min adopted a more elongated, J-shaped conformation. In the different filament types, the J-shaped protofilaments packed against each other in different ways. During the next 3 h, additional types of filaments formed. In total, we solved 24 different structures from samples taken at 120, 180, 240, 300, 360 and 720 min, with 20 structures to resolutions sufficient for atomic modelling (**Extended Data Figure 4**). Again, most filaments comprised two protofilaments that packed against each other in various ways. Some filaments with 3 or 4 protofilaments also formed, including the previously described triple and quadruple helical filaments (THFs and QHFs) ^20^.

**Figure 4.**
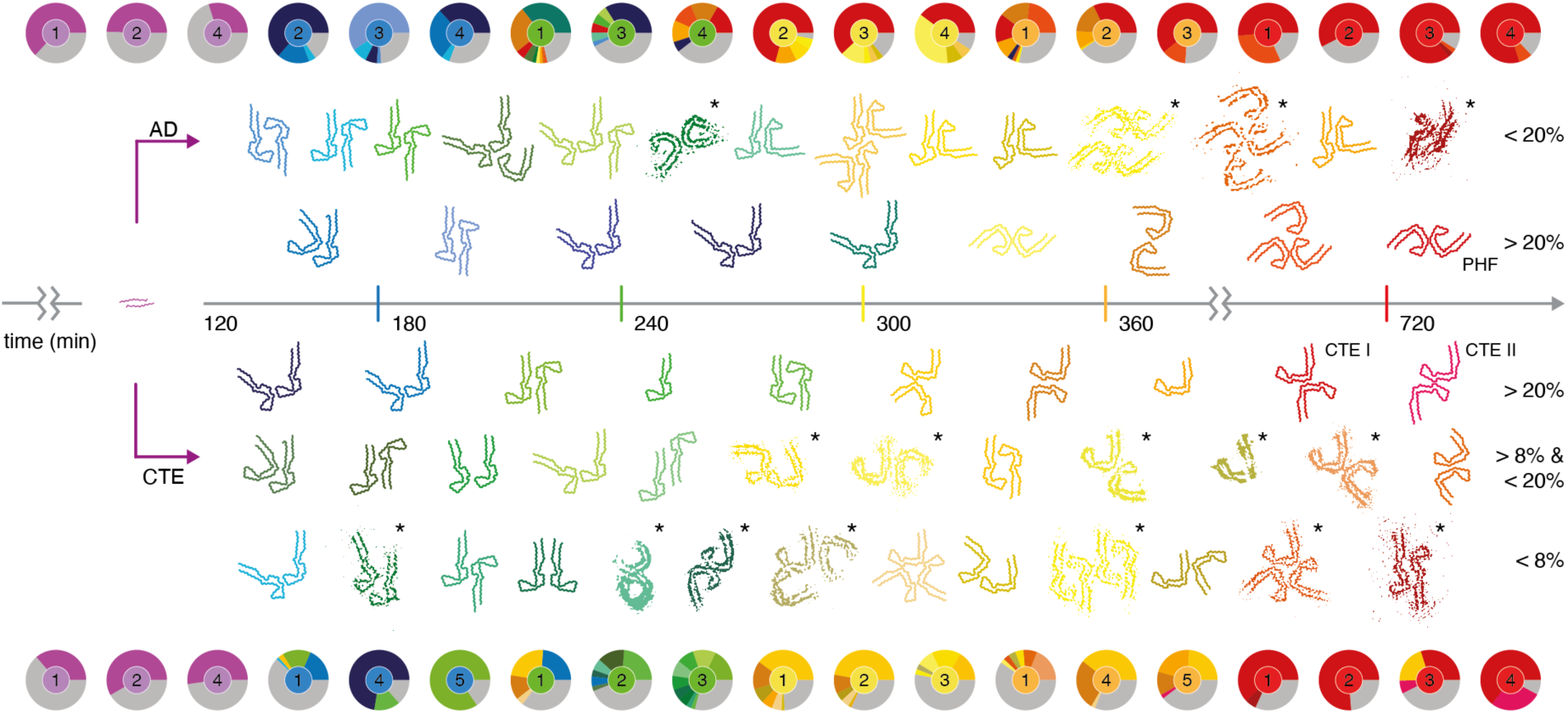
Overview of structures in the assembly reactions. Pie charts show the relative abundance of the structures determined for each replicate (1-5) of the PHF reaction (top) and the CTE reaction (bottom). Main chain traces for atomic structures are shown in the same colours as the pie chart segments for each reaction. Grey segments represent filaments for which no structures were solved. Structures that were solved at resolutions insufficient for atomic modelling are shown as thresholded densities and are indicated with asterisks. Structures and pie chart central circles are coloured per time point (120 min in purple; 180 min in blue; 240 min in green; 300 min in yellow; 360 min in orange and 720 min in red). Structures are coloured according to the time point at which they are most abundant, averaged across all replicates. More abundant structures (assessed by maximal percentage across all replicates) are closer to the time axis, whereas less abundant ones are further away. Details of all data sets and structures, including pie charts of additional replicates, are shown in the **Supplementary Information**.

As time progressed, filaments with two J-shaped protofilaments disappeared and filaments with two C-shaped protofilaments appeared (**Figure 5a**). Between 240 and 360 min, filaments with one J-shaped and one C-shaped protofilament were also present. The inter-protofilament packing of the latter resembled the asymmetrical arrangement of protofilaments in the straight filaments extracted from the brains of individuals with Alzheimer’s disease. Among J-shaped and C-shaped protofilaments, the opposing β-strands comprising amino acids 305-320 and 365-380 hardly changed their conformation, confining all differences to the β-helix turn and its surrounding amino acids. Two types of J-shaped protofilaments exist. Earlier J-shaped protofilaments tend to be straighter, whereas the β-helix turn in later J-shaped protofilaments turns inwards, towards the rest of the protofilament. This change in orientation of the β-helix turn is reflected in a distinct conformation of the _332_PGGG_335_ motif. The difference between the later J-shaped protofilament and the earliest C-shaped protofilaments coincided with a re-arrangement of the _364_PGGG_367_ motif on the opposite side of the protofilament. Finally, the change from earlier C-shaped protofilaments to the final, more closed, C-shaped protofilaments of PHFs involves a second inwards rotation of the β-helix turn, which again concurs with a re-arrangement of the _332_PGGG_335_ motif.

**Figure 5:**
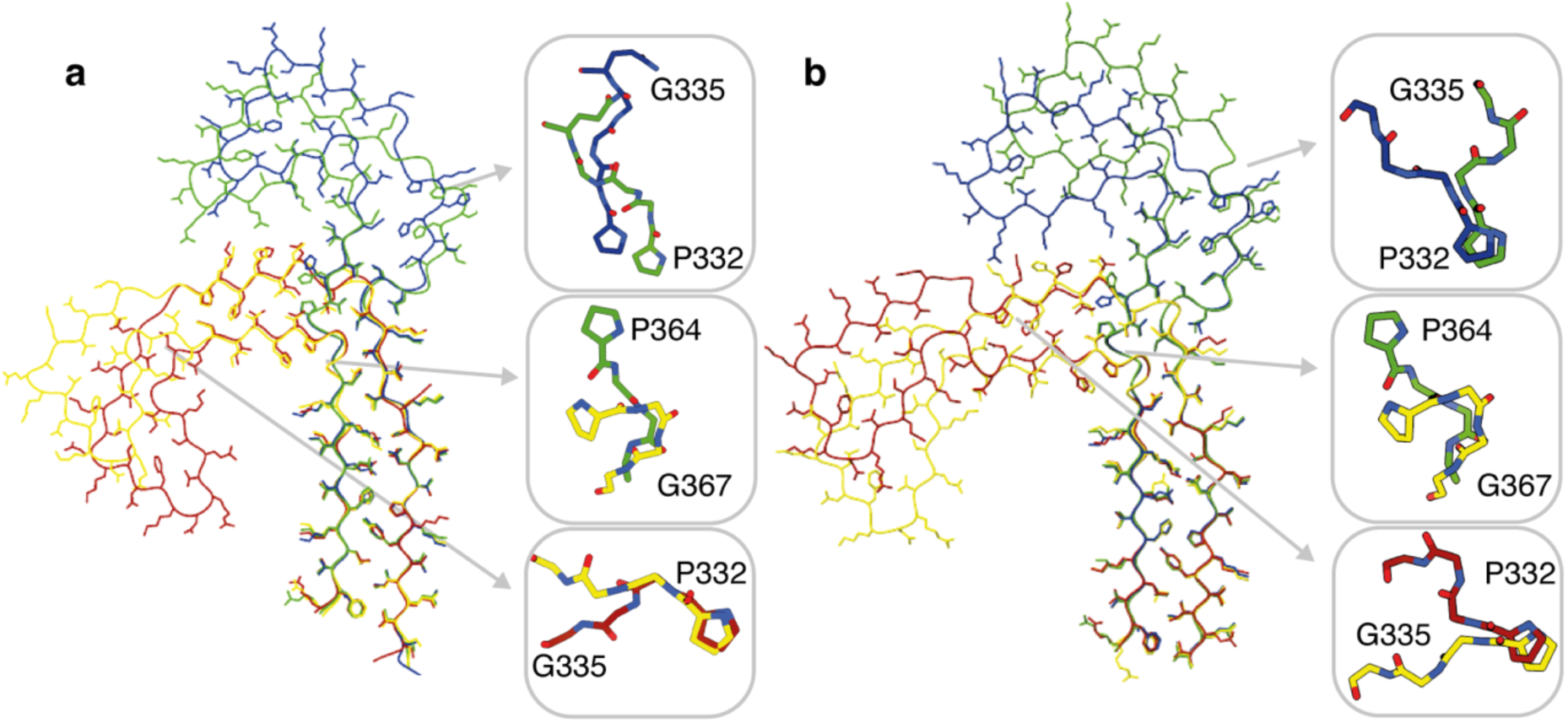
Protofilament maturation. **a.** Atomic models for protofilaments in the PHF reaction at 180 min (blue), 240 min (green), 300 min (yellow) and 720 min (red). Insets show the corresponding conformations of the 322PGGG335 and 364PGGG367 motifs. **b.** As in a, but for the CTE reaction.

Some filaments that resembled PHFs were already present at 240 min. Although they had the same double C-shaped protofilament arrangement as in PHFs, their cross-over distances tended to be more variable at earlier time points. At later time points, most filaments had cross-over distances of 750-900 Å, similar to those of PHFs extracted from the brains of individuals with Alzheimer disease ^7^. The amino- and carboxy-terminal ends of each protofilament packed against each other within the same β-rung. However, at earlier time points, filaments with cross-over distances as large as 2,900 Å formed, in which amino acids at the amino-terminus of the protofilament packed against amino acids at the carboxy-terminus that were one or more β-rungs lower. In addition, the position along the helical axis of the β-helix turn compared to the amino- and carboxy-termini of the protofilament also changed as the cross-over distances decreased. These conformational changes correlated with peptide flips at glutamic acid 342 and isoleucine 354 (**Extended Data Figure 7**).

Finally, by 720 min, most filaments had adopted the same ordered core as that of PHFs extracted from the brains of individuals with Alzheimer’s disease, although THFs remained in some replicates. Overall, the 5 replicates were relatively consistent in their timing.

### Large numbers of later filamentous intermediates form in the CTE reaction

In the CTE reaction, a greater number of intermediate structures formed than in the PHF reaction (**Figure 4**). In total, we determined the structures of 39 different filament types, with 24 structures being of sufficient resolution for atomic modelling (**Extended Data Figure 5**). As in the PHF reactions, most intermediate filament types consisted of two protofilaments with ordered cores that comprised residues 305-380, and the protofilaments packed against each other in multiple ways. Most filament types also adopted a J-shaped conformation at earlier time points and, as time progressed, more C-shaped protofilaments appeared. No filaments with one J-shaped and one C-shaped protofilament were observed. As for the intermediates in the PHF reaction, the _332_PGGG_335_ and _364_PGGG_367_ motifs appeared to play a central role in the maturation of J-shaped to C-shaped protofilaments (**Figure 5b**).

The presence of sodium chloride in the CTE reaction affected the conformation of the β-helix turn in all intermediate amyloids and the final CTE structures, which showed a more open β-helix turn than in the Alzheimer fold, together with the presence of an extra density inside the β-helix turn. This extra density was previously interpreted as sodium chloride ion pairs ^20^. In the PHF reaction, some earlier intermediate filaments also showed a similar extra density. It is likely that traces of sodium chloride, which was used during purification of recombinant tau(297-391), were still present in the PHF reaction.

Most intermediates that formed in the CTE reaction comprised two identical protofilaments; some filaments made of either 1 protofilament or 3 protofilaments were also present. Compared to the filaments in the PHF reaction, intermediates in the CTE reaction displayed a greater variation in inter-protofilament packing. Many packings seemed to be coordinated by electrostatic interactions (**Extended Data Figure 8**). Relatively small differences in the protofilament packing of individual pairs of filament types suggest that intermediate amyloid filaments may mature through subsequent sliding of their protofilaments relative to each other.

After 720 min, all reactions contained CTE type I filaments ^5^. In replicates 2, 3, 4 and 5, CTE type II filaments were also present. The different replicates of the CTE reaction were reasonably well synchronised, except for replicate 3, which ran a bit slower.

## Discussion

Polymorphism is a common phenomenon in crystallography. Ostwald’s interpretation of crystal polymorphism explains how the state that nucleates is not necessarily the most thermodynamically stable. Instead, the state that most closely resembles the solution state is kinetically advantaged ^32^. This interpretation may also be relevant for understanding the assembly of tau into amyloid filaments. Being the product of a long disease process, tau PHFs and CTE filaments likely represent a thermodynamically stable state. *In vitro* assembly of recombinant tau(297-391) converges onto the same structures over 12 h, but only after multiple polymorphic intermediate amyloids have formed and disappeared again.

In the first intermediate, the FIA, only 15 amino acids of each tau molecule are ordered; the remaining 80 amino acids are not resolved in the cryo-EM map, suggesting that they remain largely unstructured. Thereby, for 84% of the amino acids in the FIA, the first detectable nucleated state probably closely resembles the solution state. Our solution-state NMR data suggest that some of the 15 amino acids of the FIA’s ordered core may already adopt extended, β-strand-like conformations in monomeric tau, with slower dynamics than the rest of the protein, which will reduce further the differences between solution and nucleated states. The fact that β-sheets in the FIA are more twisted than in other amyloids may also play a role. The ordered core of the FIA explains the previously observed importance of the _306_VQIVYK_311_ (PHF6) motif for the assembly of full-length human tau into filaments *in vitro* ^33^ and in transgenic mice ^34^. The existence of filamentous intermediates in amyloid assembly reactions has been reported previously, for example using atomic force microscopy of amylin ^35^. It remains to be seen if primary nucleation of other proteins for which small amyloid-prone regions have been identified, such as acylphosphatase, amyloid-β, α-synuclein, TDP-43, transthyretin and the prion protein ^36–39^, occurs through similar FIA structures.

The sudden assembly of 70-80% of tau molecules into the FIA at 120 min, after a 90-min lag period, together with the complete disappearance of the FIA after 180 min, suggest that primary nucleation of the FIA is rate-limiting and that, once formed, the FIA can seed the rapid growth of other filament types through secondary nucleation mechanisms. Based on kinetic studies ^40, 41^, secondary nucleation has been proposed to play a central role in amyloid assembly, but molecular insights into how such nucleation processes happen have been scarce. Our cryo-EM structures provide a basis for new hypotheses on how such secondary nucleation processes may occur (**Figure 6**).

**Figure 6:**
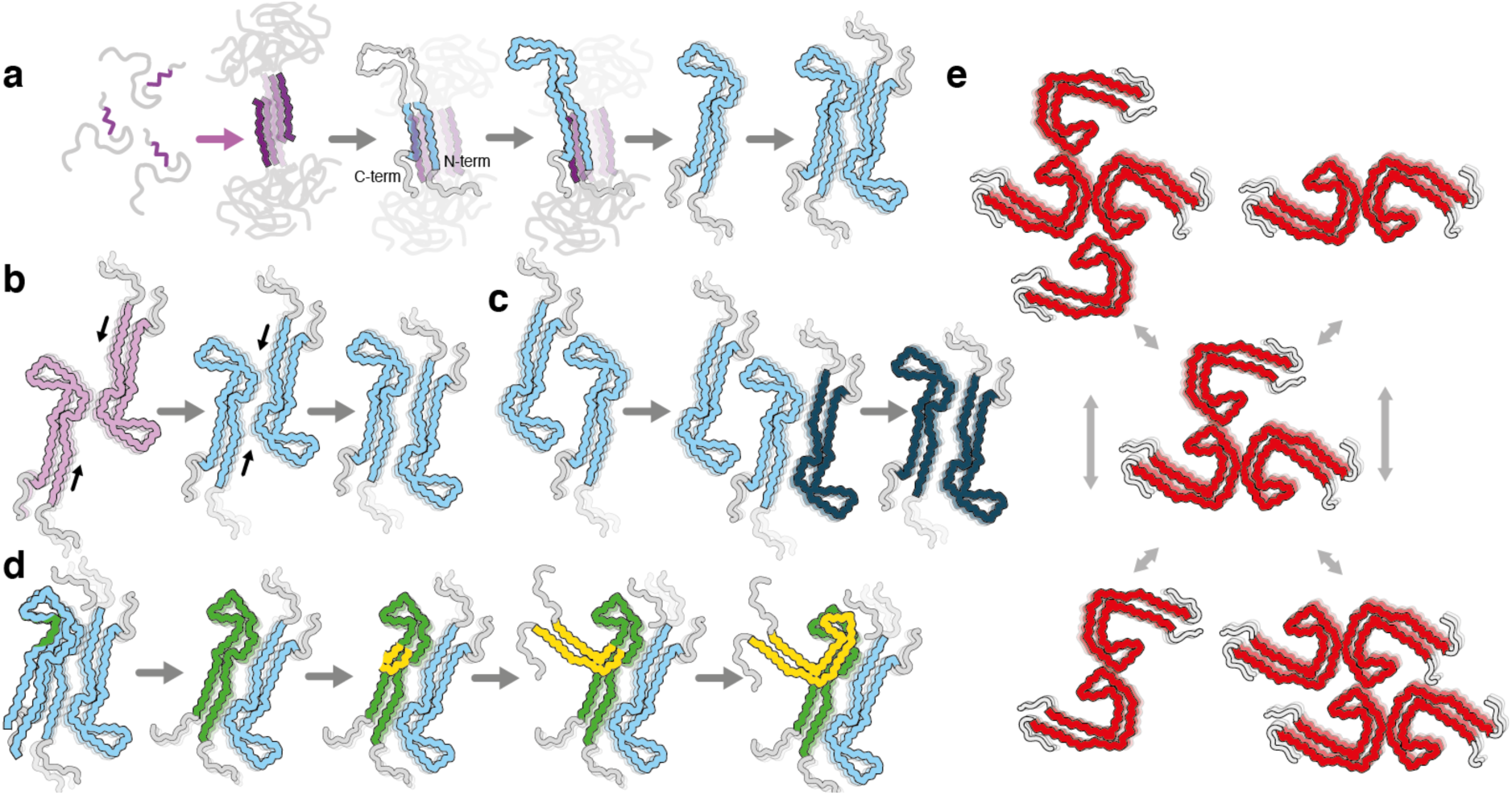
Models for primary and secondary nucleation. **a.** Primary nucleation (purple arrow) of disordered monomers with partially rigid β-strands may lead to formation of the FIA. Subsequent secondary nucleation (grey arrows) may then occur through folding back of the carboxy-terminal domain of tau monomers at the end of, or at defects in, the FIA to form the interface between amino acids 305-316 and 370-380 that remains nearly constant in all intermediate and final filament types. This folding back may then lead to the seed of an early J-shaped protofilament. Possibly, the formation of singlets of J-shaped protofilaments (as observed in the CTE reaction) is followed by packing of a second protofilament to form more stable doublets. **b.** The sliding of protofilaments relative to each other, again at the ends of, or at defects in, filaments may lead to the subsequent formation of more stable structures. **c.** Alternatively, more stable protofilament interactions may form at the sides of existing filaments. **d.** Protofilament maturation, from early J-shaped protofilaments to later C-shaped protofilaments may also happen at the end of, or at defects in, filaments. **e.** Growth, association or dissociation of protofilaments at the sides of filaments may lead to the conversion of multiple filament types.

While amino acids 302-316 from both protofilaments of the FIA pack in a homotypic, anti-parallel manner, amino acids 305-316 pack in a heterotypic arrangement against amino acids 370-380 in later filament types. Therefore, major topological re-arrangements need to occur in the conversion of the FIAs into any of the later intermediates. The observation that amino acids 305-316 and 370-380 adopt almost the same conformation in all later intermediates and the final structures suggests that this part of the protofilament forms early and is relatively stable. Possibly, amino acids 370-380 from one tau molecule in the FIA fold back and pack against amino acids 305-316 of the same molecule, instead of forming the homotypic packing with another tau molecule in the opposite protofilament. This could happen at either end of the FIAs and at defects along their length. Once the seed for a J-shaped protofilament forms in this way, it may then lead to the formation of various filament types with two J-shaped protofilaments (**Figure 6a**). Other models of secondary nucleation include: maturation of filament types through the sliding of protofilaments relative to each other (**Figure 6b**), growth, association and/or dissociation of protofilaments at the sides of filaments (**Figure 6c & 6e**) and formation of new filament types by partial structural templating at the end of, or at defects in, existing protofilaments (**Figure 6d**). It is possible that any of these mechanisms also lead to the conversion of one filament type into another. Filaments with variations in structure along their length have been observed previously, for example for heparin-induced filaments of recombinant tau ^42^, for TDP-43 filaments from human brain ^3^, and for filaments of immunoglobulin light chains from human heart ^43^. In our images, most filaments are not branched and do not appear to consist of multiple filament types, although we do observe some exceptions (**Extended Data Figure 3d-g**).

These cryo-EM snapshots do not provide insights into the kinetics of the assembly reactions. The FIA, the later intermediates and the PHF and CTE filaments are probably in exchange with monomers from the solution, leading to their continuous elongation and disassembly. Future kinetic studies are required to explore in detail the interplay between species. Our results indicate that ThT fluorescence is suboptimal for the quantification of filament formation, since many intermediate amyloids do not give rise to ThT fluorescence. ThT-negative amyloid filaments have been reported previously, e.g. for TDP-43 ^44^, as well as for transthyretin, β_2_-microglobulin, and some forms of amyloid-β ^45^. The factors that determine whether an amyloid filament is detected by ThT fluorescence remain unclear.

In human brains, tau pathology is thought to spread through templated seeding, akin to the spread of prions, where small amounts of filaments accelerate the assembly of newly formed filaments with the same structures as those of the seeds ^46^. In disease, the initial seeds may form through the same short-lived intermediates as described here, but it will be difficult to prove their presence in human brains. The molecular mechanisms of templated seeding remain poorly understood. On the one hand, growth of filaments at their ends alone does not explain the kinetics of seeded aggregation *in vitro* ^47^. On the other hand, it is not clear how structural templating happens in alternative models, where secondary nucleation has been proposed to happen on the sides of filaments ^41^. Seeded assembly *in vitro* does not necessarily reproduce the structure of the filaments that are used as seeds ^48^. Although structural templating may happen more readily in cultured cells, even there the biochemical environment seems to affect which structures form ^49^. Moreover, observations that even tau monomers ^50, 51^ can seed tau aggregation in cells, are difficult to understand with the existing models of seeded aggregation. Akin to the experiments described here, time-resolved cryo-EM of *in vitro* seeded assembly reactions may provide a better understanding of the molecular mechanisms of templated seeding.

Distinct structures of tau filaments extracted from the brains of individuals with different diseases have led to a structure-based classification of tauopathies ^9^. Extra densities in the cryo-EM maps suggest that unidentified molecules may co-assemble with tau. The biochemical environment in which filaments are formed may thus play a defining role in the formation of specific structures in the different diseases. Our *in vitro* assembly reactions recapitulate this for Alzheimer’s disease and CTE. The presence of magnesium chloride or sodium chloride in the buffer defines the structure of the final reaction products, as well as those of intermediates that form after 180 min. However, independently of the reaction conditions, the same FIA forms after 120 min. If the FIA can form in a wide range of biochemical environments, then the same primary nucleation event is likely to lead to the formation of filaments from other tauopathies. Identification of the corresponding reaction conditions will be required to test this hypothesis.

Our experiments reveal the structures of intermediate amyloid species that form during the *in vitro* assembly of tau(297-391) into filaments with the Alzheimer or CTE fold. Whether these short-lived species can be isolated as biochemically stable entities against which antibodies could be raised remains to be tested. Because small oligomeric species would be difficult to visualise using cryo-EM, we cannot exclude their presence in our experiments. Nevertheless, our data suggest that the *in vitro* formation of disease-specific tau folds happens through a mechanism whereby tau monomers nucleate directly into FIAs, which then grow and turn into mature filaments through multiple routes of secondary nucleation, involving many different intermediate amyloids. This model, in which prefibrillar oligomeric species are not required, provides a new perspective on the molecular mechanisms of amyloid formation, with important implications for the development of new therapies.

## Acknowledgements

We thank Cyril Charlier and Fabien Ferrage for providing the Mathematica script for IMPACT analysis; Harrison Wang and Tuomas Knowles for helpful discussions; Jake Grimmett, Toby Darling and Ivan Clayson for help with high-performance computing; and Max Wilkinson and Tony Crowther for critical reading of the manuscript. This work was supported by the facilities for biophysics, electron microscopy, NMR and scientific computing of the Medical Research Council (MRC) Laboratory of Molecular Biology, and by the Francis Crick Institute through provision of access to the MRC Biomedical NMR Centre. The Francis Crick Institute receives its core funding from Cancer Research UK (CC1078), the UK Medical Research Council (CC1078), and the Wellcome Trust (CC1078). This work was supported by the MRC, as part of United Kingdom Research and Innovation (UKRI) [MC_U105184291 to M.G. and MC_UP_A025-1013 to S.H.W.S.], and a Marshall scholarship to D.L.

## Author contributions

S.L. performed biochemistry and filament assembly; S.L and A.K. acquired cryo-EM data; S.L., D.L., D.K., A.M. and S.H.W.S. analysed cryo-EM data; D.L. and D.K. wrote software; J.L.W. and S.M.V.F. performed NMR; S.H.M. performed AUC; M.G. and S.H.W.S. supervised the project. All authors contributed to the writing of the manuscript.

## Additional Information

For the purpose of open access, the MRC Laboratory of Molecular Biology has applied a CC BY public copyright licence to any Author Accepted Manuscript version arising.

## Data Availability

Cryo-EM maps have been deposited in the Electron Microscopy Data Bank (EMDB) under accession numbers XXXX. Refined atomic models have been deposited in the Protein Data Bank (PDB) under accession number XXXX.

## Methods

### Expression and purification of tau(297-391)

Expression of tau(297-391) was carried out in *E. coli* BL21 (DE3)-gold cells (Agilent Technologies, USA), as described ^52^. In brief, one plate of cells was resuspended in 1 litre 2xTY (Tryptone Yeast) supplemented with 100 mg/l ampicillin and grown to an OD_600_ of 0.8. For uniformly ^15^N and ^13^C labelled tau, bacteria were grown in isotope-enriched M9 minimal media containing 1 g/l of ^15^N-ammonium chloride and 2 g/l of ^13^C-glucose (Sigma, USA) supplemented with 1.7 g/l YNB (Yeast Nitrogen Base) (Sigma, USA). Cells were induced by the addition of 1 mM IPTG for 4 h at 37 °C, harvested by centrifugation (4,000 g for 20 min at 4 °C), resuspended in washing buffer (WB: 50 mM MES at pH 6.0; 10 mM EDTA; 10 mM DTT, supplemented with 0.1 mM PMSF and cOmplete EDTA-free protease cocktail inhibitors, at 10 ml/g of pellet) and heated at 95 °C for 5 min. Cell lysis was performed using sonication (at 40% amplitude using a Sonics VCX-750 Vibracell Ultra Sonic Processor for 7 min, 5 s on / 10 s off). Lysed cells were centrifuged at 20,000 g for 35 min at 4 °C, filtered through 0.45 μm cut-off filters and loaded onto a HiTrap CaptoS 5 ml column (GE Healthcare, USA) for cation exchange. The column was washed with 10 column vol of WB and eluted using a gradient of WB containing 0–1 M NaCl. Fractions of 3.5 ml were collected and analysed by SDS-PAGE (Tris-Glycine 4–20% gels). Protein-containing fractions were pooled and precipitated using 0.28 g/ml ammonium sulphate and left on a rocker for 30 min at 4 °C. Precipitated proteins were then centrifuged at 20,000 g for 35 min at 4 °C, and resuspended in 2 ml of 10 mM phosphate buffer at pH 7.2 with 10 mM DTT, and loaded onto a 16/600 75 pg column size exclusion column. Size exclusion fractions were analysed by SDS-PAGE, protein-containing fractions were pooled and concentrated to 20 mg/ml using molecular weight concentrators with a cut-off filter of 3 kDa. Purified protein samples were flash frozen in 50-100 μl aliquots for future use. Protein concentrations were determined using a NanoDrop2000 (Thermo Fisher Scientific, USA).

### Analytical ultracentrifugation

Tau at a concentration of 6 mg/ml in 10 mM sodium phosphate, pH 7.2, 10 mM DTT was loaded into 12 mm 2-sector cells, placed in an An50Ti rotor and centrifugated 50,000 rpm at 4 °C using an Optima XL-I analytical ultracentrifuge (Beckman). The data were analysed in SEDFIT-16.1c ^53^ using a c(s) distribution model. The partial-specific volumes (v-bar), density and viscosity of the buffer were calculated using Sednterp ^54^. Data were plotted with the program GUSSI 1.4.2 ^55^ and Prism 9.5.1 (GraphPad Software).

### Nuclear magnetic resonance (NMR)

Solution-state NMR data were acquired at 278 K using 14.1 Tesla (T), 18.8 T and 22.3 T Bruker Avance III spectrometers fitted with 5 mm TCI triple resonance cryoprobes, operating at proton frequencies of 600, 800 and 950 MHz respectively. All NMR samples were prepared in 50 mM phosphate buffer at pH 7.4, with 150 mM NaCl, 10 mM DTT, and with 5% D_2_O as a lock solvent.

Assignment of backbone NH, N, Cα, Cβ and C’ resonances of isotopically enriched (^15^N/^13^C) human tau(297-391) at 300 μM was completed at 600 MHz. Standard 3D datasets were acquired as pairs to provide own and preceding carbon connectivities, using 18-39% non-uniform sampling (NUS) to aid faster data acquisition. Both the HNCO & HN(CA)CO, and HNCA & HN(CO)CA experimental pairs were collected with 2048, 64 and 128 complex points in the ^1^H, ^15^N and ^13^C dimensions respectively. The CBCA(CO)NH and HNCACB pair were collected with 2048, 64 and 96 complex points in the ^1^H, ^15^N and ^13^C dimensions respectively. Additional ^15^N connectivities were established using (H)N(COCA)NNH experiments with 2048, 80 and 128 complex points in the ^1^H, direct and indirect ^15^N dimensions, respectively. C’-detect experiments c_hcacon_ia3d and c_hcanco_ia3d were also collected to aid backbone assignment with 1024, 64 and 128 points collected in the direct ^13^C, ^15^N and indirect ^13^C dimensions, respectively.

All raw NMR data was processed using Topspin versions 3.2 or 4 (Bruker), or using NMRPipe ^56^, with compressed sensing for reconstruction of non-uniform sampling data ^57^ and analysed using NMRFAM-Sparky and MARS ^58^.

Secondary chemical shift analysis was performed to probe secondary structure elements. Random coil Cα, Cβ and C’ values for tau(297-391) were calculated ^59–61^, and subtracted from the experimentally derived values, to give ΔCα and ΔCβ and ΔC’, respectively. Negative values for ΔCβ – (ΔCα + ΔC’)/2 are indicative of amino acids in an extended backbone conformation; positive values indicate helical conformations.

Molecular motions were probed at proton frequencies of 600, 800 and 950 MHz (^15^N frequencies of 60.8, 80.8 and 96.3 MHz, respectively). Longitudinal relaxation was probed with a standard Bruker ^15^N T1 pseudo-3D experiment, collected with 10, 20, 40, 80, 120, 160, 320, 640, 1280, and 2000 ms delays and a standard Bruker ^15^N T2 pseudo-3D experiment with 16.9, 33.8, 50.7, 67.6, 84.5, 101.4, 118.3, 169, 202.8 and 253.5 ms delays. Both experiments used a recovery delay of 5 s. Picosecond motions were monitored with the standard Bruker interleaved 2D ^15^N{^1^H} heteronuclear nuclear Overhauser effect (NOE) experiment with a recovery delay of 5 s. Additional longitudinal and transverse cross-correlated cross-relaxation (CCCR) measurements were collected as described ^62^, with ΔT relaxation periods of 100 and 40 ms, respectively. Calculation of the exchange-free R2 rates (R2eff) was performed as described ^63^.

IMPACT analysis was completed using a Mathematica (Wolfram, Oxford UK) script, as described ^24^. Corresponding longitudinal and transverse CCCR measurements and ^15^N{^1^H} heteronuclear NOE values collected at three different field strengths were used to create a spectral density analysis landscape. These frequency-specific data were fitted to multiple correlation times, varying in number from 4 to 9, and across a range of correlation times (from 2ps-2ns up to 100ps-100ns) using Monte Carlo simulations. Akaike’s information criterion was used to evaluate the statistical relevance of each condition, indicating that the best representation comprised 5 times scales, ranging from 36 ps to 36 ns.

### Assembly of tau

Filaments were assembled as described ^48^, with minor modifications. Assembly reactions were performed in aliquots of 40 μl of purified monomeric tau(297-391) at 6 mg/ml, in a 384-well microplate that was sealed with a black seal and a plastic seal and placed in a Fluostar Omega (BMG Labtech, Aylesbury, United Kingdom). Reactions were performed for 720 min using 200 rpm orbital shaking at 37 °C. AD reactions were performed in 10 mM phosphate buffer at pH 7.2, 100 mM MgCl_2_ and 10 mM DTT. CTE reactions were performed in 50 mM phosphate buffer at pH 7.2, 150 mM NaCl and 10 mM DTT. For continuous ThT monitoring, 1.5 μM of ThT was added to the reaction and measurements were taken every 10 min. For off-line monitoring, multiple replicates of the reactions were performed, and for each time point ThT was added to the entire volume of a separate reaction replicate at a concentration of 1.5 μM.

### Quantification of pelletable tau

Multiple replicas of the reactions were also performed for the quantification of pelletable tau. At 0, 60, 90, 120, 360 and 720 min, the entire volume of individual reaction replicas was collected for ultracentrifugation. Reactions were centrifuged at 400,000 g at 20 °C for 15 min in polycarbonate centrifuge tubes (Beckman Coulter Inc. USA). The pellets were resuspended in 40 μl reaction buffer, to match the volume of the supernatants. Loading buffer was added to supernatants and pellets, which were then heated for 5 min at 95 °C, and 1.5 μl of each were run by SDS-PAGE (4-20% Tris-Glycine gels). Band intensities were quantified using ImageJ and data were plotted using Prism 9.5.1 (GraphPad Software).

### Electron cryo-microscopy

At specific time points, the microplates were taken from the shaker and 3 μl of the reaction mixture were applied to glow-discharged R1.2/1.3, 300 mesh carbon Au grids. The grids were plunge-frozen in liquid ethane using a Vitrobot Mark IV (Thermo Fisher Scientific). After taking each aliquot, the microplate was re-sealed and returned to the shaker to continue the assembly reaction within 10 min.

Cryo-EM data were acquired at the MRC Laboratory of Molecular Biology (LMB) and at the Research and Development facility of Thermo Fisher Scientific in Eindhoven (TFS). At LMB, images were recorded on a Krios G2 (Thermo Fisher Scientific) electron microscope that was equipped with a Falcon-4 camera (Thermo Fisher Scientific) without an energy filter. At TFS, images were recorded on a Krios G4 (Thermo Fisher Scientific) with a cold field-emission gun, a Falcon-4 camera and a Selectris X (Thermo Fisher Scientific) energy filter that was used with a slit width of 10 eV. All images were recorded at a dose of 30–40 electrons per Å^2^, using EPU software (Thermo Fisher Scientific), and converted to tiff format using relion_convert_to_tiff ^64^ prior to processing (**Supplementary Information**).

### Cryo-EM data processing

Movie frames were gain corrected, aligned, and dose weighted using RELION’s motion correction program ^65^. Contrast transfer function (CTF) parameters were estimated using CTFFIND-4.1 ^66^. Helical reconstructions were carried out using RELION-4.0 ^28, 67^. Filaments were picked manually or automatically using a modified version of Topaz ^68, 69^. Picked particles were extracted in boxes of either 1024 or 768 pixels and down-scaled to 256 or 128 pixels for initial classification.

Reference-free 2D classification, with at least 150 classes and ignoring the CTF until its first peak, was performed for at least 35 iterations to assess the presence of different polymorphs and cross-over distances. Polymorphs were identified by a novel hierarchical clustering approach that was inspired by the CHEP algorithm ^70^ (see below). Selected particles were re-extracted in boxes of 384 pixels for initial three-dimensional refinement. Initial three-dimensional references were generated *de novo* from 2D class average images using relion_helix_inimodel2d ^71^. For the FIA and structures which had low particle numbers (<5,000), a new algorithm using regularisation by denoising (to be described elsewhere) improved initial refinements, as conventional refinements resulted in high noise levels in the reconstruction due to overfitting. Subsequently, 3D classifications and 3D auto-refinements were used to select particles leading to the best reconstructions and optimise helical parameters. For some data sets, 3D classification was also used to separate out closely related polymorphs. For maps that were used for atomic modelling, Bayesian polishing ^65^ and CTF refinement ^72^ were used to increase resolution. Final maps were sharpened using standard post-processing procedures in RELION, and reported resolutions were estimated using a threshold of 0.143 in the Fourier shell correlation (FSC) between two independently refined half-maps (**Supplementary Information**).

### Polymorph identification and quantification

Picked filaments were hierarchically clustered by the unweighted pair group method with arithmetic mean (UPGMA), based on the cosine distance of the class distributions of particles for each filament from an initial 2D classification. Clusters were selected either by flattening the dendrogram at a specified threshold or interactively. Clusters below a minimum threshold of particles, typically 1000, were merged. Additional 2D classifications were performed for each identified cluster, iterating the clustering and 2D classification procedure until visually homogeneous populations of 2D classes were obtained. Filamentous class averages were then selected and output particles used for refinement.

Reported percentages of filaments in each data set were calculated based on the number of extracted particles used for the initial refinement of a particular filament type, relative to the total number of picked particles. For auto-picked data sets, an initial round of reference-free 2D classification was sometimes used to remove false positives from the picking procedure first. The reported percentages may not reflect the relative amounts of filament types in the original assembly reaction, because of limitations in our image analysis and because some filament types may disperse better than others in the grid holes.

Scripts for clustering filament types as well as for generating dataset summaries are available at https://github.com/dbli2000/FilamentTools.

### Atomic modelling

Atomic models were built either manually using COOT ^73^ or automatically using ModelAngelo ^74^. Coordinate refinement of models comprising 3 β-rungs was performed in ISOLDE ^75^. To ensure consistency, dihedral angles from the middle rung were applied to the top and bottom rungs. Subsequently, separate model refinements were performed on the first half-map for each refined structure. The resulting models were then evaluated by comparing them to this half-map (FSC_work_), as well as to the other half-map (FSC_test_) to monitor overfitting (**Supplementary Information**). Figures of structures, including electrostatic potential and hydrophobicity surfaces, were prepared using ChimeraX ^76^.

## Extended Data Figures

**Extended Data Figure 1:**
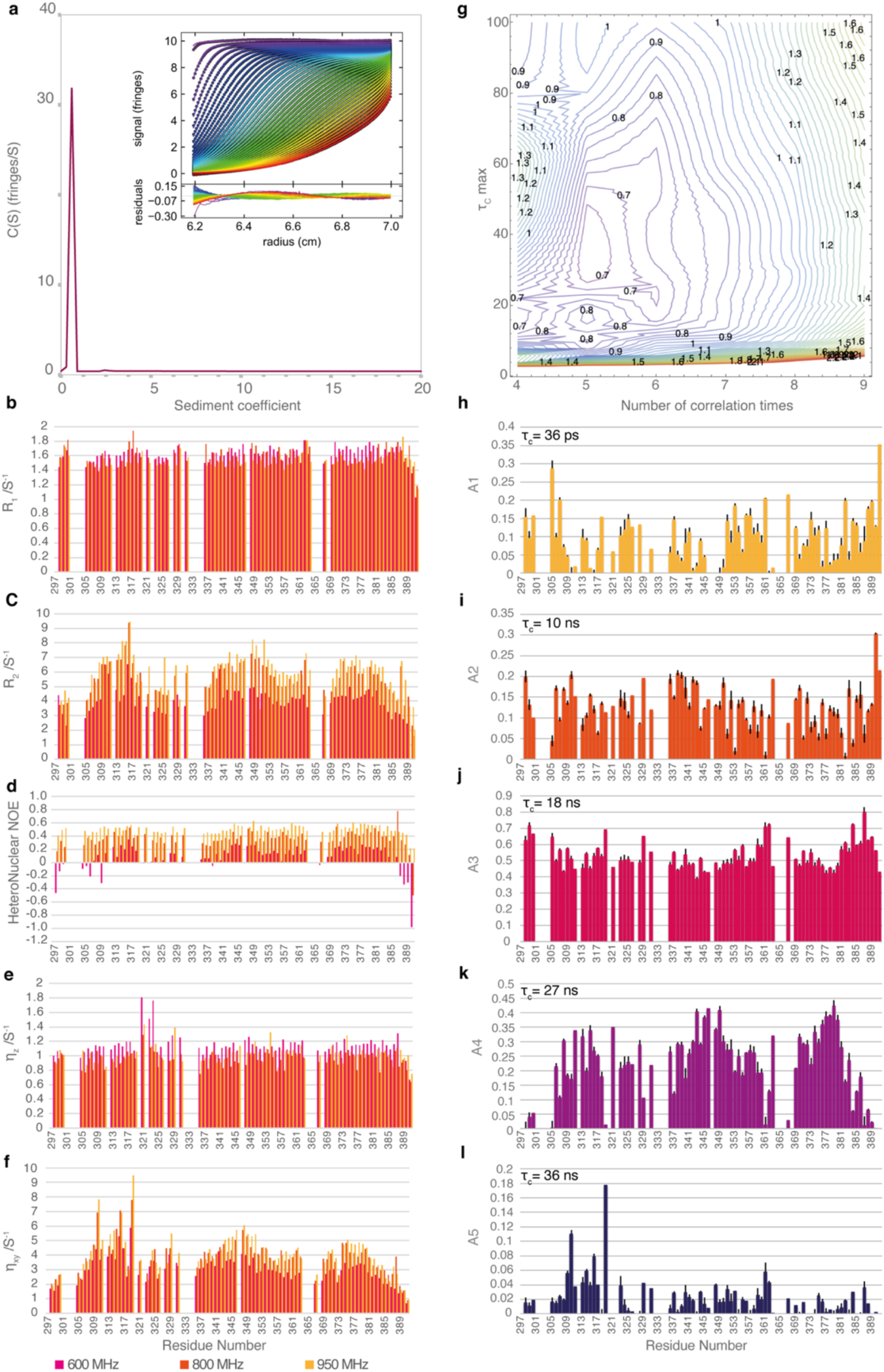
Analytical ultracentrifugation and nuclear magnetic resonance. **a.** Analytical ultracentrifugation (AUC) sedimentation velocity analysis of tau(297-391) in solution. The c(S) distribution shows tau(297-391) sedimented with coefficient of 0.6 S (Sw,20 = 1.0 S) with a frictional ratio of 1.777 corresponding to a mass of 10.3 kDa, consistent with a monomer in an extended conformation. The panel inset shows interference profiles with best fits of a c(S) model (coloured lines) and their residuals to the fits underneath. The different colours represent scans at different times: blue is the earliest time points where very little material has sedimented; through to red where the material has sedimented. **b-d.** Longitudinal R1, transverse R2 and heteronuclear ^15^N{^1^H} NOE measurements of tau(297-391) collected at 600 (magenta), 800 (orange) and 950 MHz (yellow) proton resonance frequencies. **e-f.** Exchange-free longitudinal and transverse cross-relaxation experiments collected at the same three proton resonance frequencies. **g.** Akaike’s information criterion for assessment of the fitness of a range of correlation times (between 4-9) and time scale conditions τmin – τmax (ranging from 2ps-2ns to 100ps-100ns) that best fit the spectra density analysis. **h-l.** Individual distributions of five coefficients (A1-A5) of the five correlation times τc = 36 ps, 10 ns, 18 ns, 27 ns and 36 ns as determined by the IMPACT analysis.

**Extended Data Figure 2:**
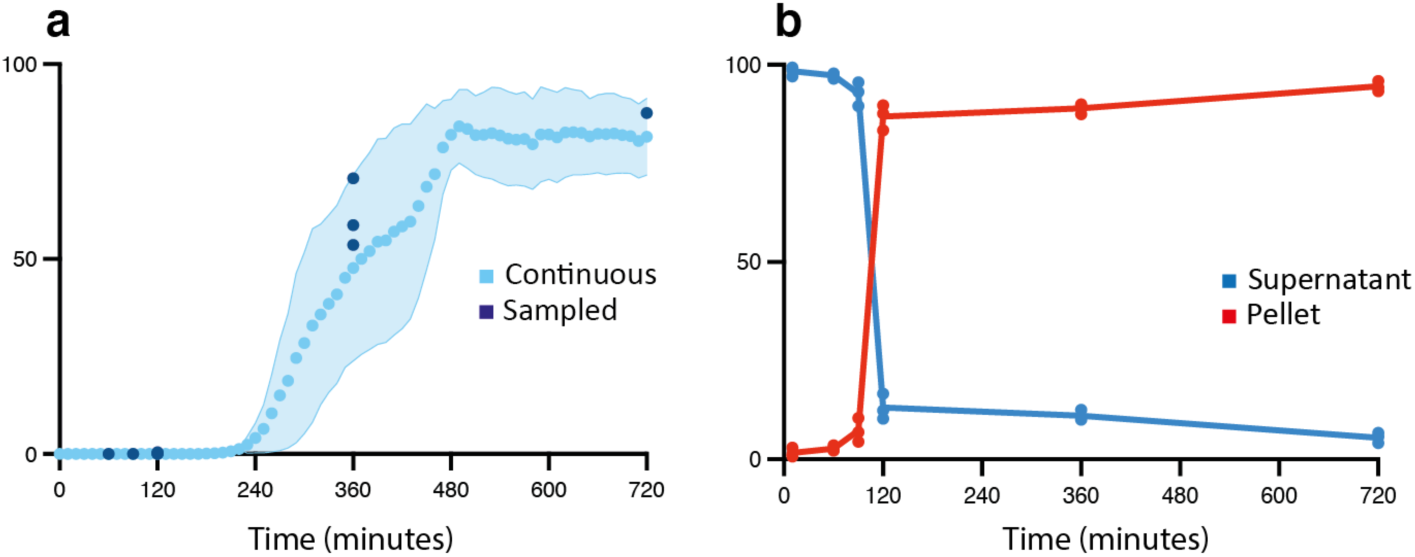
ThT fluorescence and pelletable tau in the CTE reaction. **a.** ThT fluorescence profile of the CTE reaction. Light blue circles indicate the average of 3 replicates; light blue shading indicates the standard deviation among 3 replicates; dark blue circles represent off-line ThT measurements. **b.** The amount of tau in the pellet and in the supernatant (in % of the total amount of tau) after centrifugation for 15 min at 400,000 g, as quantified by SDS-PAGE.

**Extended Data Figure 3:**
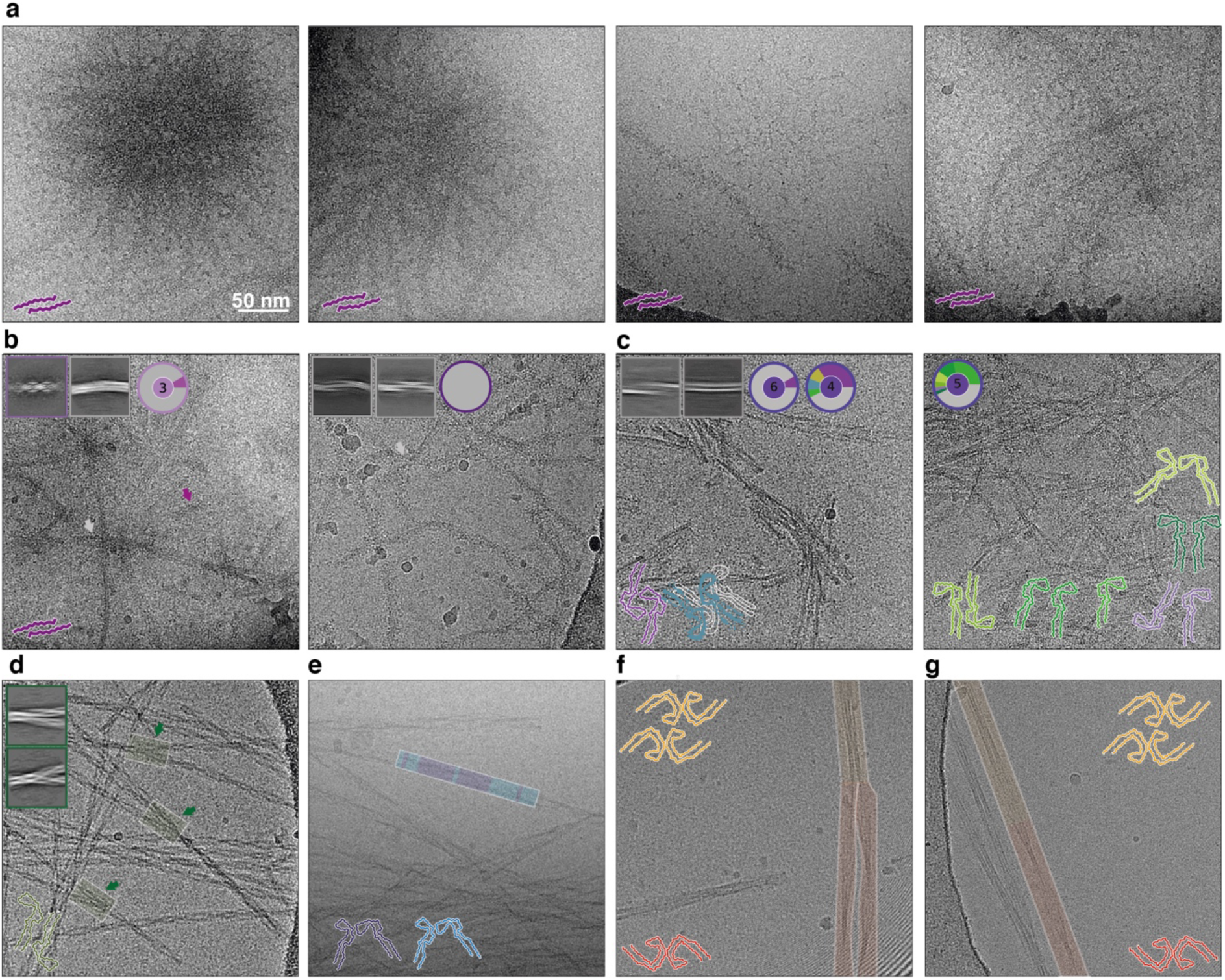
Various observations in micrographs. **a.** At 120 min, micrographs of the PHF reactions showed FIAs (cartoon at bottom left). In some instances, many FIAs appeared to originate from a single point, reminiscent of nucleation and growth of a crystal from an impurity (left two micrographs). In other instances, longer isolated FIAs were observed. **b.** At 140 min in the PHF reaction, some FIAs remained but most filaments yielded images that did not allow 3D reconstruction. Insets show 2D class averages; circular pie charts show the distribution of unsolvable filaments (grey) versus structures solved (coloured). **c.** At 160 min in the CTE reaction, nine structures could be solved from three replicates. **d.** At 180 min in the CTE reaction, some filament types appear to be branching, with specific structures in 2D class averages (insets). **e.** At 180 min in the CTE reaction, some filaments consist of multiple filament types (purple or blue). **f.** At 360 min in the PHF reaction, some QHFs (yellow) appear to be branching into two separate PHFs (red). **g.** At 360 min in the PHF reaction, some QHFs appear to convert into PHFs.

**Extended Data Figure 4:**
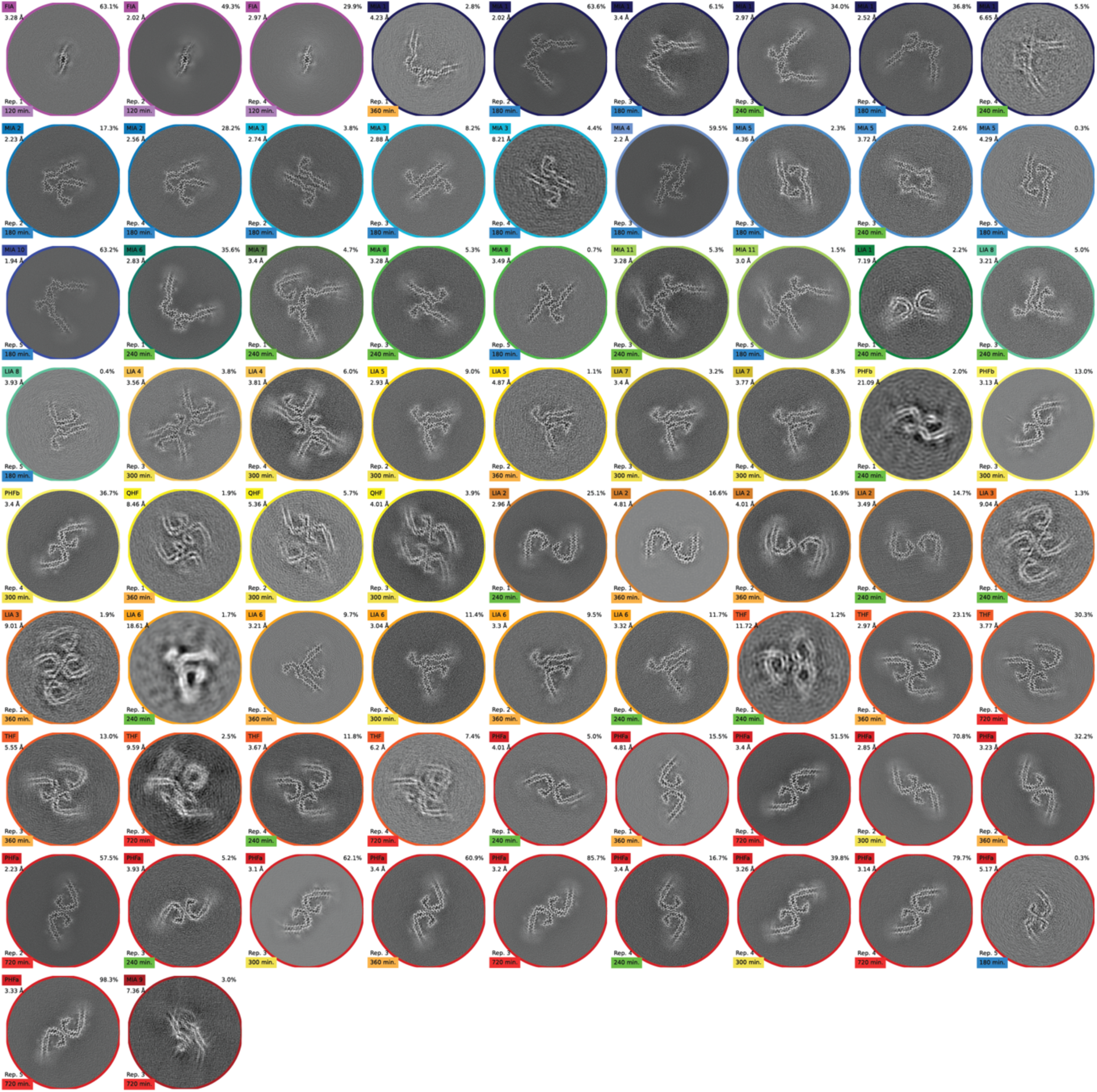
Cryo-EM reconstructions from the PHF reaction. **a.** Projected slices, with an approximate thickness of 4.7 Å, orthogonal to the helical axis for the filaments formed in the PHF reaction. Filament names and resolutions are indicated in the top left; percentages of filament types in each cryo-EM data set are shown in the top right and the replicate and time point are indicated in the bottom left of each image. Circles around the slices are coloured as the structures of Figure 4 in the main text.

**Extended Data Figure 5:**
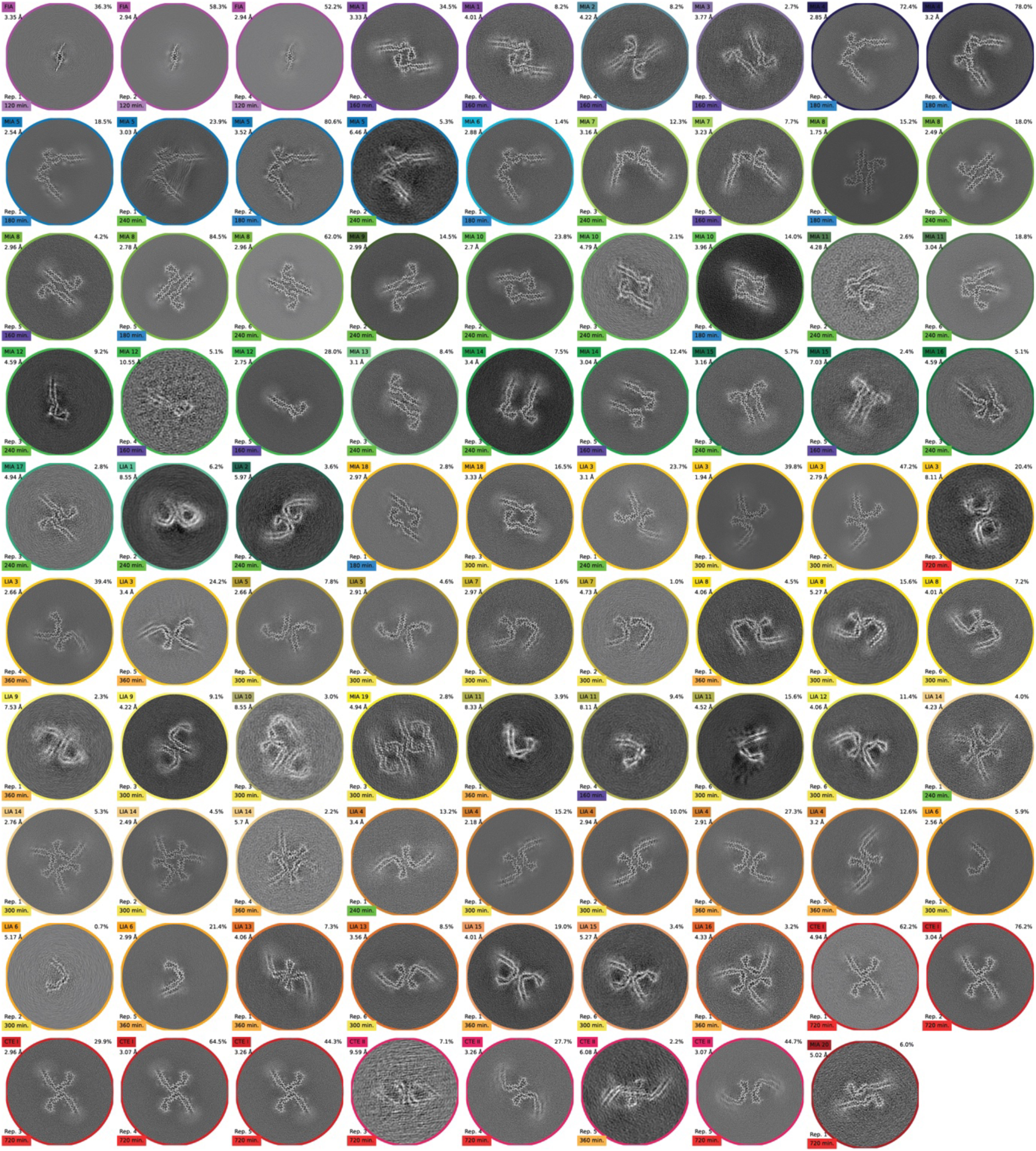
Cryo-EM reconstructions from the CTE reaction. **a.** Projected slices, with an approximate thickness of 4.7 Å, orthogonal to the helical axis for the filaments formed in the CTE reaction. Filament names and resolutions are indicated in the top left; percentages of filament types in each cryo-EM data set are shown in the top right and the replicate and time point are indicated in the bottom left of each image. Circles around the slices are coloured as the structures of Figure 4 in the main text.

**Extended Data Figure 6:**
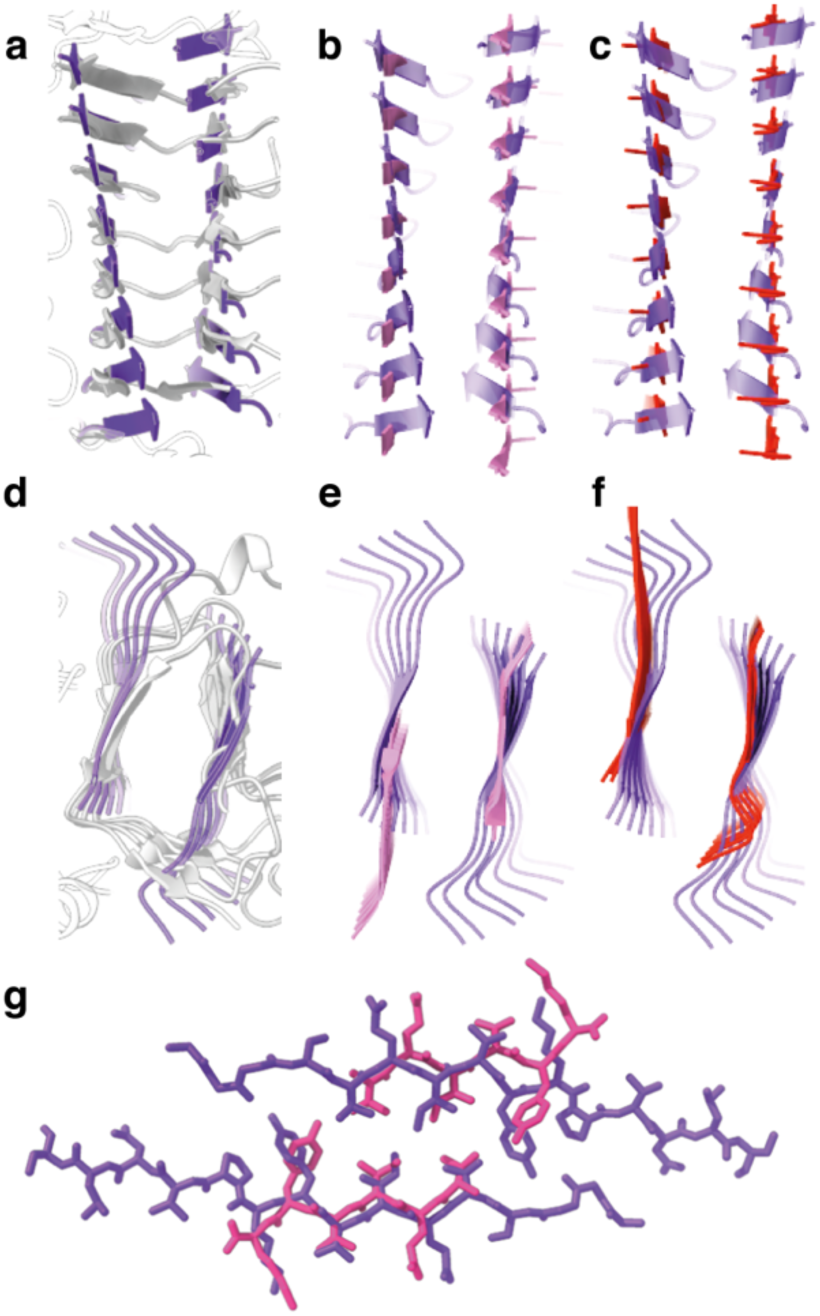
Twisted β-sheets in the FIA and other structures. **a.** Side view of the FIA (purple) and PDB entry 1PCL (grey) ^29^, illustrating that they adopt similarly twisted β-sheets. **b.** Side view of PDB entry 2ON9 (pink) ^30^ and the FIA (transparent purple). β-Sheets in the crystal structure of the 306VQIVYK311 peptide are straight. **c.** Side view of a PHF at 720 min (red) and the FIA (transparent purple), illustrating that β-sheets in the PHF are less twisted than in the FIA. **d-f.** As in panel a-c, but top views. **g.** Top view of an all-atom representation of the FIA and crystal structure 2ON9.

**Extended Data Figure 7:**
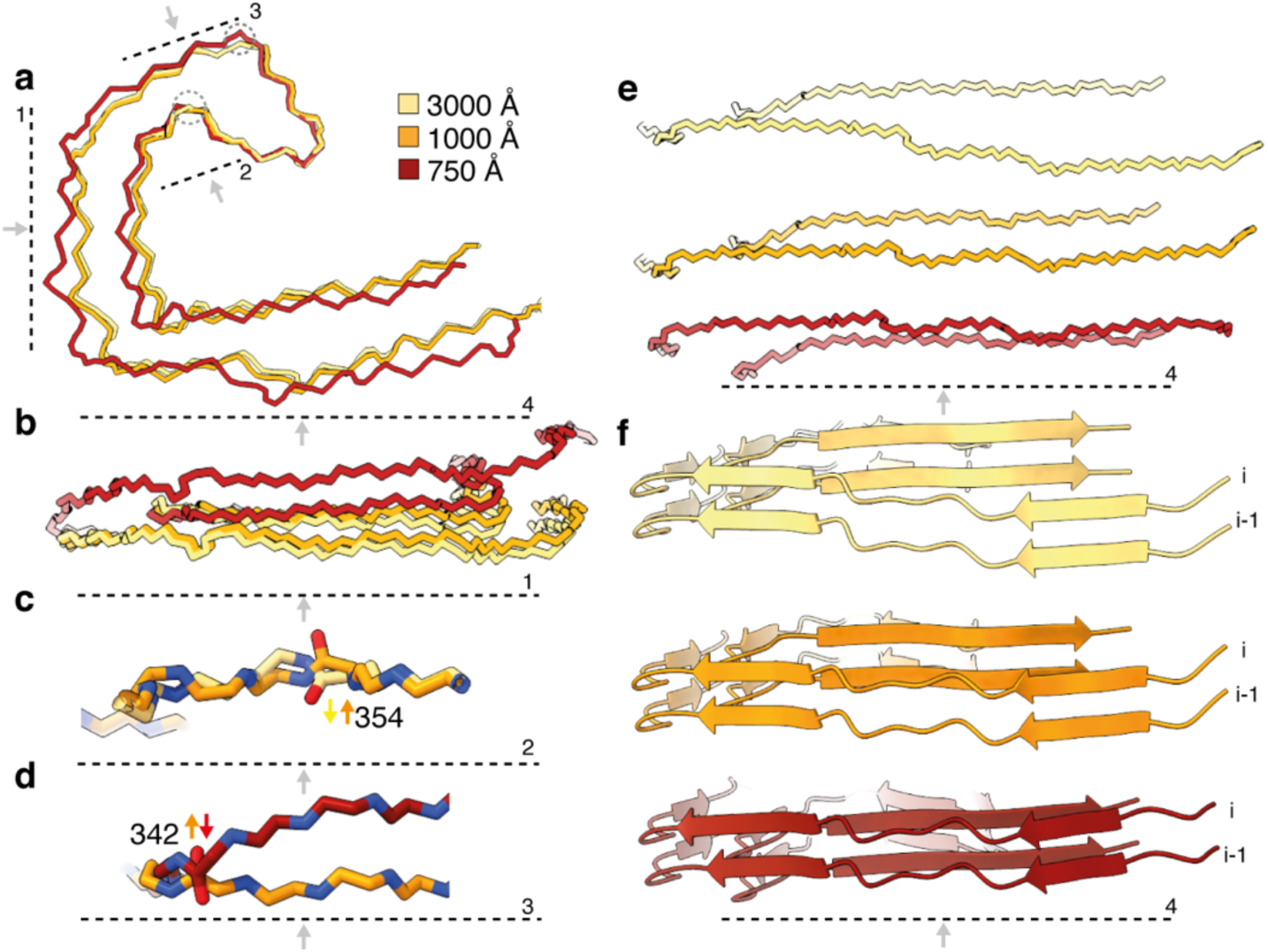
PHF crossover distances. **a.** Backbone traces, aligned at amino acids 339-354, for PHFs with crossover distances of 750 Å (red), 1000 Å (orange) and 3000 Å (yellow). Grey dotted lines and arrows indicate viewing planes and directions in subsequent panels. **b.** Side view of amino acids 305-320 and 365-380. **c.** The carbonyl of isoleucine 354 flips from PHFs with a crossover distance of 1000 Å, compared to those with a crossover distance of 3000 Å. **d.** The carbonyl of glutamic acid 342 flips from PHFs with a crossover distance of 750 Å, compared to those with a crossover distance of 1000 Å. **e.** Side view of backbone traces of amino acids 305-320 and 365-380 for PHFs with crossover distances of 3000 Å (top), 1000 Å (middle) and 750 Å (bottom). **f.** As in panel e, but for cartoon representations of two β-rungs.

**Extended Data Figure 8:**
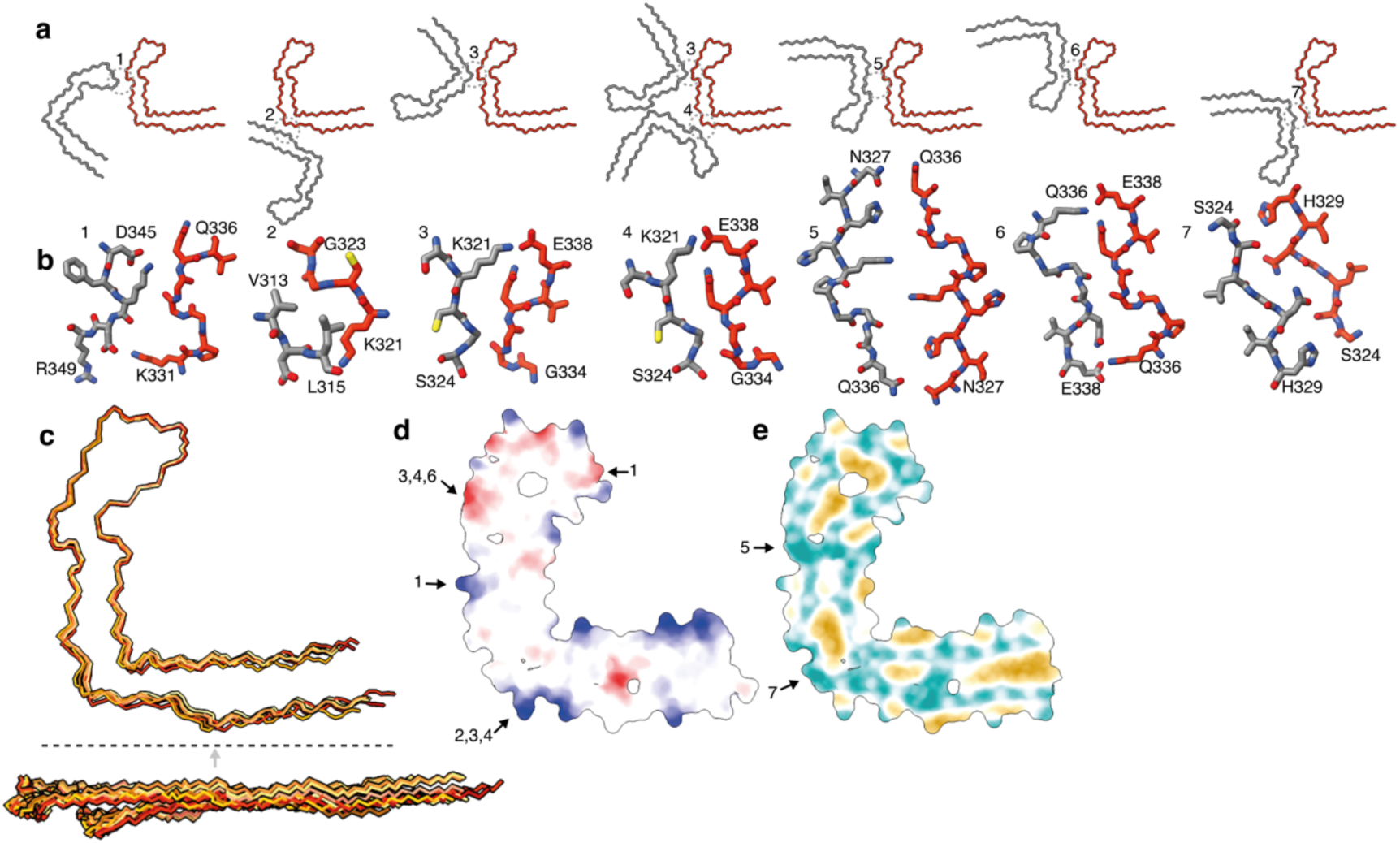
CTE protofilament interactions. **a.** Backbone traces for C-shaped structures that are present at 300-720 min in the CTE reaction. The red protofilament is aligned across all structures; the grey protofilaments adopt varying orientations relative to the red one. The dashed circles and numbers are referred to in panels b, d and e. **b.** All-atom representation of the protofilament interactions shown in panel a. **c.** A superposition of backbone traces (top) and side view (bottom) shows that the different protofilaments from panel a adopt similar conformations. **d.** Coulomb electrostatic potential (positive charges in blue; negative charges in red) of the CTE protofilament. **e.** Hydrophobicity representation (hydrophobic parts in yellow; charged parts in cyan).

