## Supplementary information for "Disease-specific tau filaments assemble via polymorphic intermediates"

### Overview

#### Summaries of data sets from the AD (p. 2-22) and CTE (p. 23-49) reactions

**a.** Hierarchical classification of individual filament segments according to their assigned 2D class average (vertical) and the picked filament ID(horizontal). **b.** Pie chart with the relative amounts of different filament types. Grey represents unsolved filaments. Filament types are the same as in Figure 4 of the main text. Different colours represent different time points (120 min in purples; 180 min in blues; 240 min in greens; 300 min in yellows; 360 min in oranges and 720 min in reds). Structures are coloured according to the time point at which they are most abundant, averaged across all replicates. Unique names of filament types are indicated and the same names are used throughout this document. **c.** Top 10 2D class averages of unsolved filaments (classes containing >50% unused particles) as ordered by number of particles. **d.** XY-cross-sections, with a projected depth of approximately 4.7 Å for each filament type.

#### Resolution estimates for structures from the AD (p. 50) and CTE (p. 51-52) reactions

For each structure, solvent-corrected Fourier-shell correlation (FSC) curves between independently refined half-maps are shown in black; FSC curves between the refined model and the reconstruction from all particles are shown in red; FSC curves between a model refined against half map 1 against half map 1 are shown in dashed yellow; FSC curves of the same model against half map 2 are shown in blue.

#### Cryo-EM data acquisition parameters (p. 53)

Data acquisition parameters for the G4 and G2 Titan Krios microscopes at Thermo Fisher Scientific (TFS) and at the Laboratory of Molecular Biology (LMB), respectively. Statistics about cryo-EM reconstructions are reported separately in the Supplementary Information.

#### Atomic structure refinement statistics from the AD (p. 54-64) and CTE (p. 65-76) reactions

Refinement statistics are shown for the atomic models that were refined in the highest-resolution map available for each unique structure.

#### SDS-PAGE gels for the quantification of pelletable tau (p. 77)

Coomassie stained sodium dodecyl-sulfate polyacrylamide gel electrophoresis (SDS-PAGE, 4-20% Tris-glycine) of pelletable material in the assembly reactions. 1.5 uL of supernatant or pellet was loaded onto each well. **a.** For the AD reaction. **b.** For the CTE reaction. S: supernatant, P: pellet. Time is denoted in minutes.

**a**

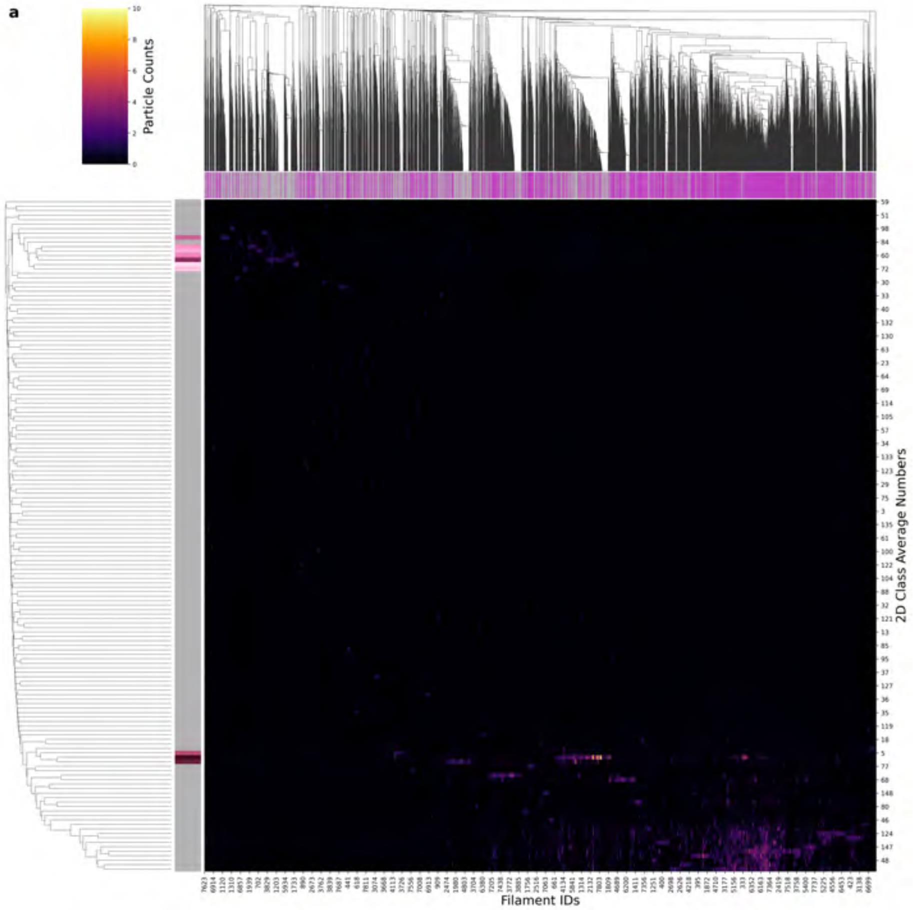

| Condition | PHF |
|---|---|
| **Timepoint** | 120 minutes |
| **Replicate** | 1 |
| **Picking** | Auto |
| **Micrographs** | 9293 |
| **Extracted Particles** | 1714140 |
| **Pixel Size** | 0.727 Å |
| **Classified Particles** | 60982 |

**b**

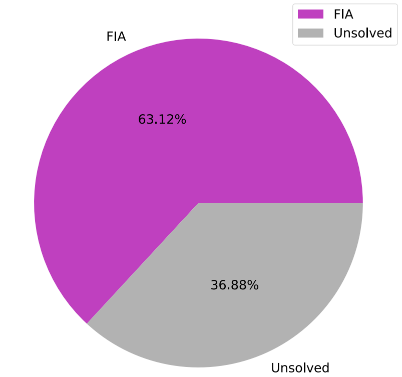

**c**

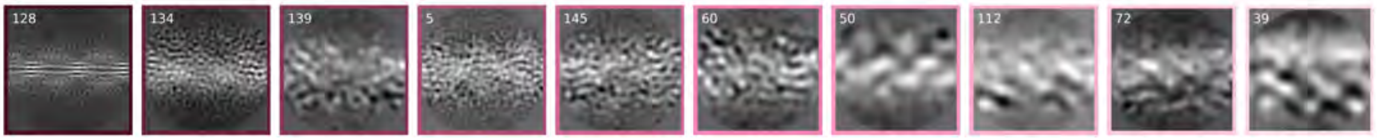

**d**

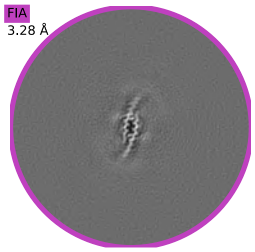

a

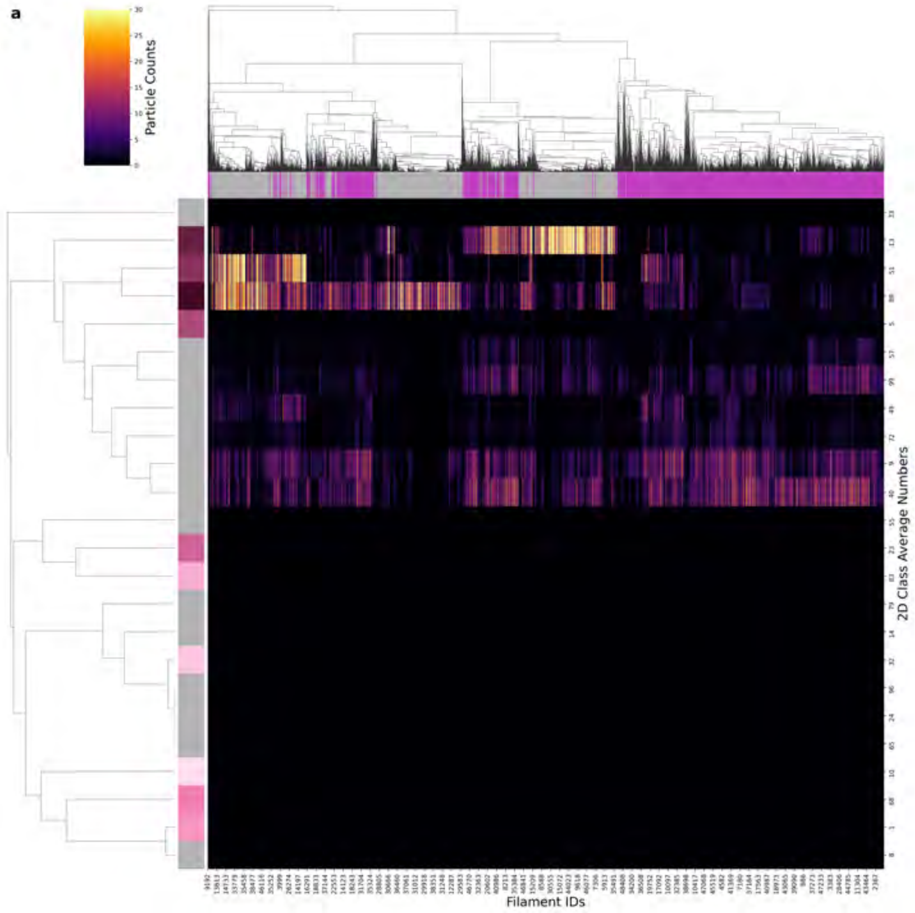

| Condition | PHF |
|---|---|
| Timepoint | 120 minutes |
| Replicate | 2 |
| Picking | Auto |
| Micrographs | 9340 |
| Extracted Particles | 1680613 |
| Pixel Size | 0.876 Å |
| Classified Particles | 1680613 |

b

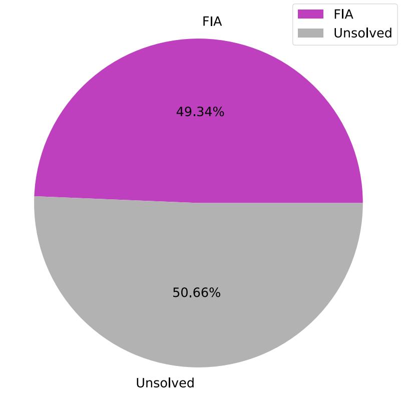

c

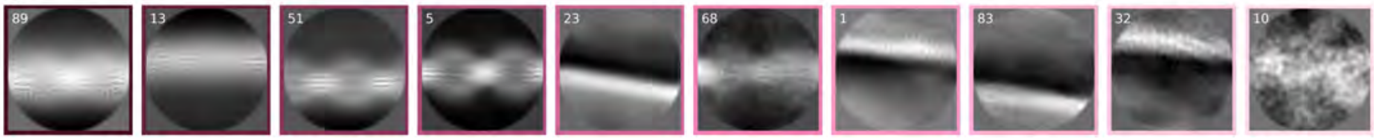

d

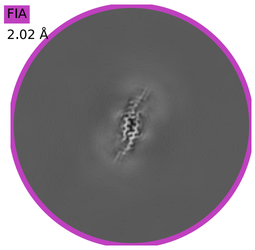

a

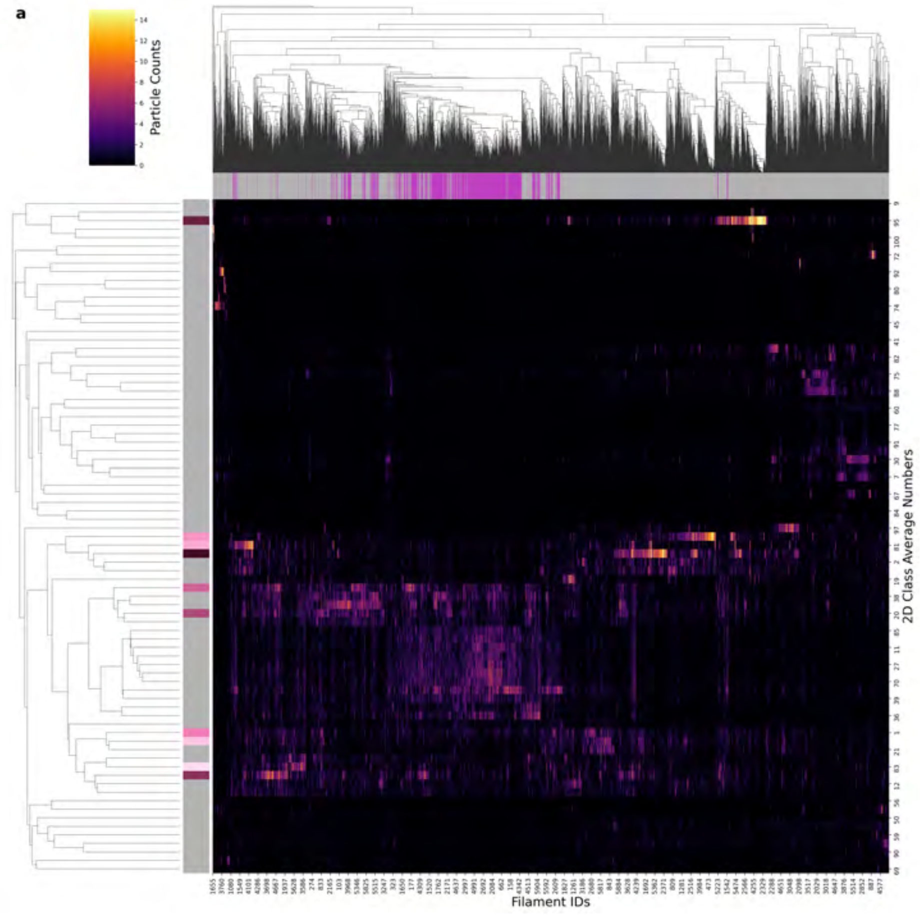

| Condition | PHF |
|---|---|
| Timepoint | 120 minutes |
| Replicate | 4 |
| Picking | Auto |
| Micrographs | 1503 |
| Extracted Particles | 197780 |
| Pixel Size | 0.727 Å |
| Classified Particles | 197780 |

b

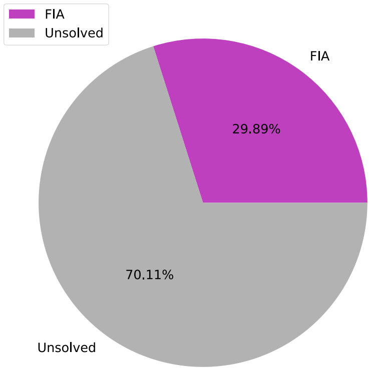

c

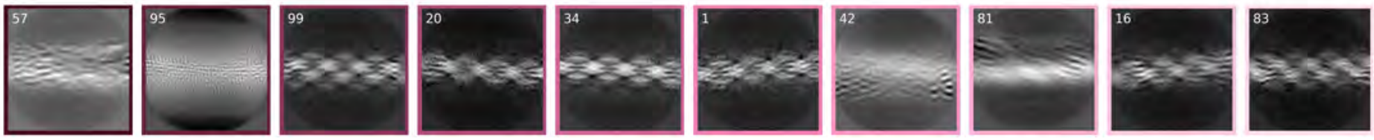

d

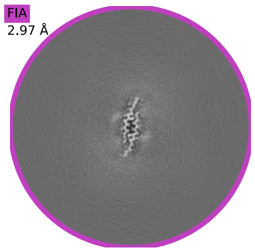

a

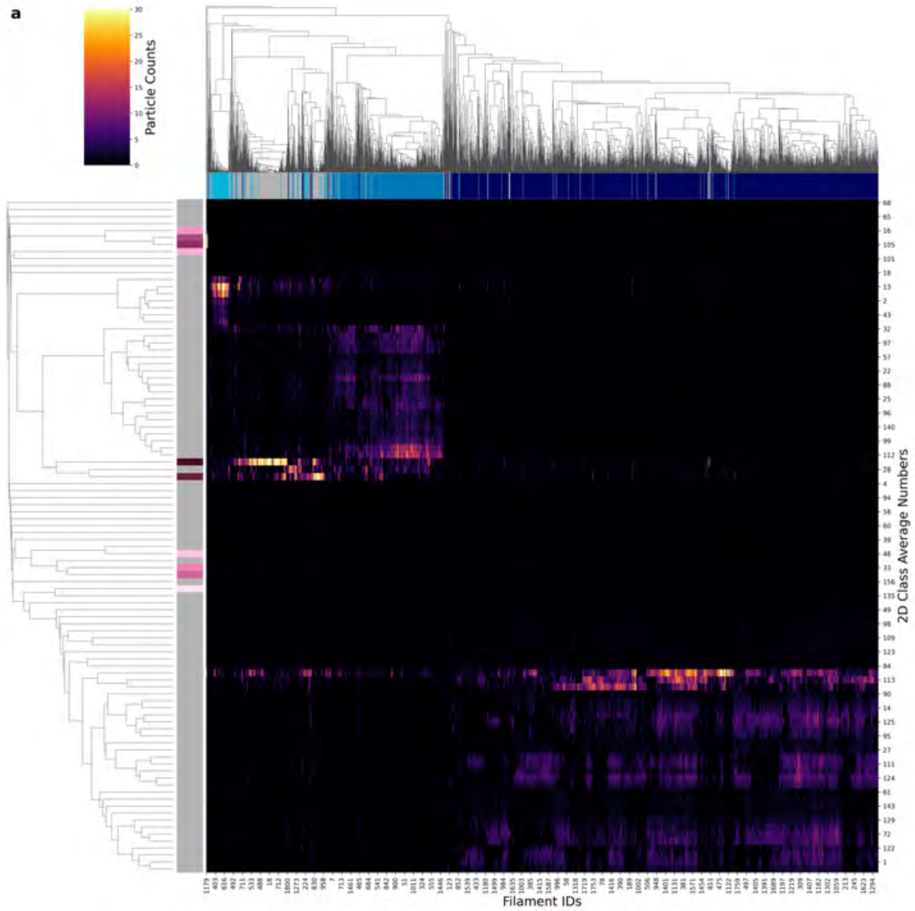

| Condition | PHF |
|---|---|
| Timepoint | 180 minutes |
| Replicate | 2 |
| Picking | Manual |
| Micrographs | 2430 |
| Extracted Particles | 122414 |
| Pixel Size | 0.727 Å |
| Classified Particles | 122414 |

b

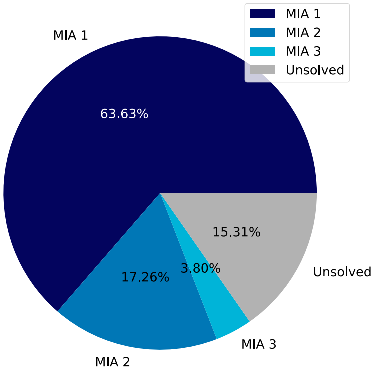

c

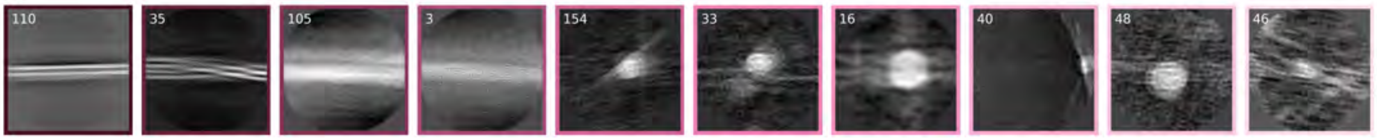

d

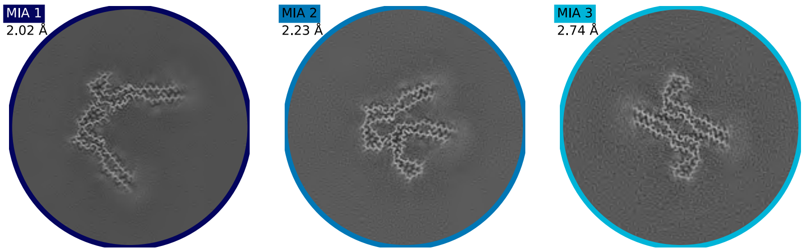

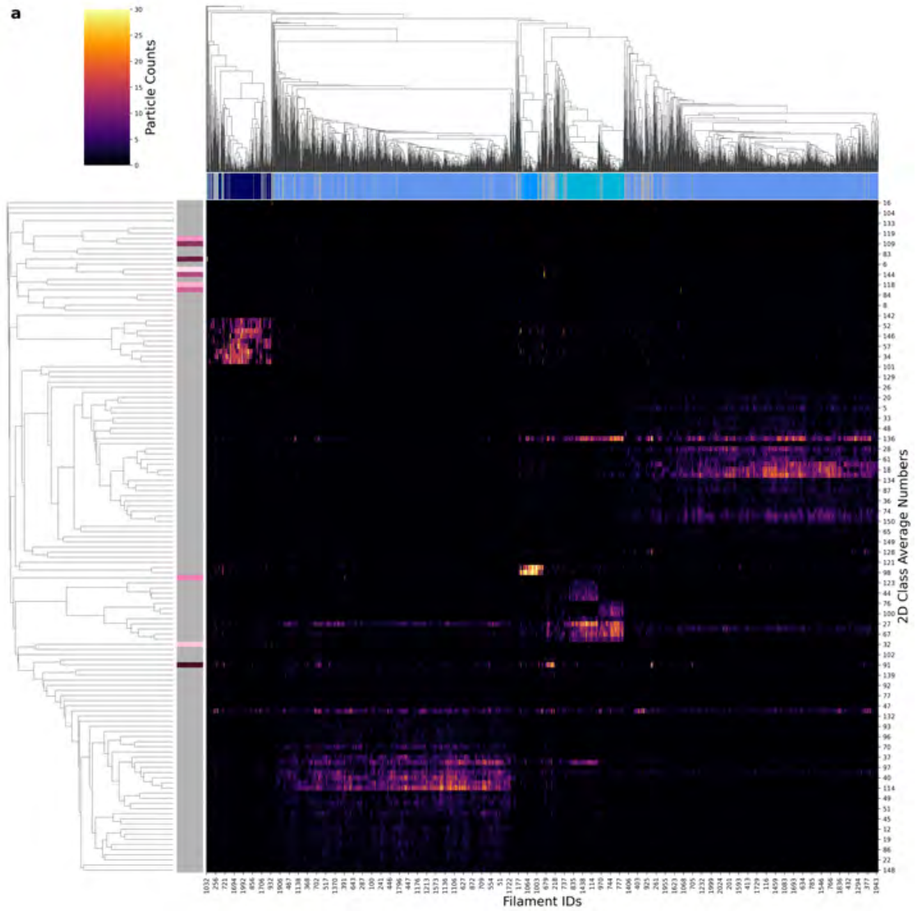

| Condition | PHF |
|---|---|
| Timepoint | 180 minutes |
| Replicate | 3 |
| Picking | Manual |
| Micrographs | 2427 |
| Extracted Particles | 159594 |
| Pixel Size | 0.727 Å |
| Classified Particles | 159594 |

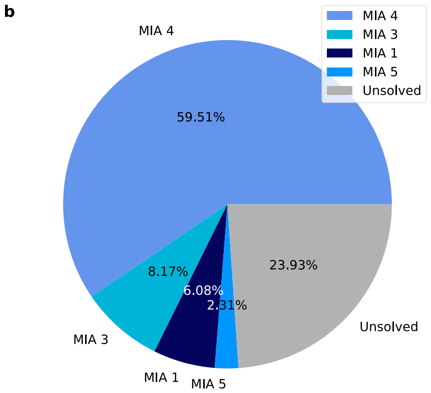

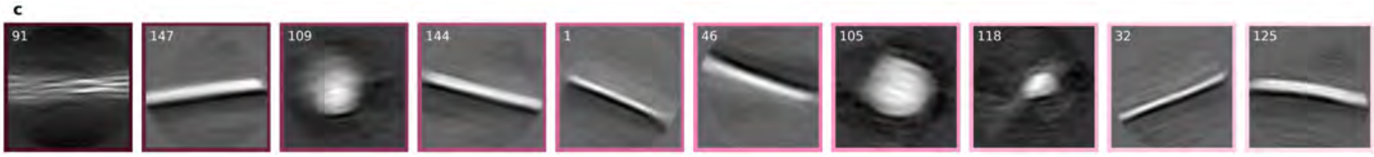

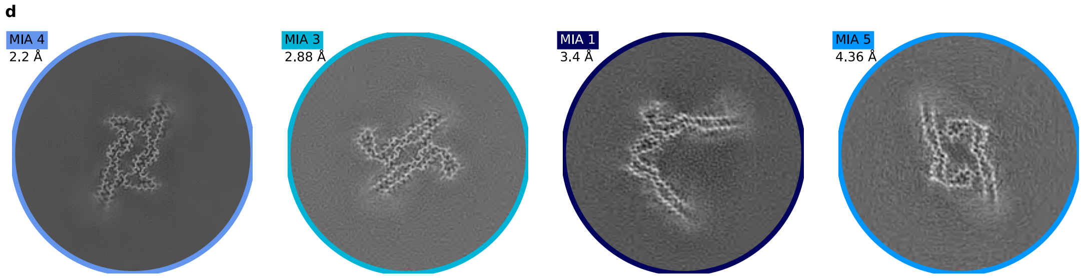

a

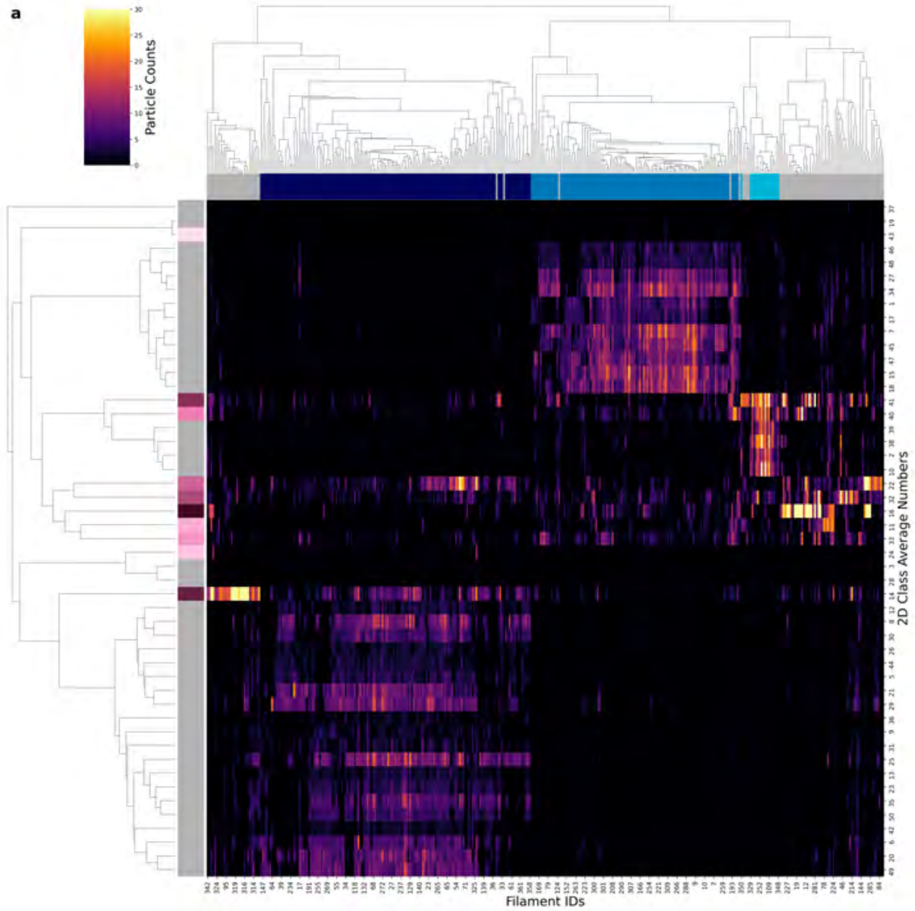

| | |
|---|---|
| **Condition** | PHF |
| **Timepoint** | 180 minutes |
| **Replicate** | 4 |
| **Picking** | Manual |
| **Micrographs** | 596 |
| **Extracted Particles** | 30133 |
| **Pixel Size** | 0.727 Å |
| **Classified Particles** | 30133 |

b

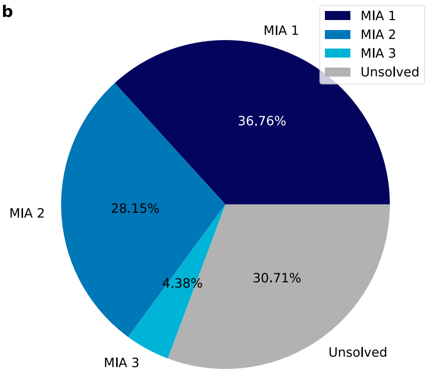

c

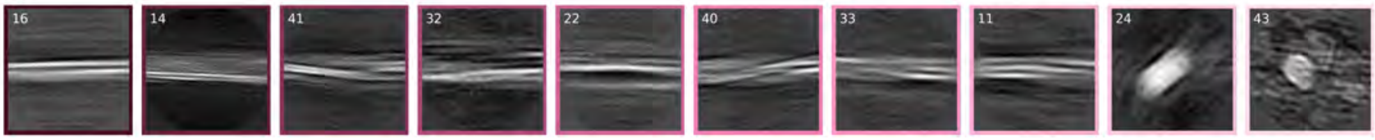

d

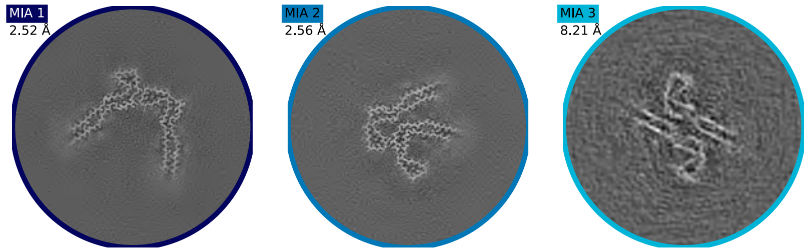

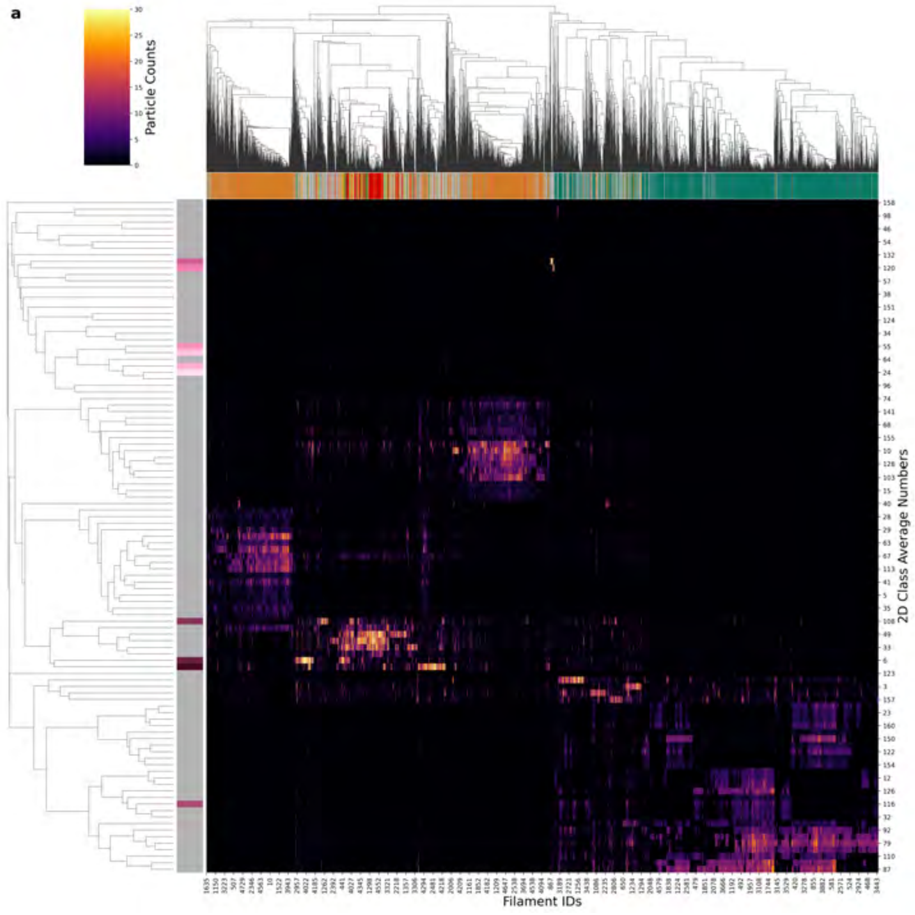

| Condition | PHF |
|---|---|
| Timepoint | 240 minutes |
| Replicate | 1 |
| Picking | Manual |
| Micrographs | 7262 |
| Extracted Particles | 352893 |
| Pixel Size | 0.824 Å |
| Classified Particles | 352893 |

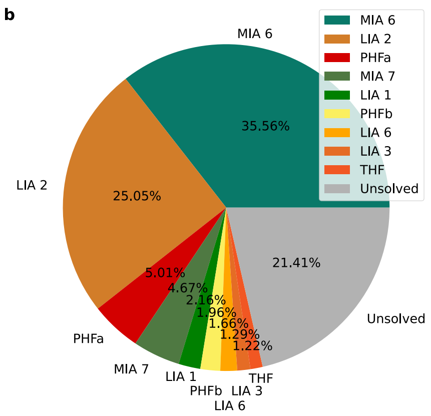

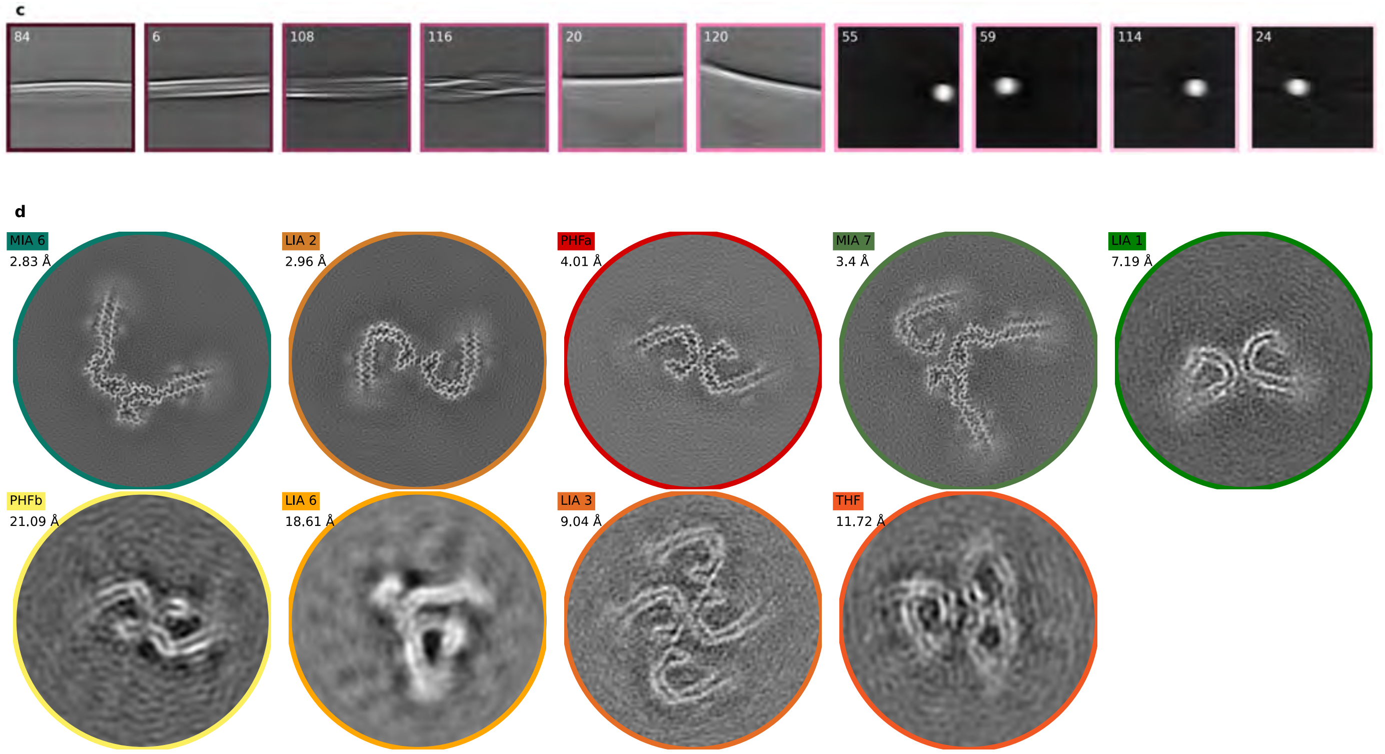

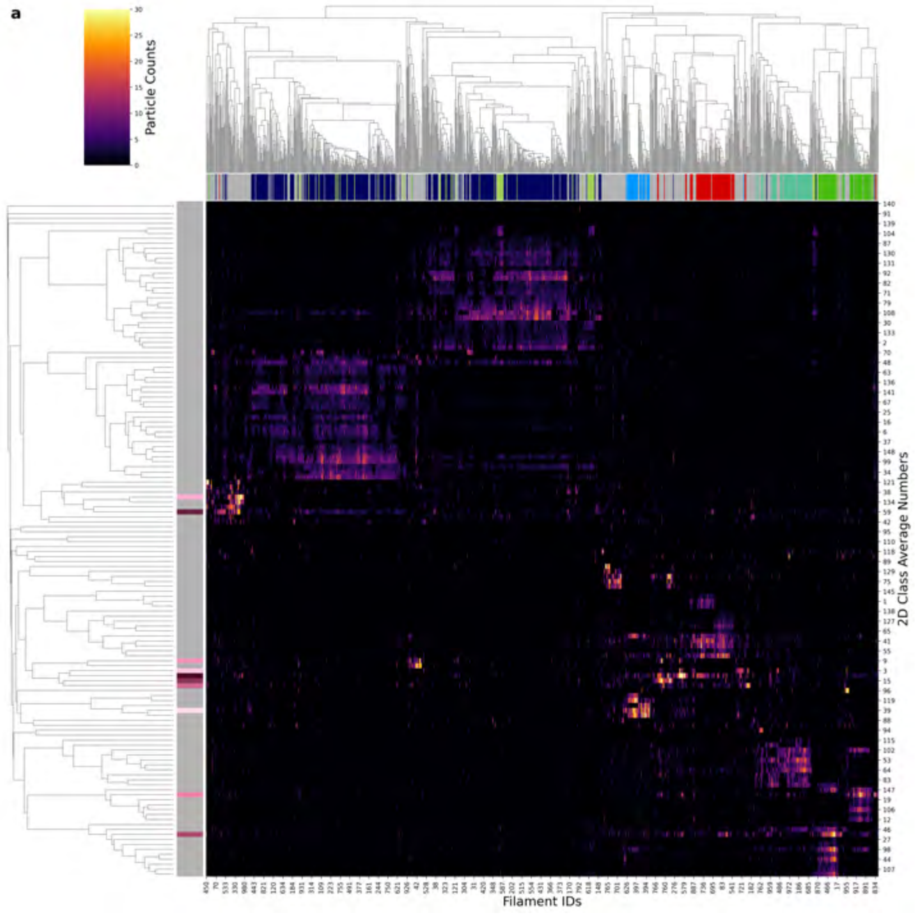

| Condition | PHF |
|---|---|
| Timepoint | 240 minutes |
| Replicate | 3 |
| Picking | Manual |
| Micrographs | 2320 |
| Extracted Particles | 80292 |
| Pixel Size | 0.727 Å |
| Classified Particles | 80292 |

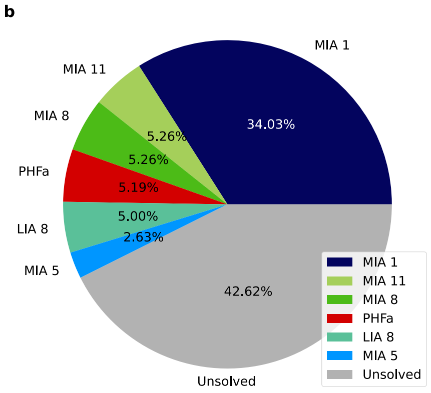

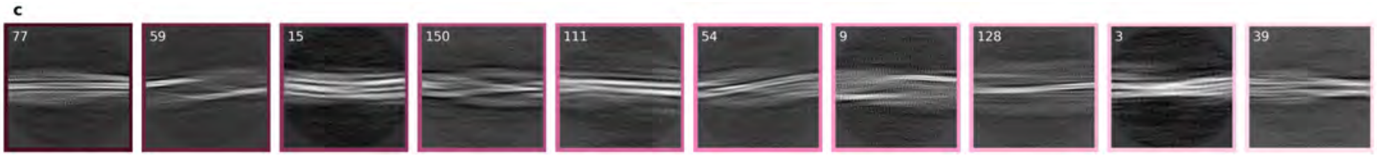

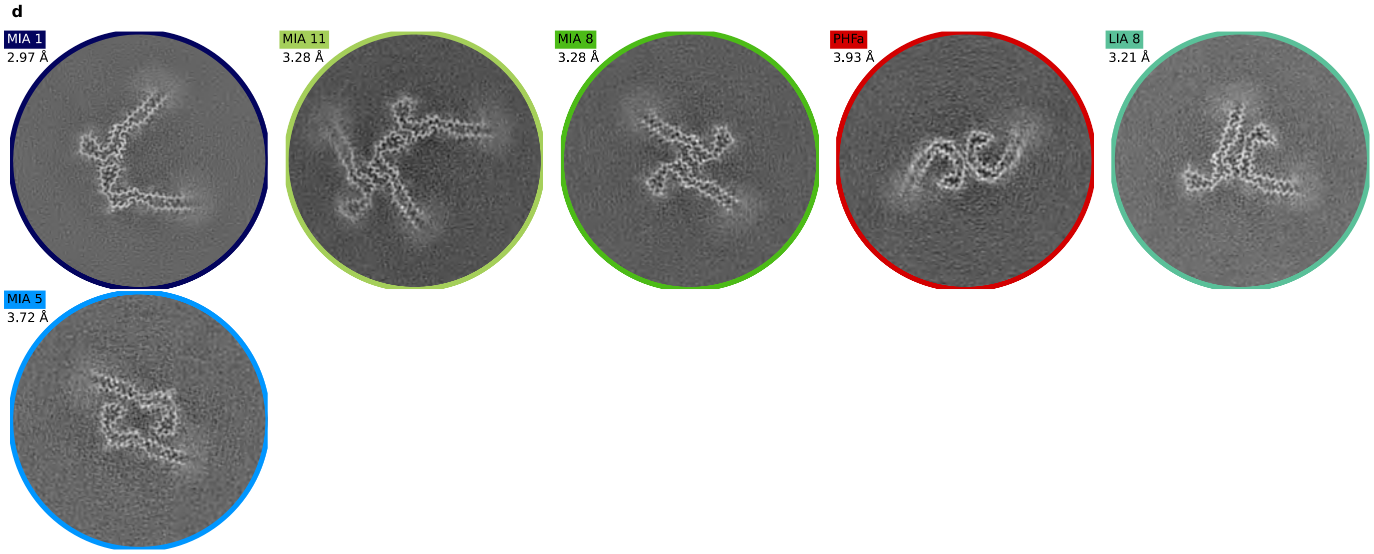

a

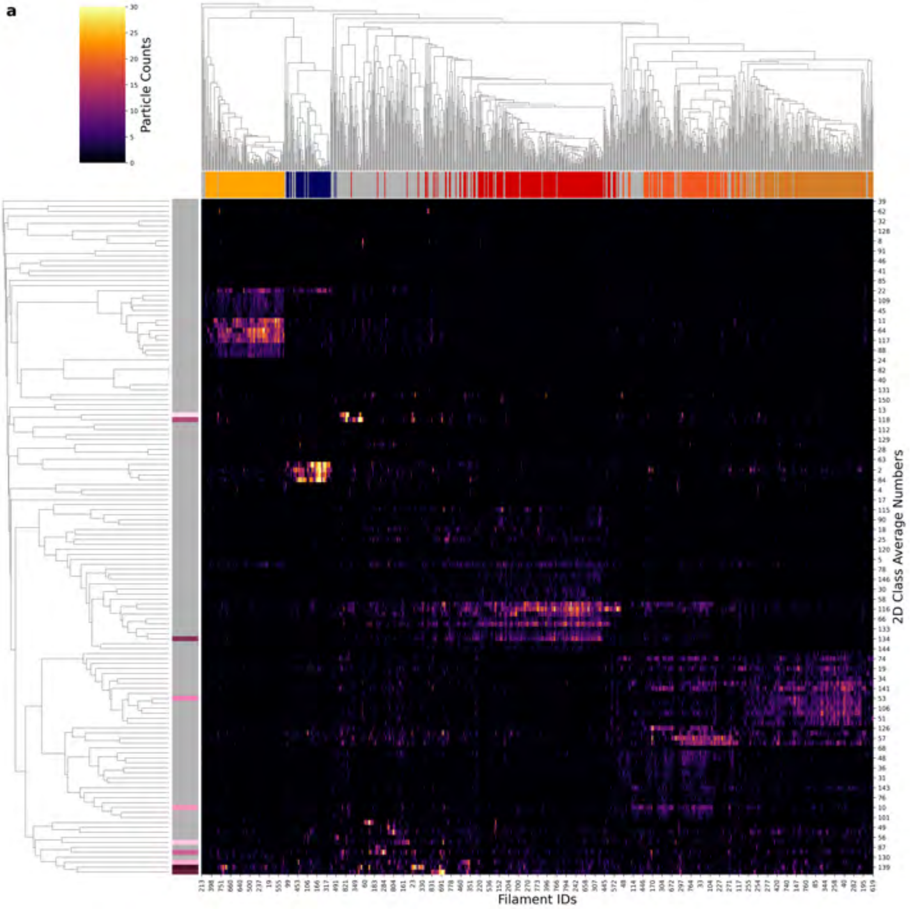

| Condition | PHF |
|---|---|
| Timepoint | 240 minutes |
| Replicate | 4 |
| Picking | Manual |
| Micrographs | 2560 |
| Extracted Particles | 72343 |
| Pixel Size | 0.727 Å |
| Classified Particles | 72343 |

b

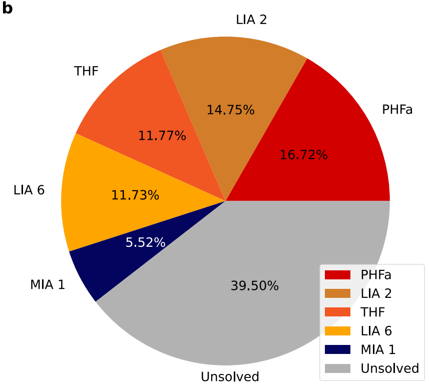

c

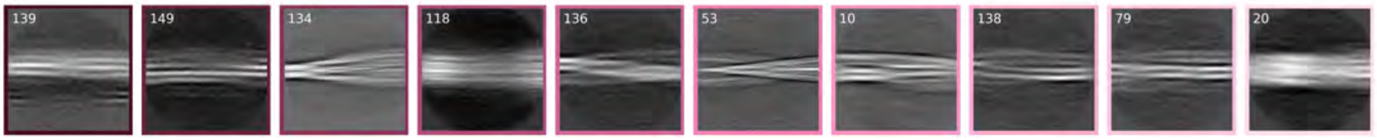

d

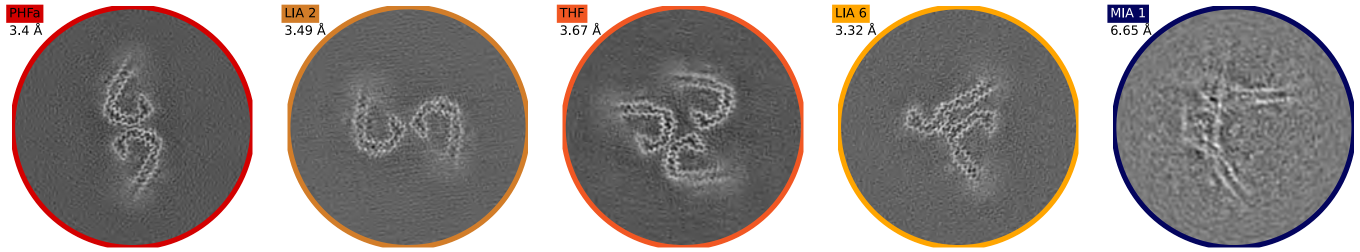

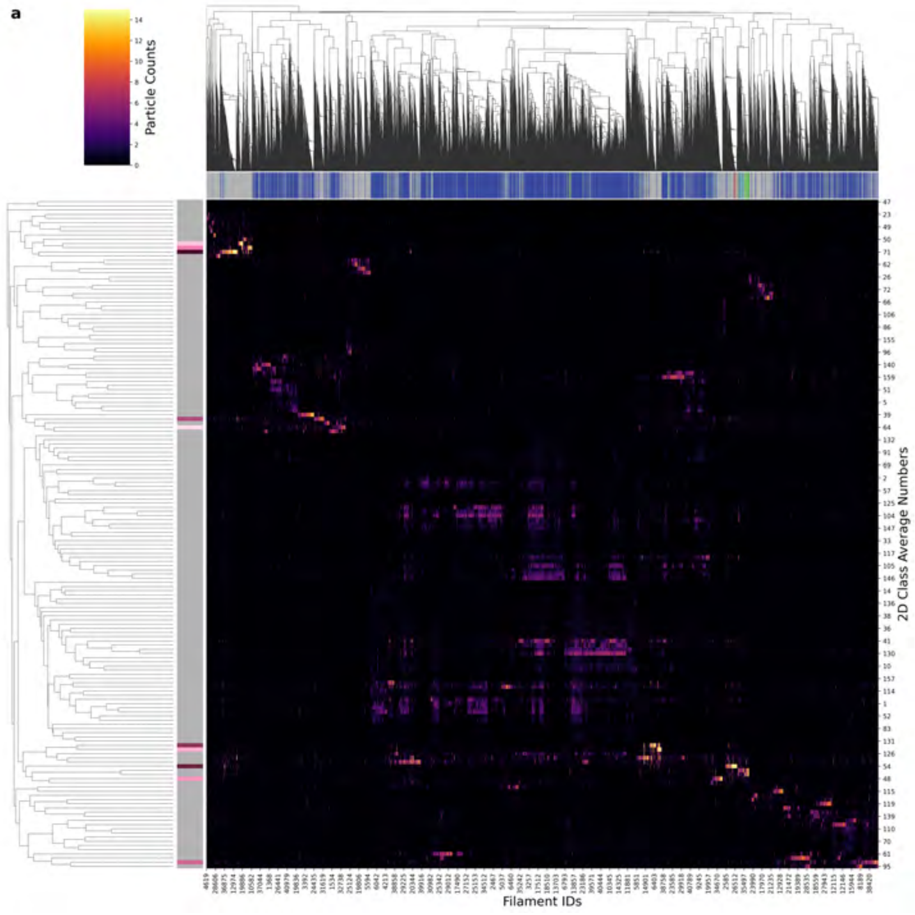

| Condition | PHF |
|---|---|
| Timepoint | 240 minutes |
| Replicate | 5 |
| Picking | Auto |
| Micrographs | 2228 |
| Extracted Particles | 1178114 |
| Pixel Size | 0.727 Å |
| Classified Particles | 1178114 |

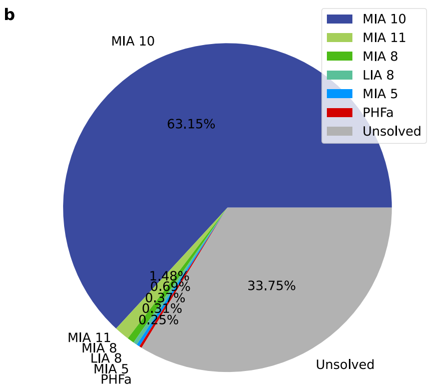

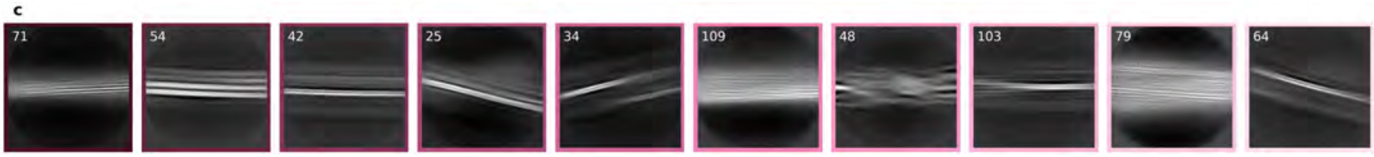

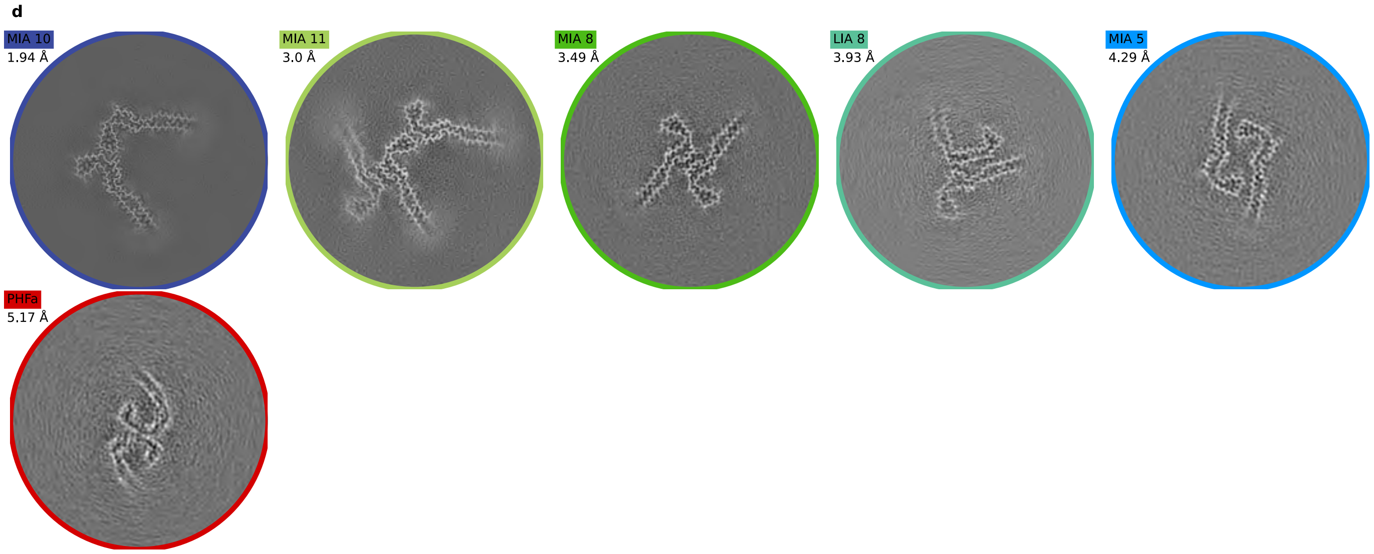

**a**

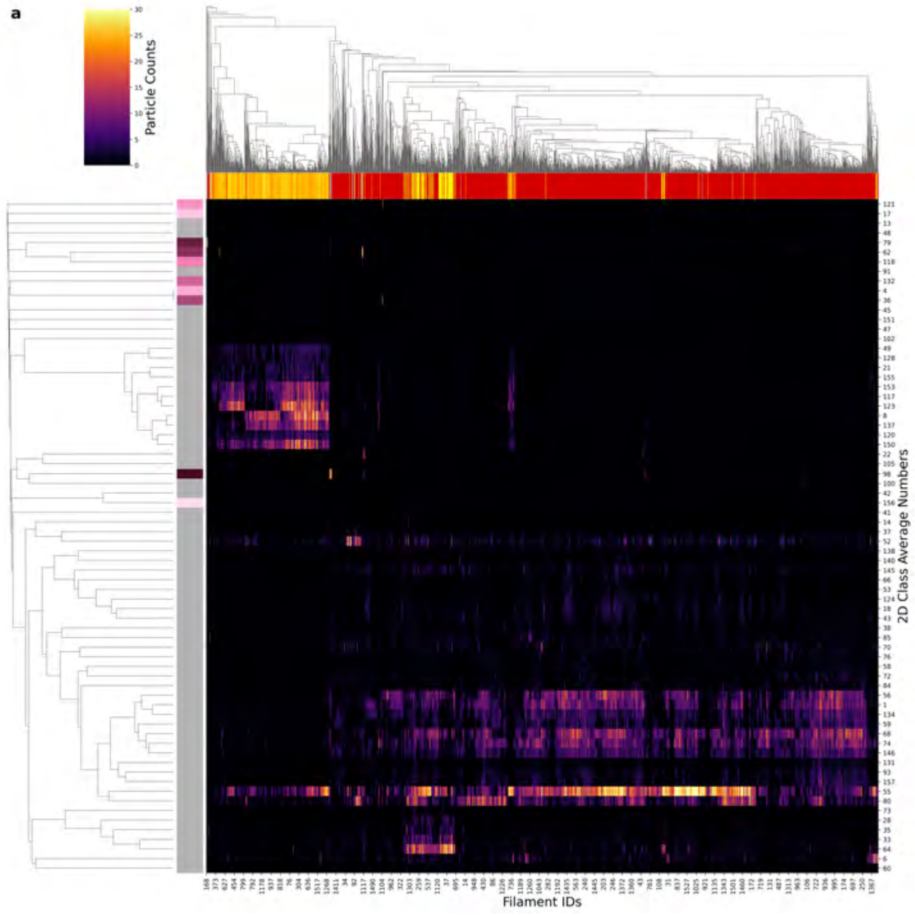

| Condition | PHF |
|---|---|
| Timepoint | 300 minutes |
| Replicate | 2 |
| Picking | Manual |
| Micrographs | 4083 |
| Extracted Particles | 103645 |
| Pixel Size | 0.824 Å |
| Classified Particles | 103645 |

**b**

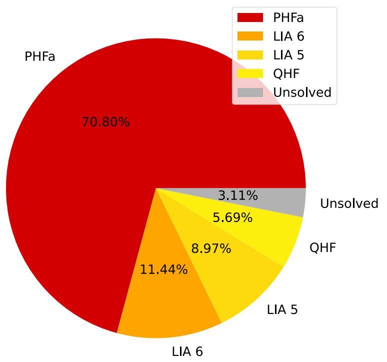

**c**

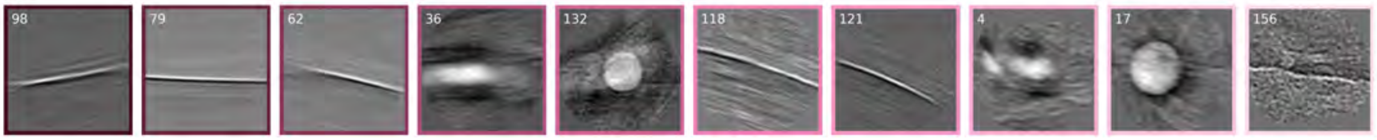

**d**

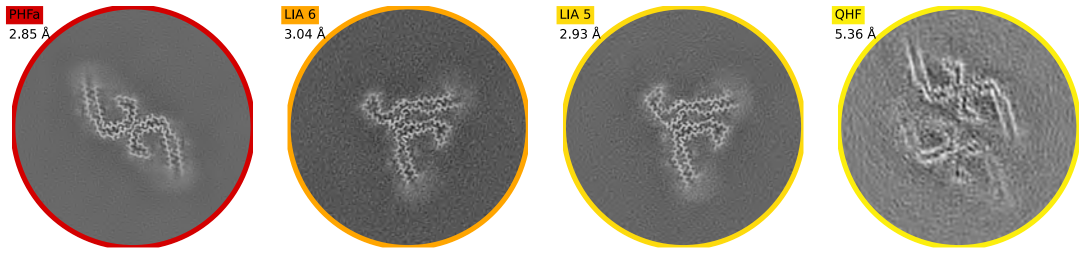

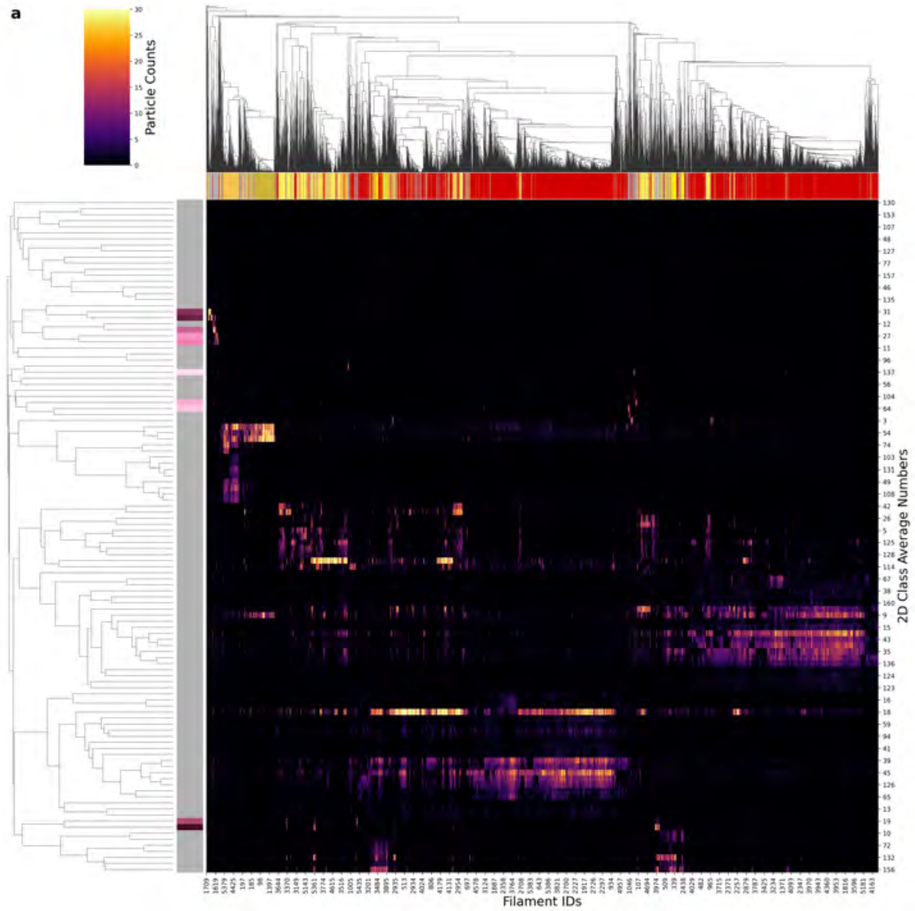

| Condition | PHF |
|---|---|
| Timepoint | 300 minutes |
| Replicate | 3 |
| Picking | Manual |
| Micrographs | 3923 |
| Extracted Particles | 441748 |
| Pixel Size | 0.824 Å |
| Classified Particles | 441748 |

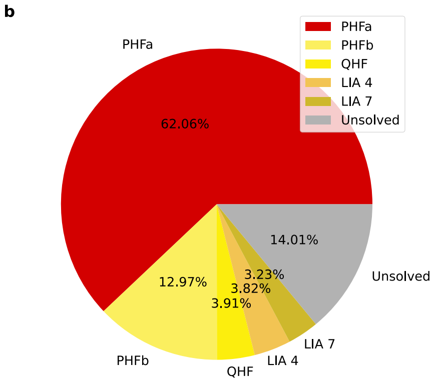

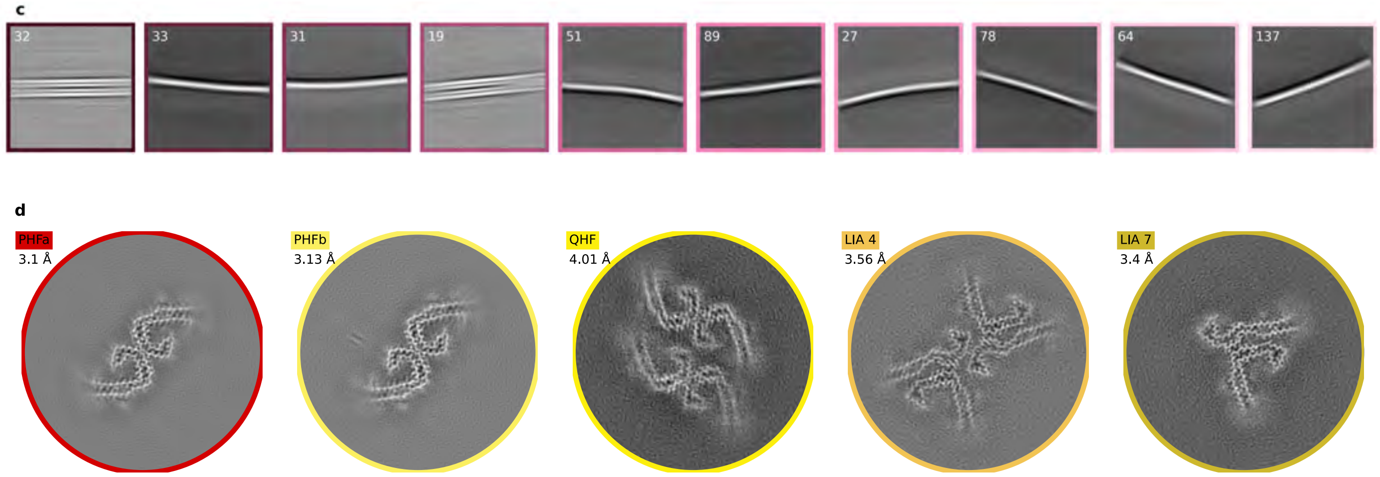

a

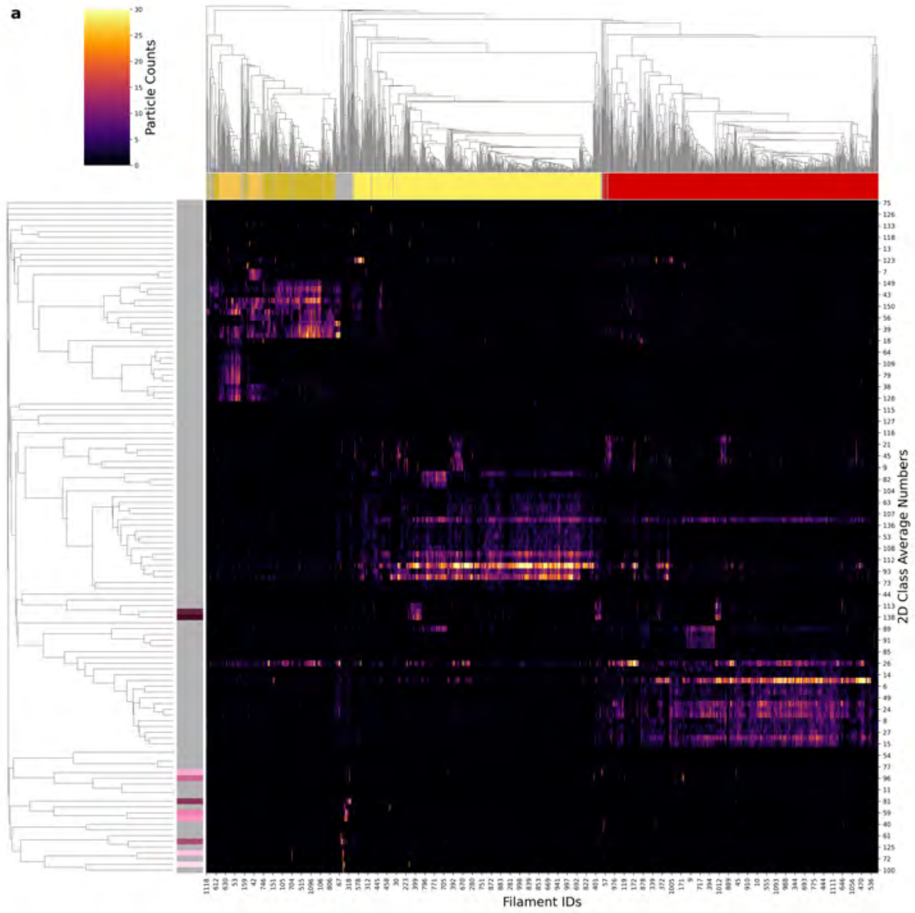

| Condition | PHF |
|---|---|
| Timepoint | 300 minutes |
| Replicate | 4 |
| Picking | Manual |
| Micrographs | 3923 |
| Extracted Particles | 100371 |
| Pixel Size | 0.824 Å |
| Classified Particles | 100371 |

b

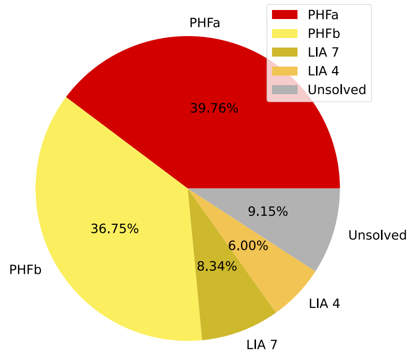

c

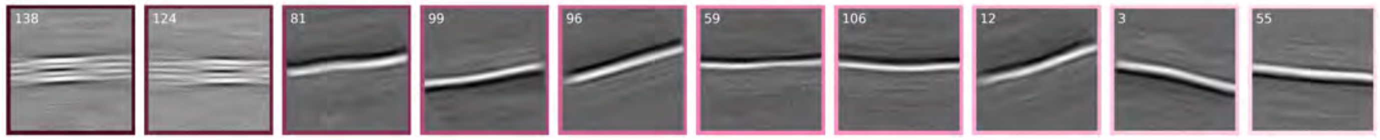

d

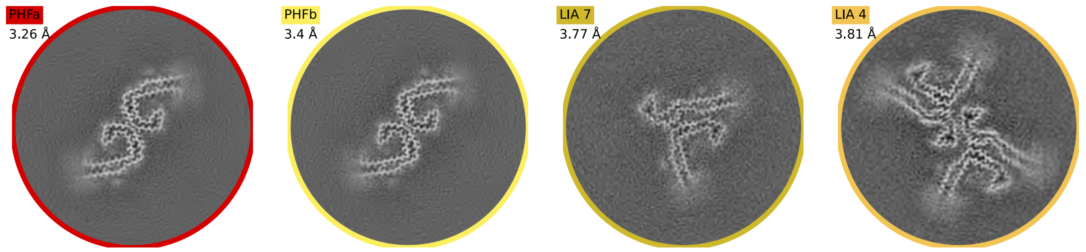

a

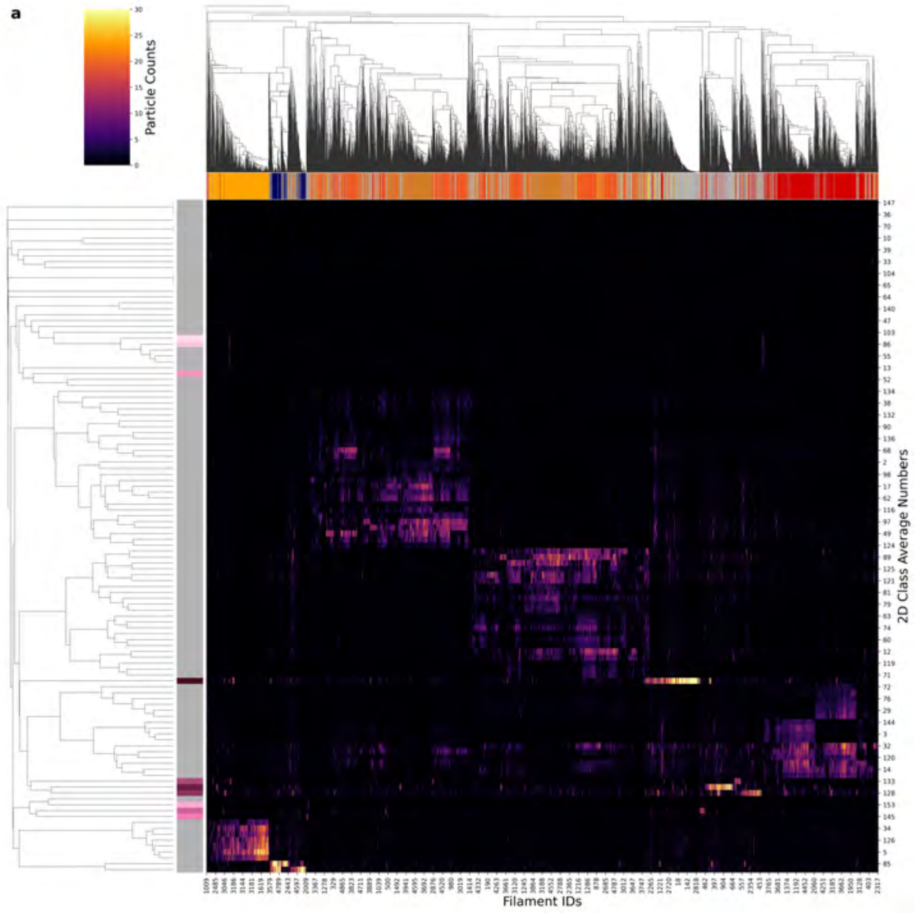

| Condition | PHF |
|---|---|
| Timepoint | 360 minutes |
| Replicate | 1 |
| Picking | Manual |
| Micrographs | 2562 |
| Extracted Particles | 337152 |
| Pixel Size | 0.727 Å |
| Classified Particles | 337152 |

b

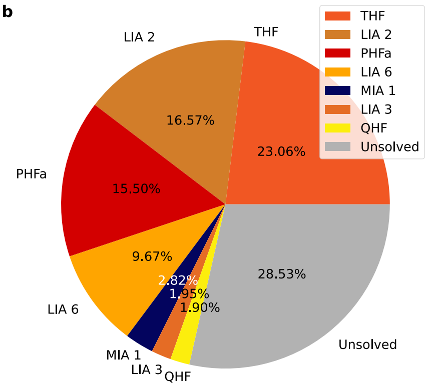

c

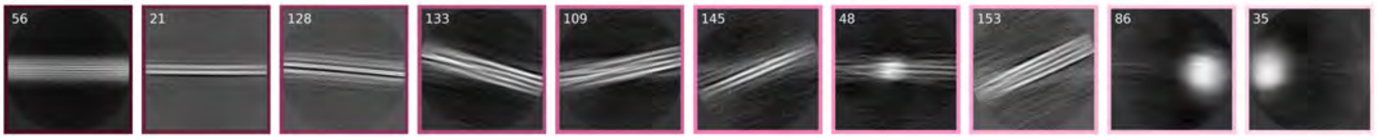

d

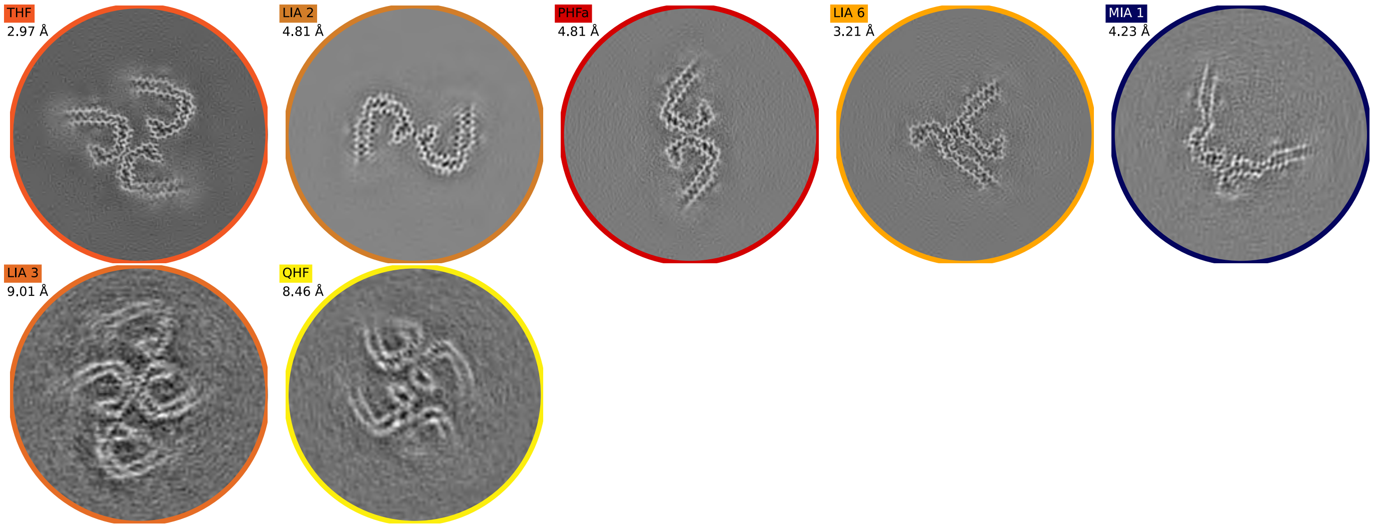

a

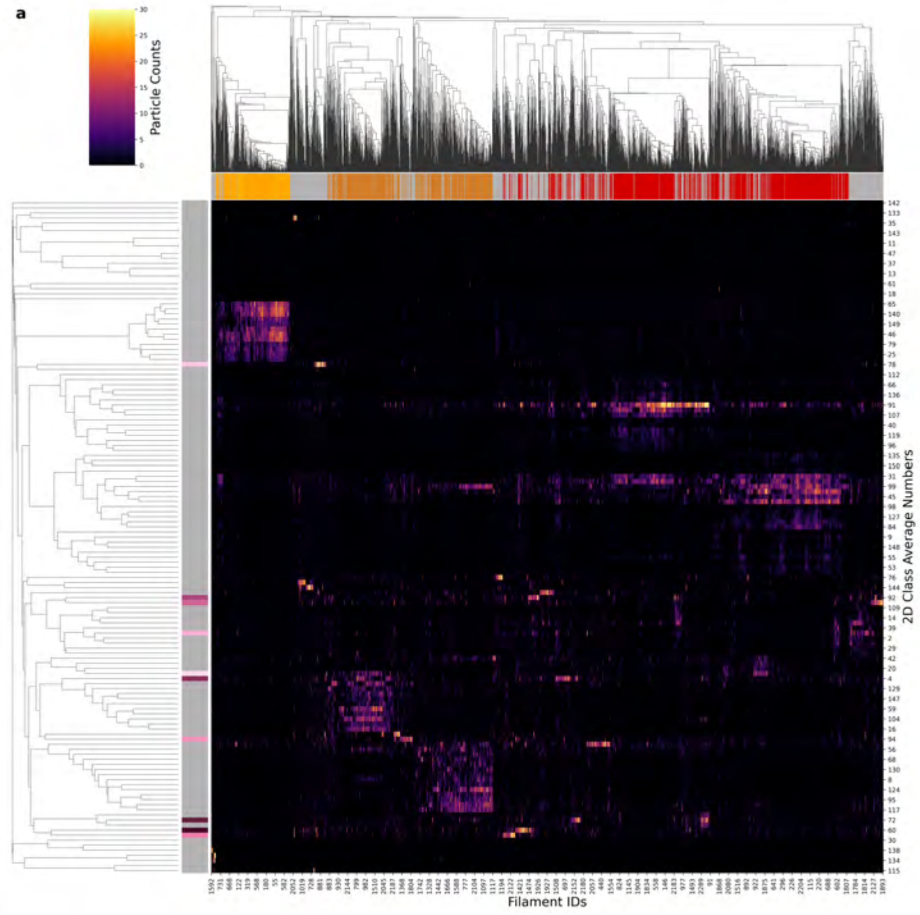

| Condition | PHF |
|---|---|
| Timepoint | 360 minutes |
| Replicate | 2 |
| Picking | Manual |
| Micrographs | 4519 |
| Extracted Particles | 178033 |
| Pixel Size | 0.824 Å |
| Classified Particles | 178033 |

b

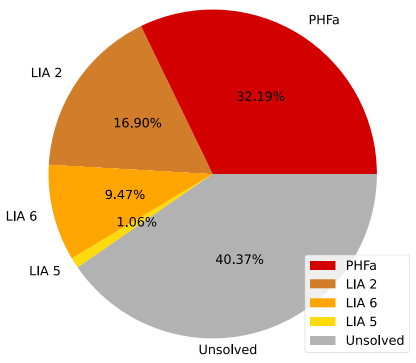

c

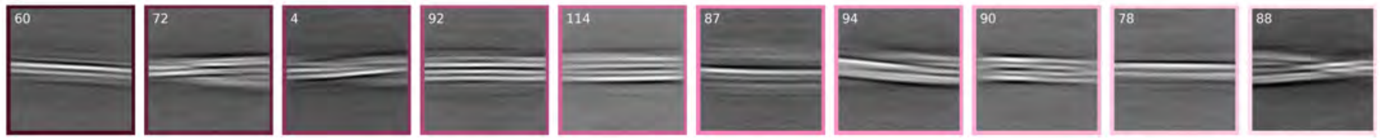

d

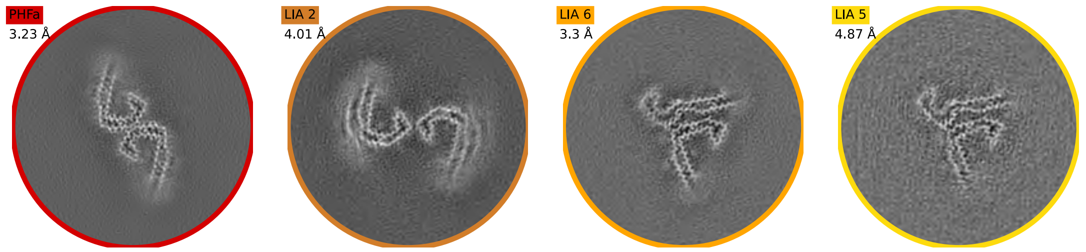

a

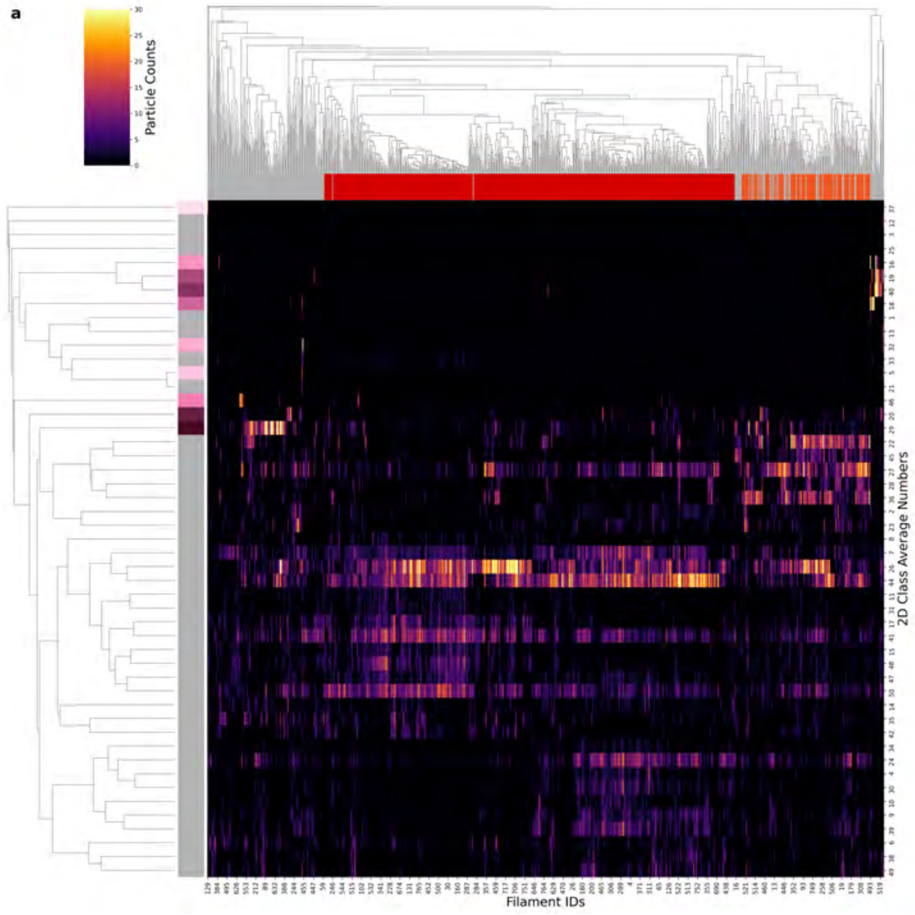

| Condition | PHF |
|---|---|
| Timepoint | 360 minutes |
| Replicate | 3 |
| Picking | Manual |
| Micrographs | 1063 |
| Extracted Particles | 54253 |
| Pixel Size | 0.824 Å |
| Classified Particles | 54253 |

b

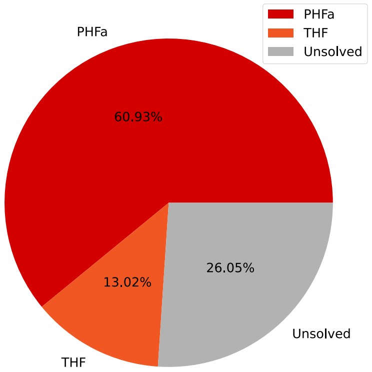

c

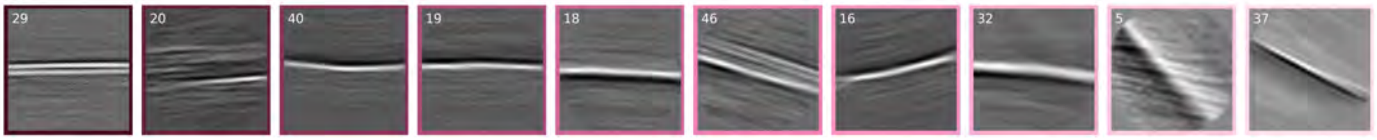

d

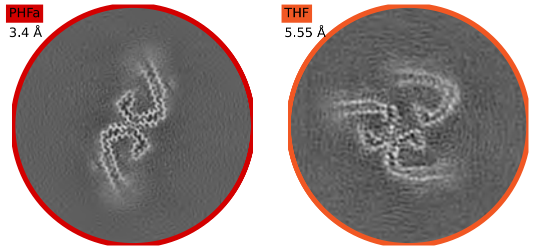

a

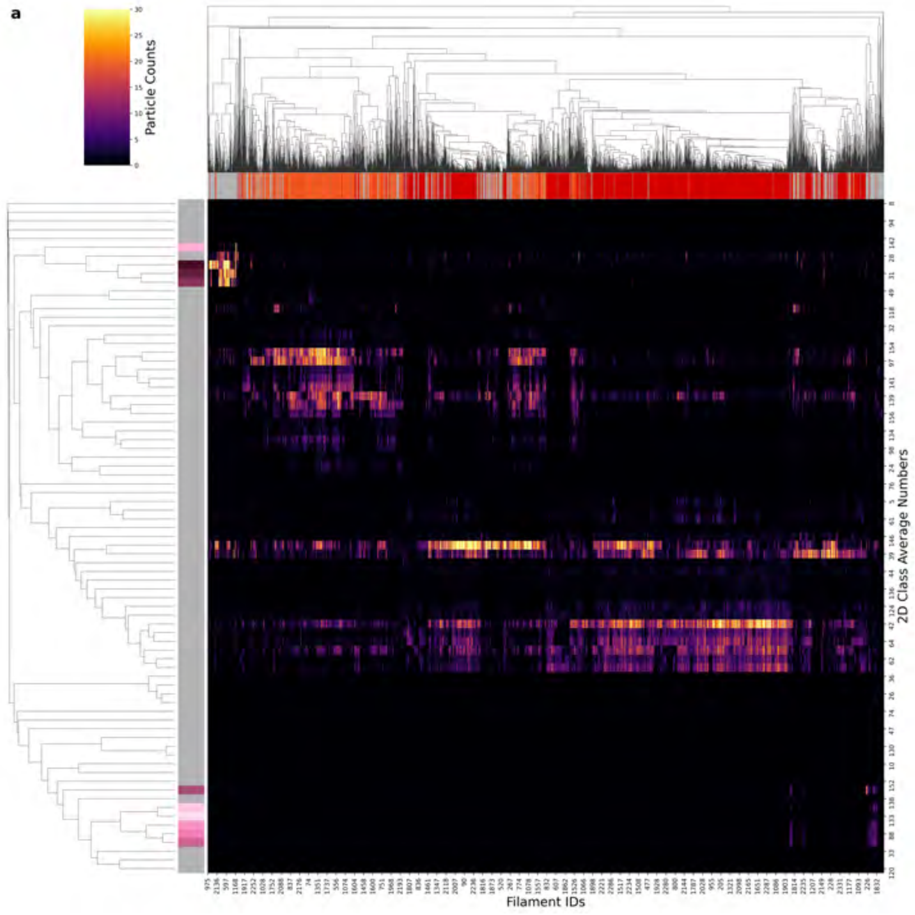

| Condition | PHF |
|---|---|
| Timepoint | 720 minutes |
| Replicate | 1 |
| Picking | Manual |
| Micrographs | 1105 |
| Extracted Particles | 179876 |
| Pixel Size | 0.824 Å |
| Classified Particles | 179876 |

b

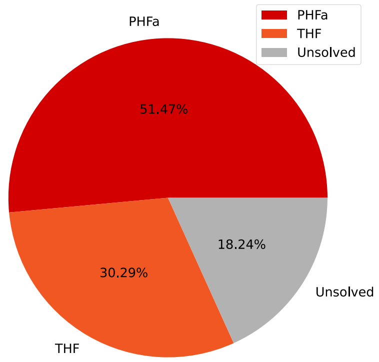

c

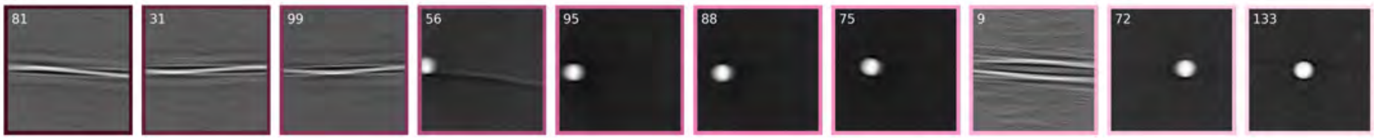

d

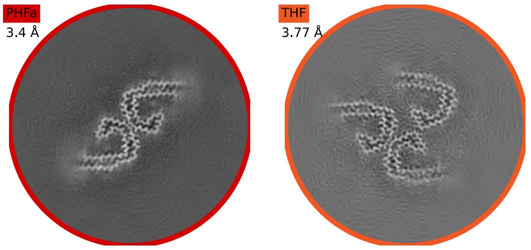

a

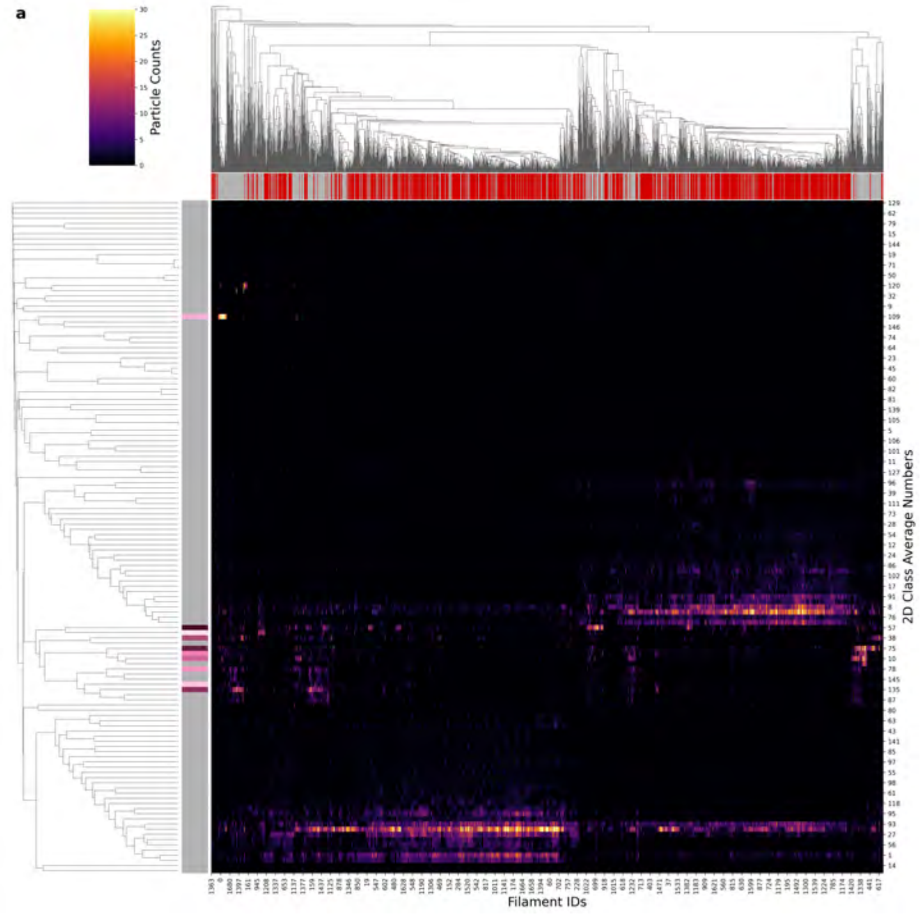

| | |
|---|---|
| **Condition** | PHF |
| **Timepoint** | 720 minutes |
| **Replicate** | 2 |
| **Picking** | Manual |
| **Micrographs** | 3379 |
| **Extracted Particles** | 125865 |
| **Pixel Size** | 0.727 Å |
| **Classified Particles** | 125865 |

b

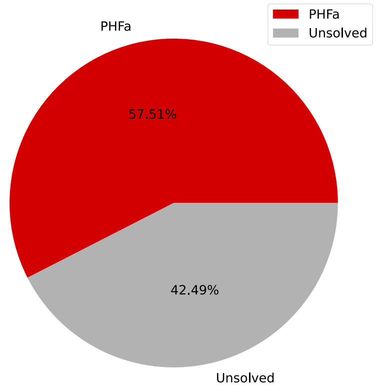

c

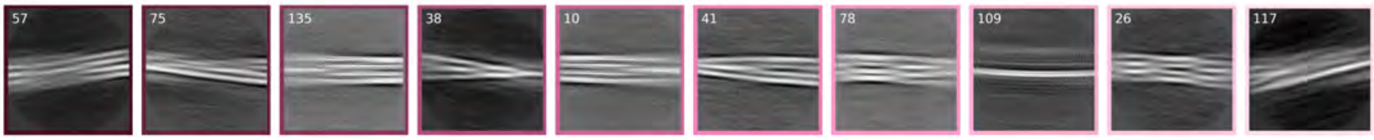

d

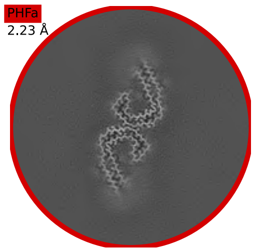

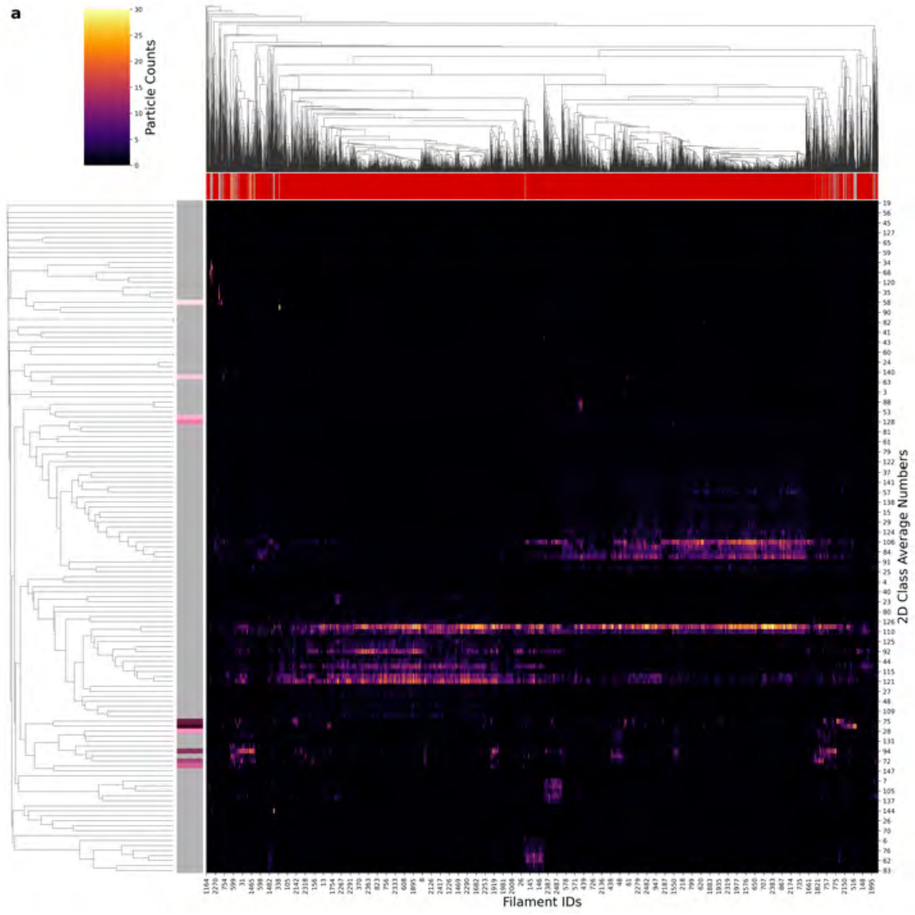

| | |
|---|---|
| **Condition** | PHF |
| **Timepoint** | 720 minutes |
| **Replicate** | 3 |
| **Picking** | Manual |
| **Micrographs** | 5267 |
| **Extracted Particles** | 175528 |
| **Pixel Size** | 0.824 Å |
| **Classified Particles** | 175528 |

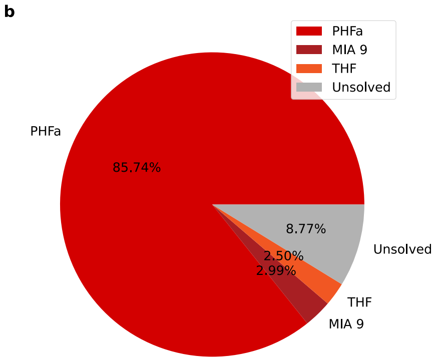

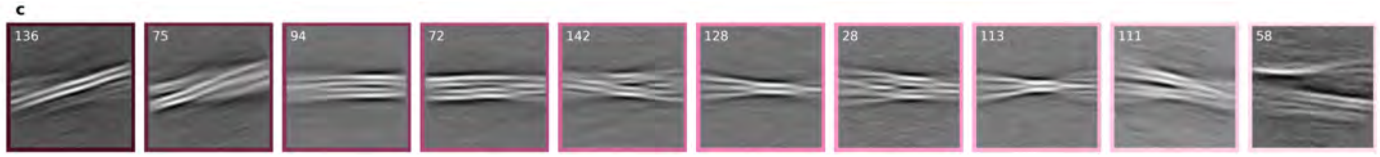

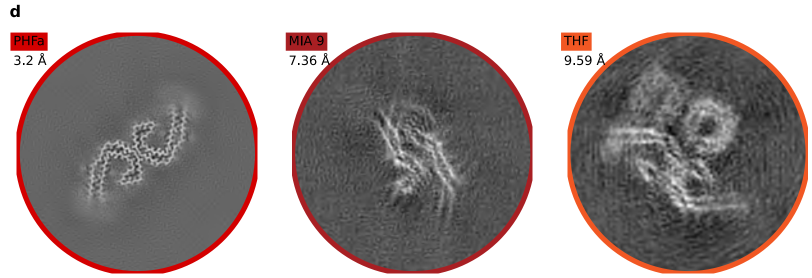

**a**

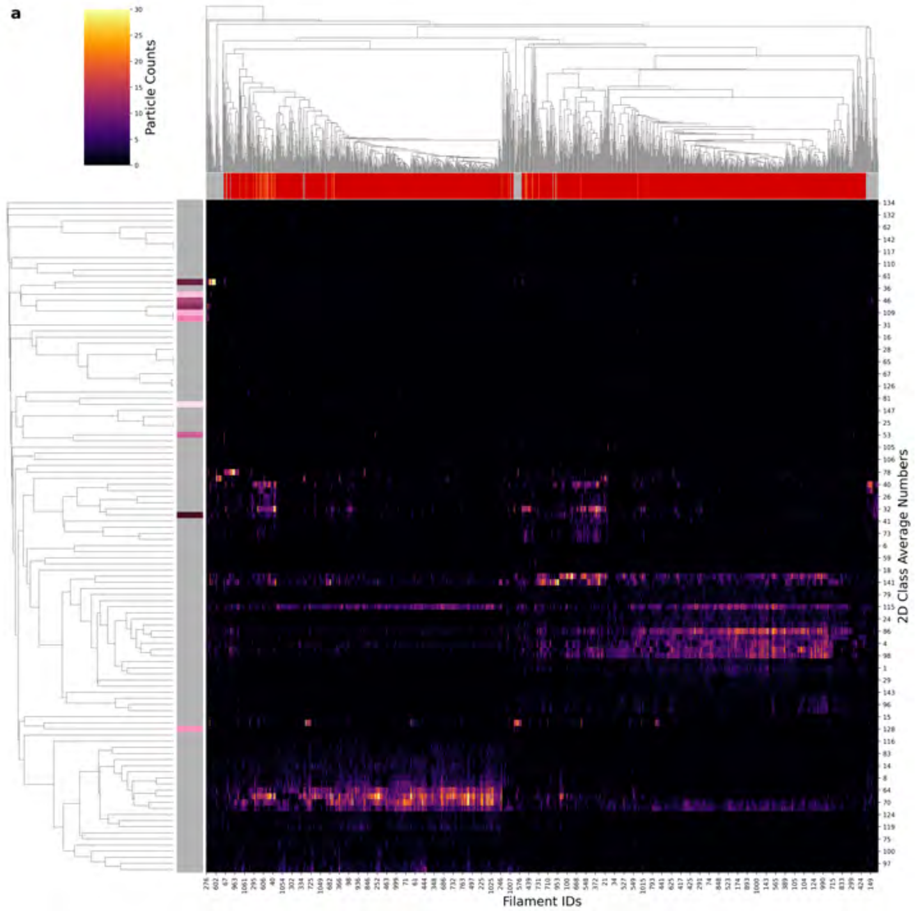

| Condition | PHF |
|---|---|
| **Timepoint** | 720 minutes |
| **Replicate** | 4 |
| **Picking** | Manual |
| **Micrographs** | 3379 |
| **Extracted Particles** | 77560 |
| **Pixel Size** | 0.727 Å |
| **Classified Particles** | 77560 |

**b**

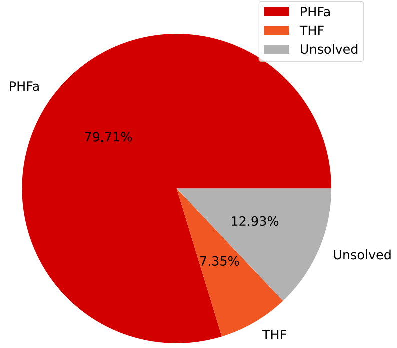

**c**

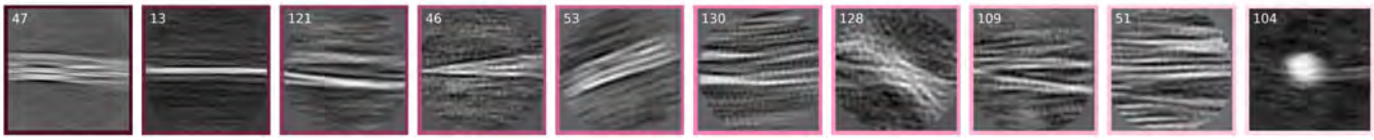

**d**

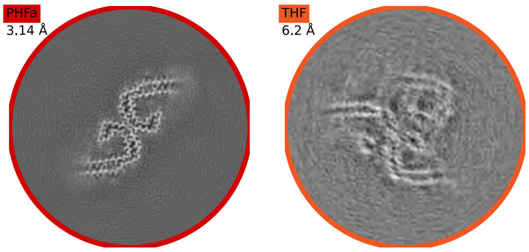

a

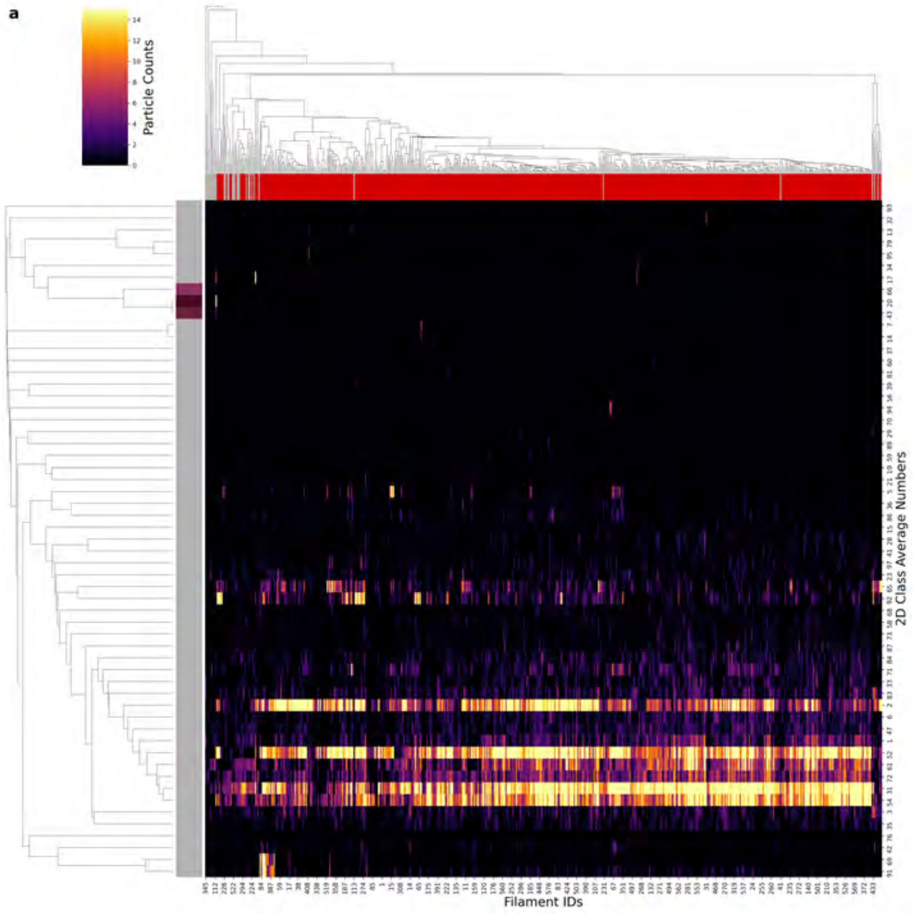

| | |
|---|---|
| **Condition** | AD |
| **Timepoint** | 720 minutes |
| **Replicate** | 5 |
| **Picking** | Manual |
| **Micrographs** | 1880 |
| **Extracted Particles** | 46371 |
| **Pixel Size** | 0.824 Å |
| **Classified Particles** | 46371 |

b

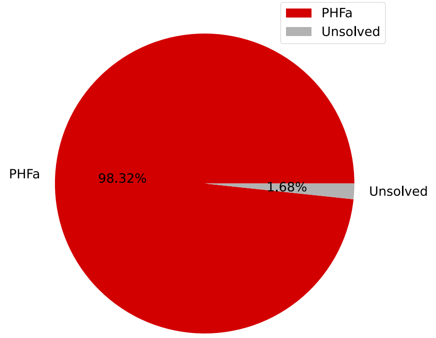

c

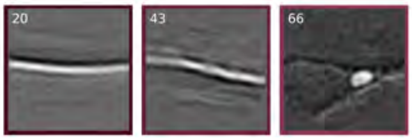

d

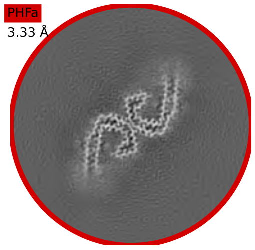

a

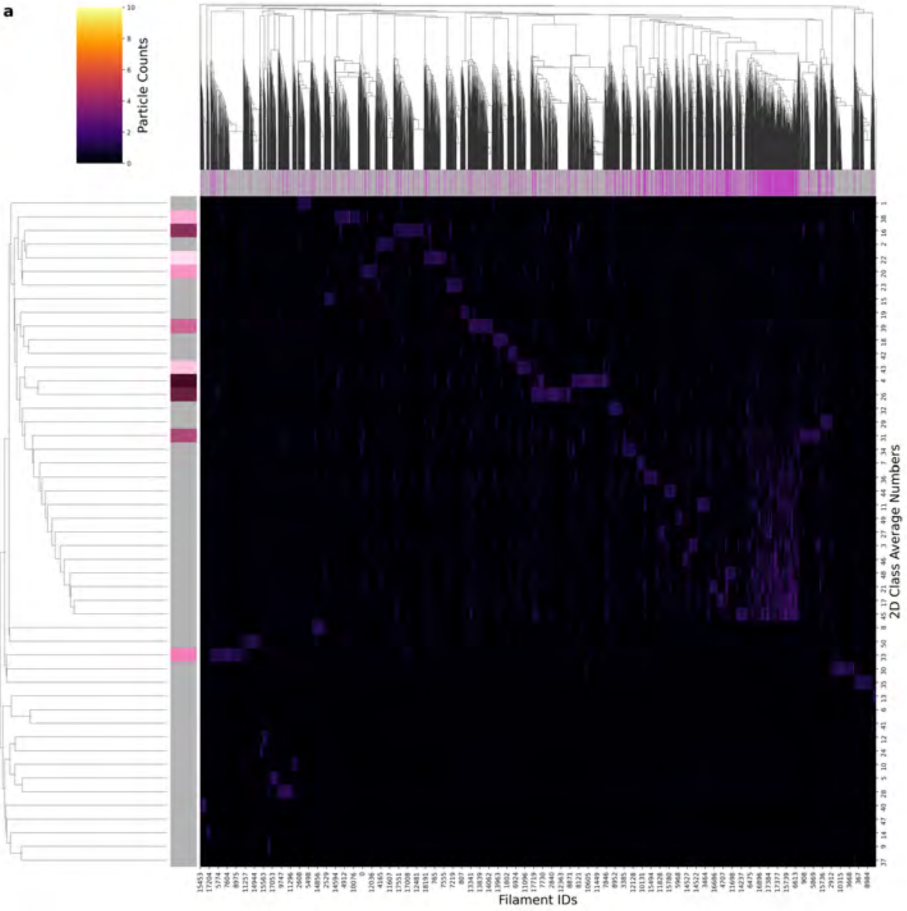

| Condition | CTE |
| --- | --- |
| Timepoint | 120 minutes |
| Replicate | 1 |
| Picking | Auto |
| Micrographs | 1727 |
| Extracted Particles | 330525 |
| Pixel Size | 0.672 Å |
| Classified Particles | 82995 |

b

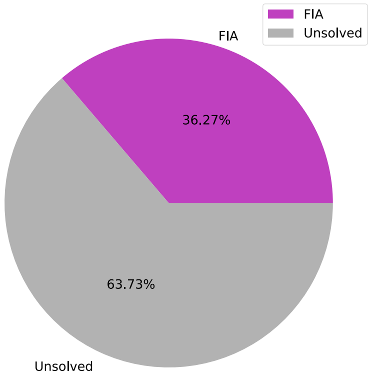

c

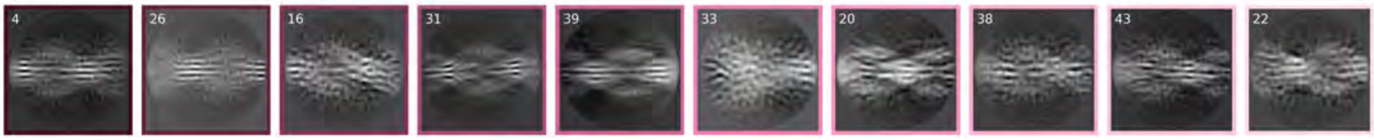

d

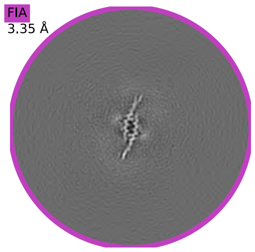

**a**

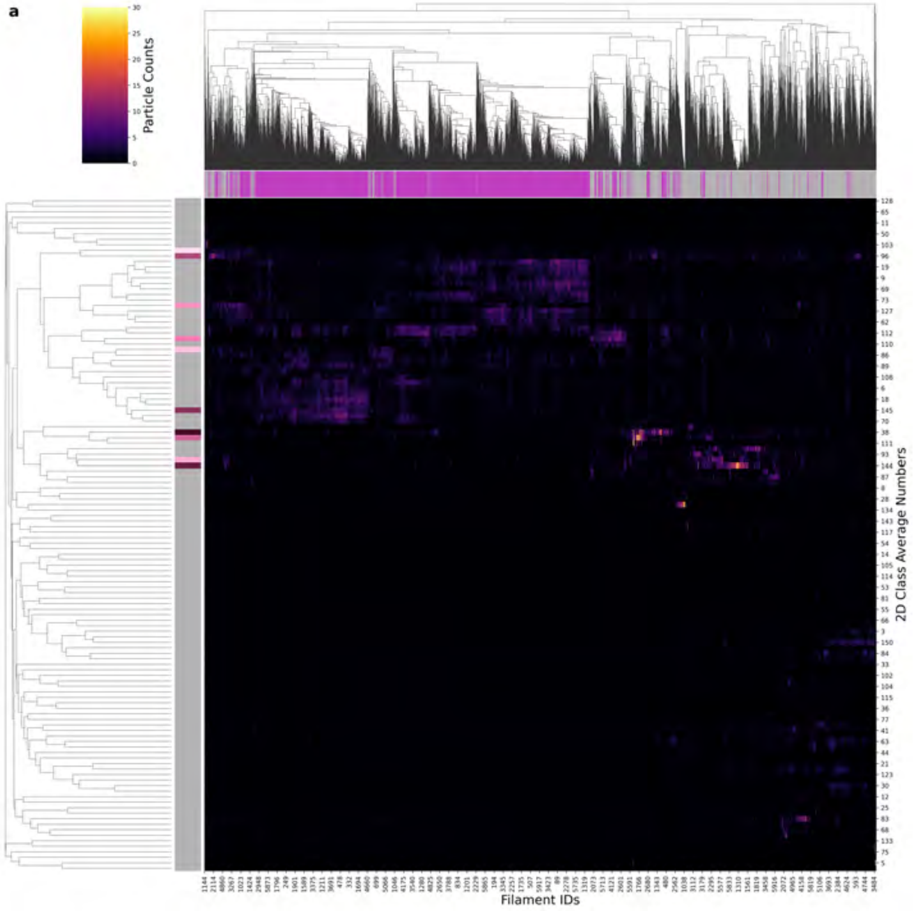

| Condition | CTE |
|---|---|
| **Timepoint** | 120 minutes |
| **Replicate** | 2 |
| **Picking** | Auto |
| **Micrographs** | 3948 |
| **Extracted Particles** | 221378 |
| **Pixel Size** | 0.727 Å |
| **Classified Particles** | 221378 |

**b**

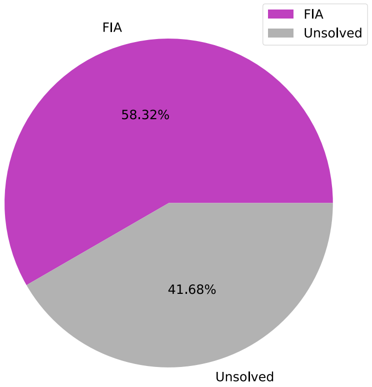

**c**

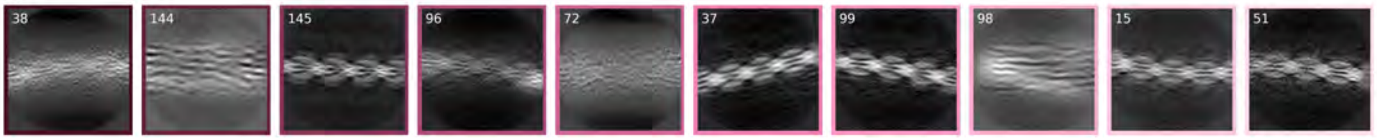

**d**

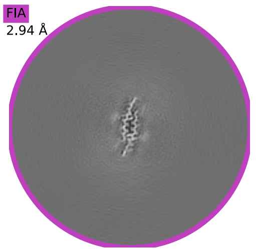

**a**

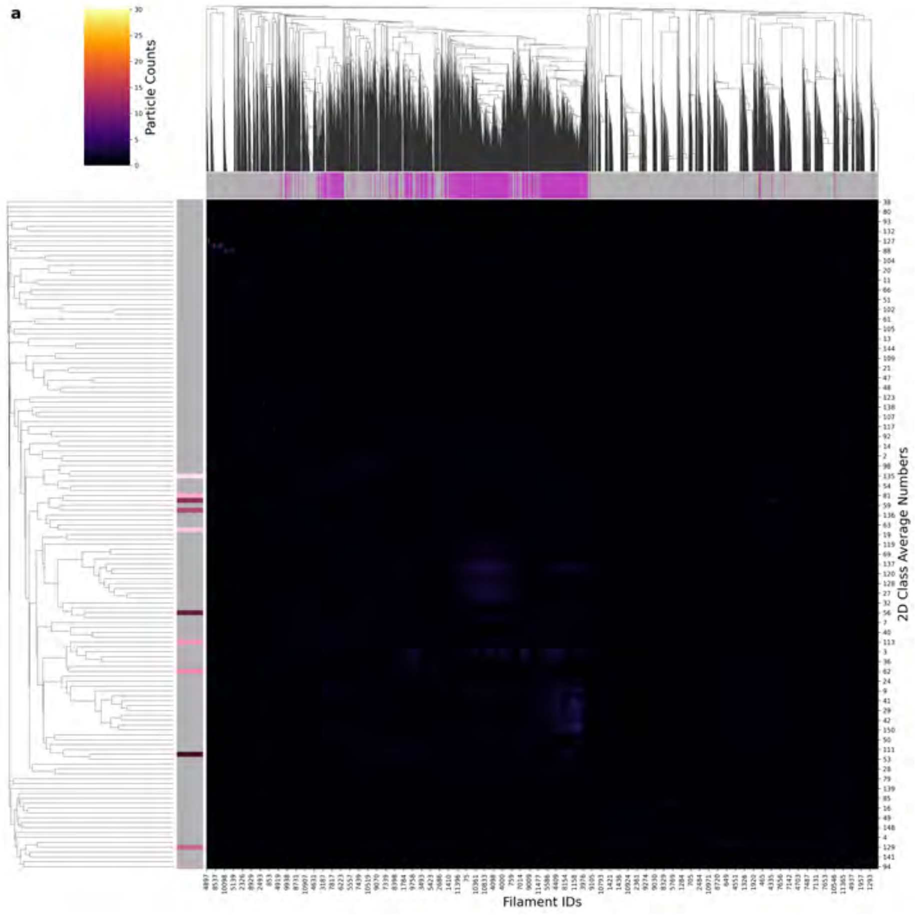

| Condition | CTE |
|---|---|
| **Timepoint** | 120 minutes |
| **Replicate** | 4 |
| **Picking** | Autopick |
| **Micrographs** | 3940 |
| **Extracted Particles** | 127070 |
| **Pixel Size** | 0.727 Å |
| **Classified Particles** | 127070 |

**b**

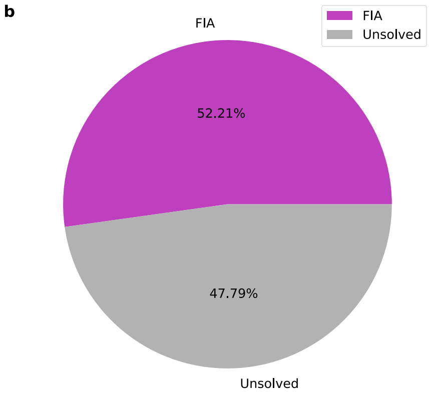

**c**

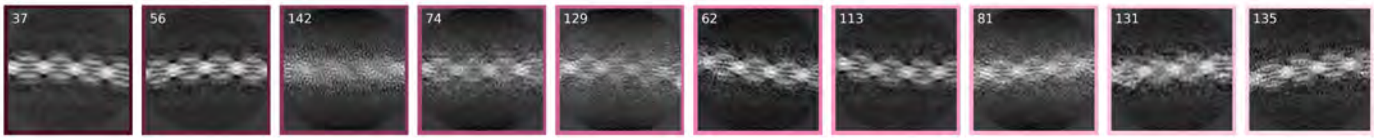

**d**

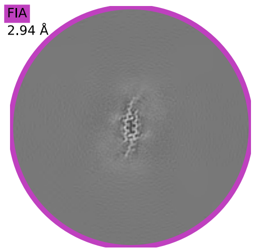

a

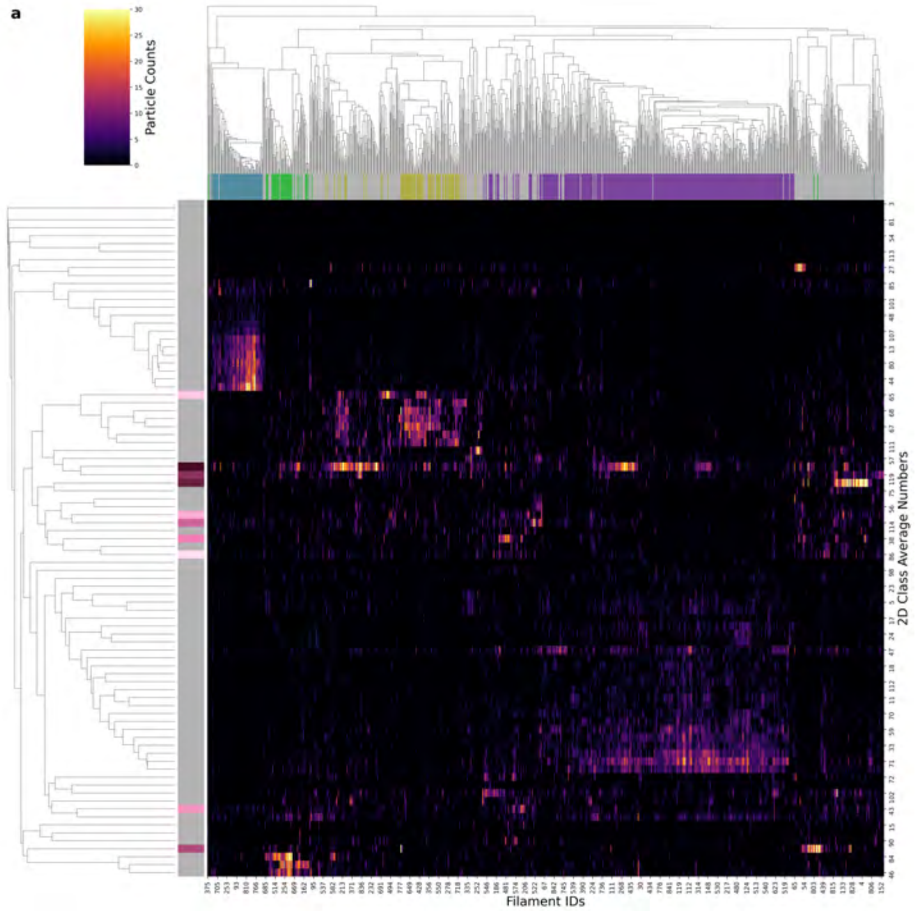

| | |
|---|---|
| **Condition** | CTE |
| **Timepoint** | 160 minutes |
| **Replicate** | 4 |
| **Picking** | Manual |
| **Micrographs** | 932 |
| **Extracted Particles** | 59766 |
| **Pixel Size** | 0.824 Å |
| **Classified Particles** | 59766 |

b

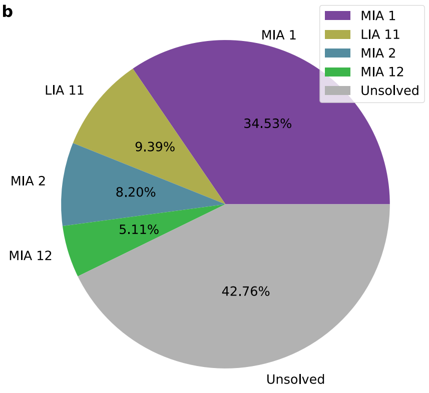

c

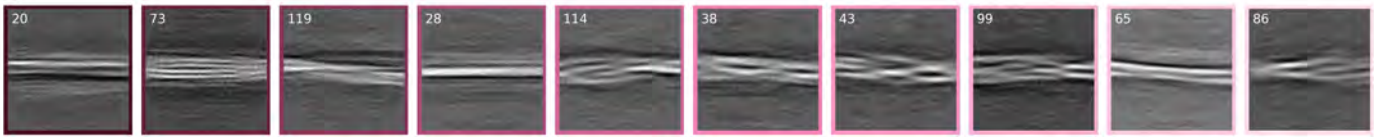

d

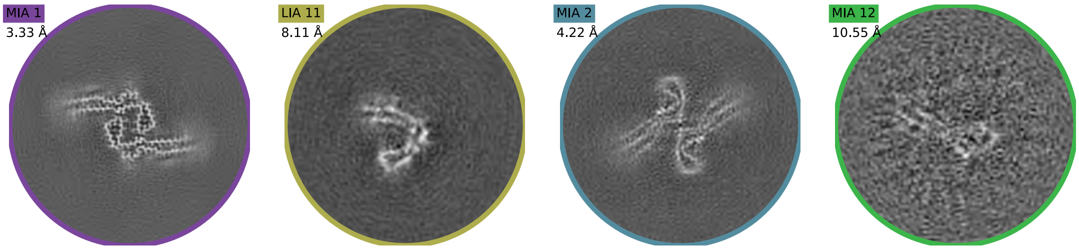

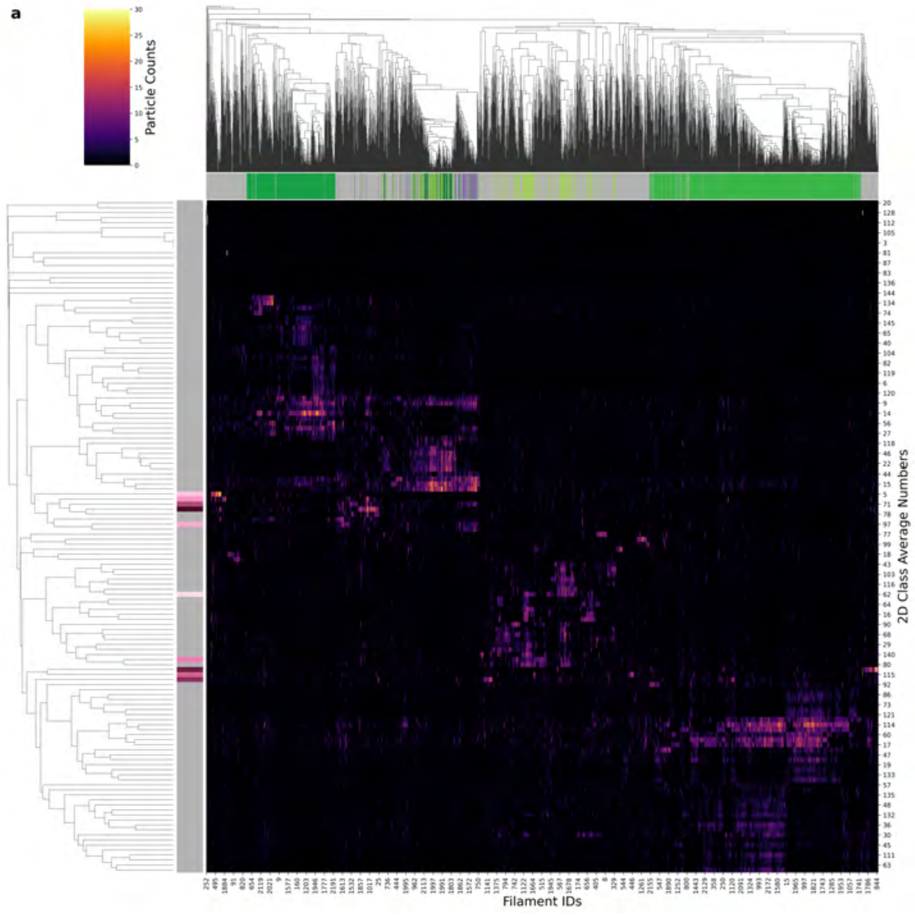

| Condition | CTE |
|---|---|
| Timepoint | 160 minutes |
| Replicate | 5 |
| Picking | Manual |
| Micrographs | 2067 |
| Extracted Particles | 158080 |
| Pixel Size | 0.824 Å |
| Classified Particles | 158080 |

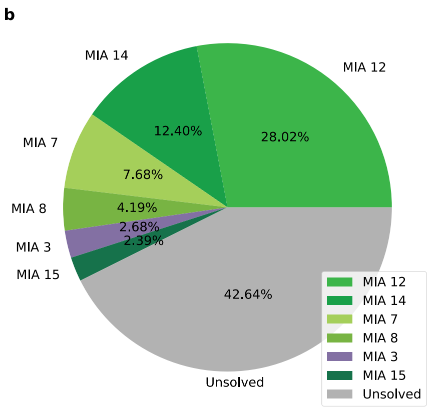

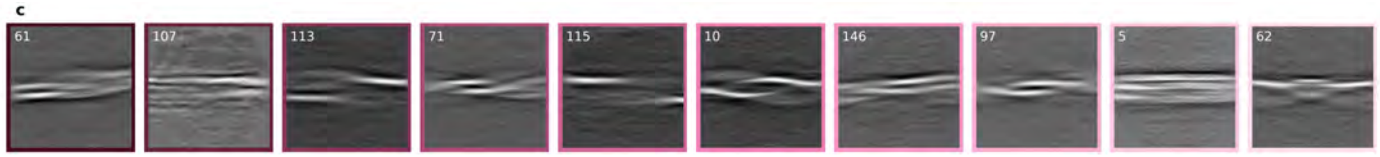

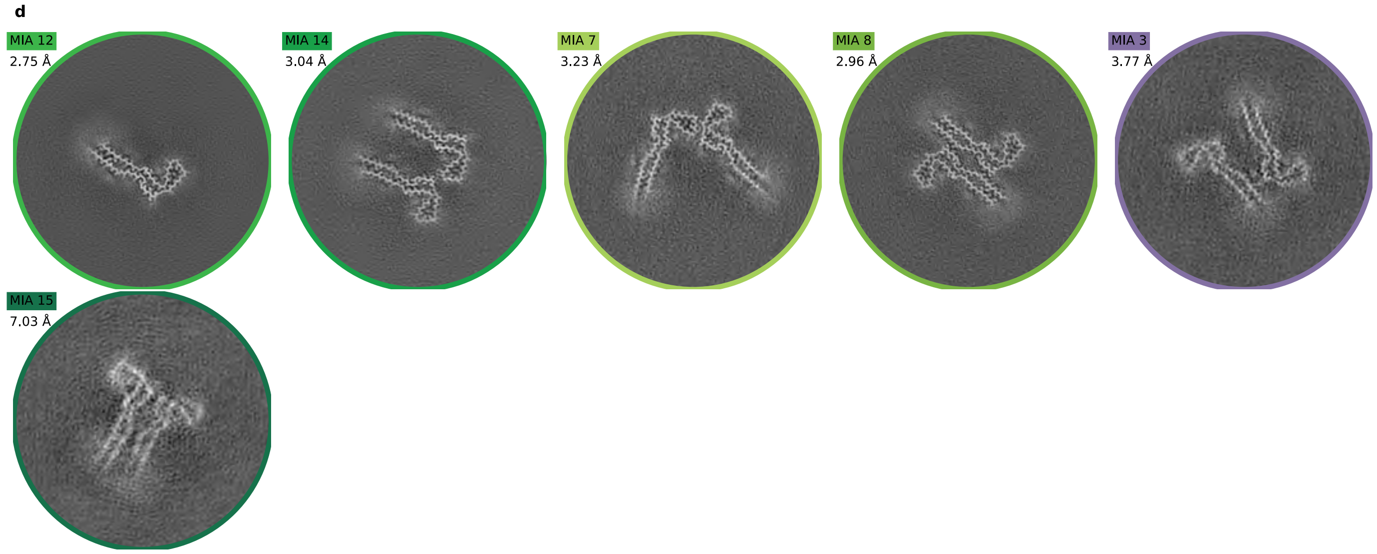

a

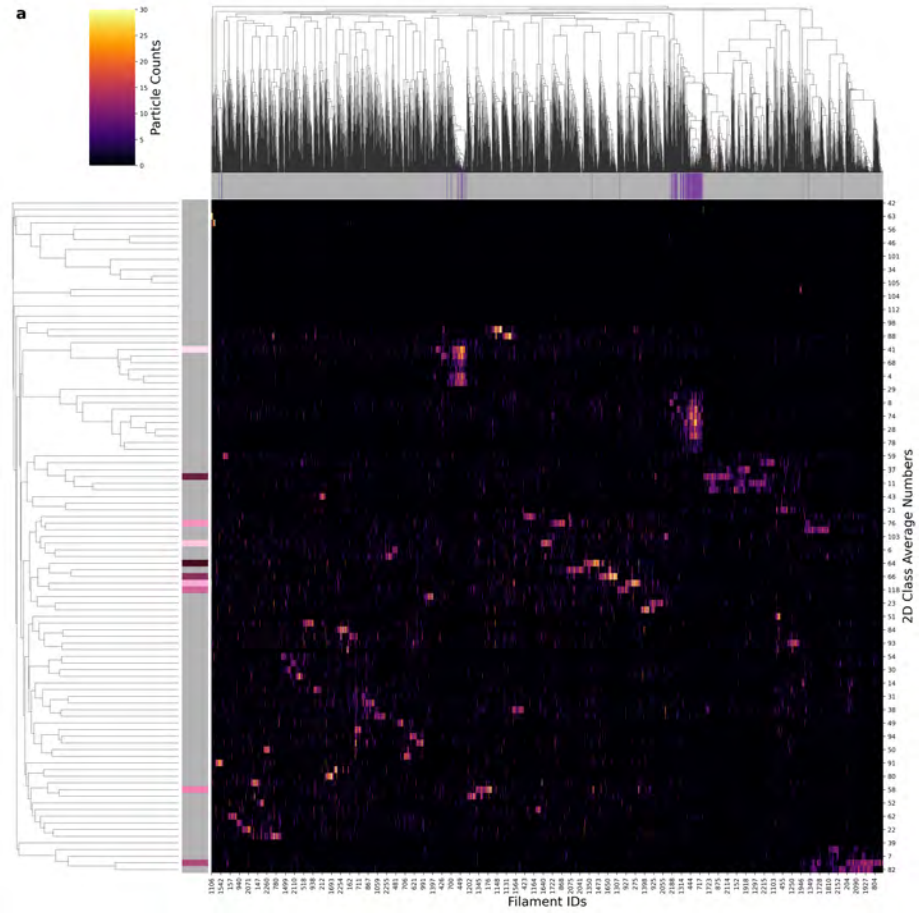

| Condition | CTE |
| --- | --- |
| **Timepoint** | 160 minutes |
| **Replicate** | 6 |
| **Picking** | Manual |
| **Micrographs** | 5427 |
| **Extracted Particles** | 103098 |
| **Pixel Size** | 0.824 Å |
| **Classified Particles** | 103098 |

b

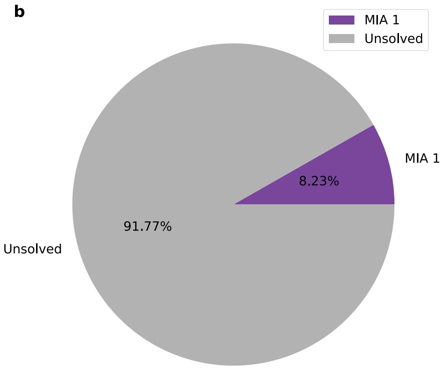

c

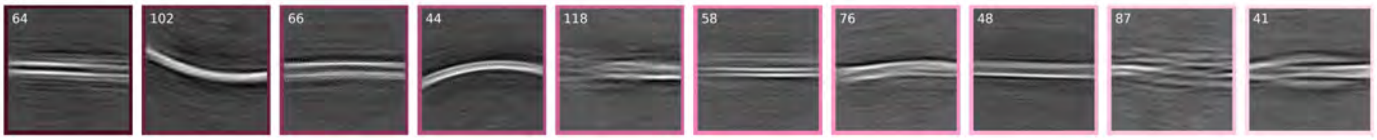

d

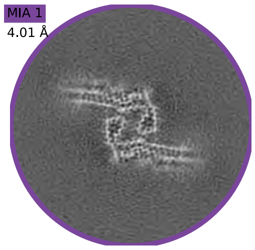

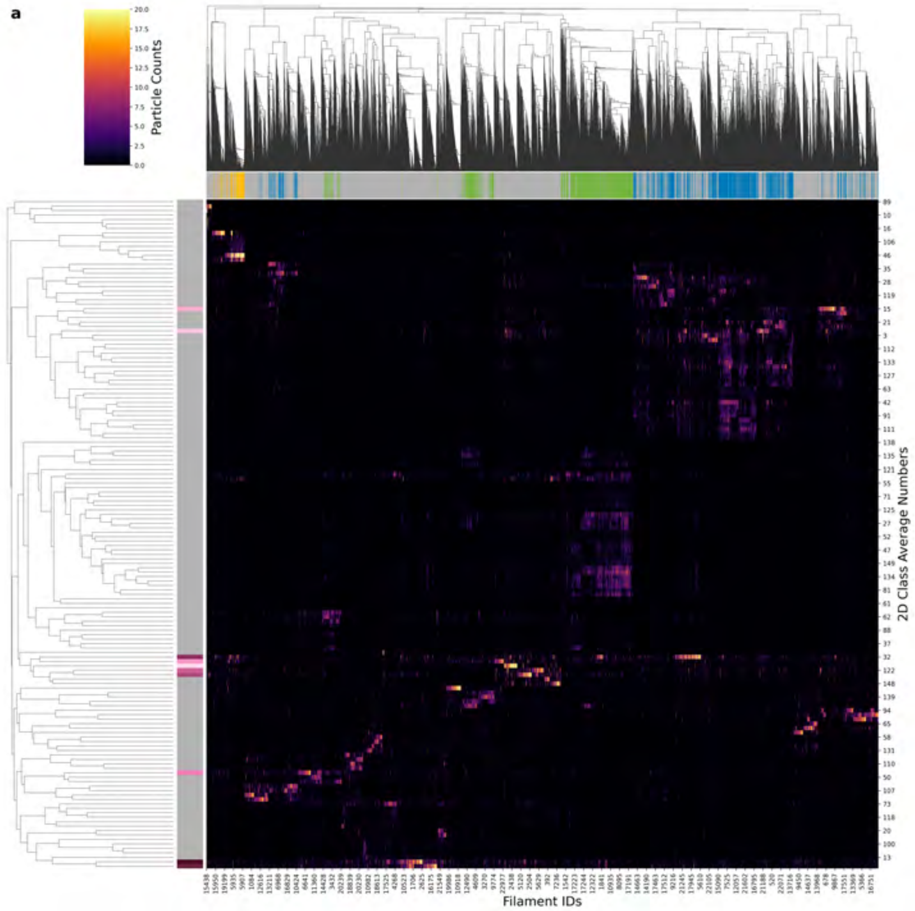

| Condition | CTE |
|---|---|
| Timepoint | 180 minutes |
| Replicate | 1 |
| Picking | Auto |
| Micrographs | 2514 |
| Extracted Particles | 960020 |
| Pixel Size | 0.727 Å |
| Classified Particles | 960020 |

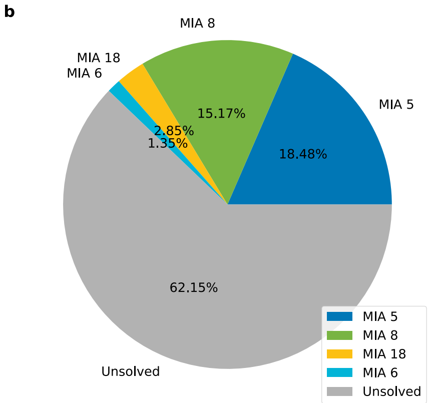

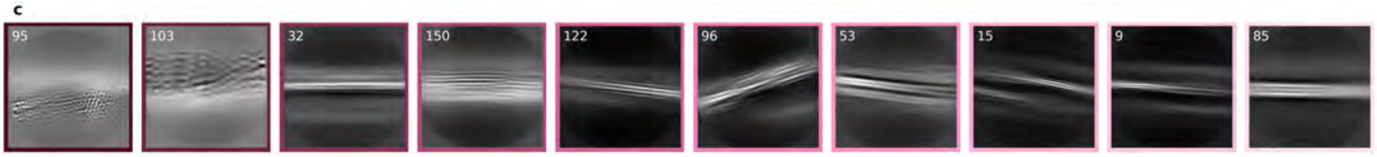

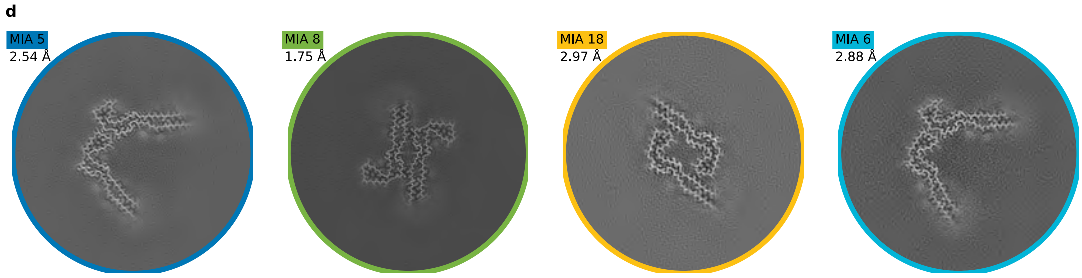

a

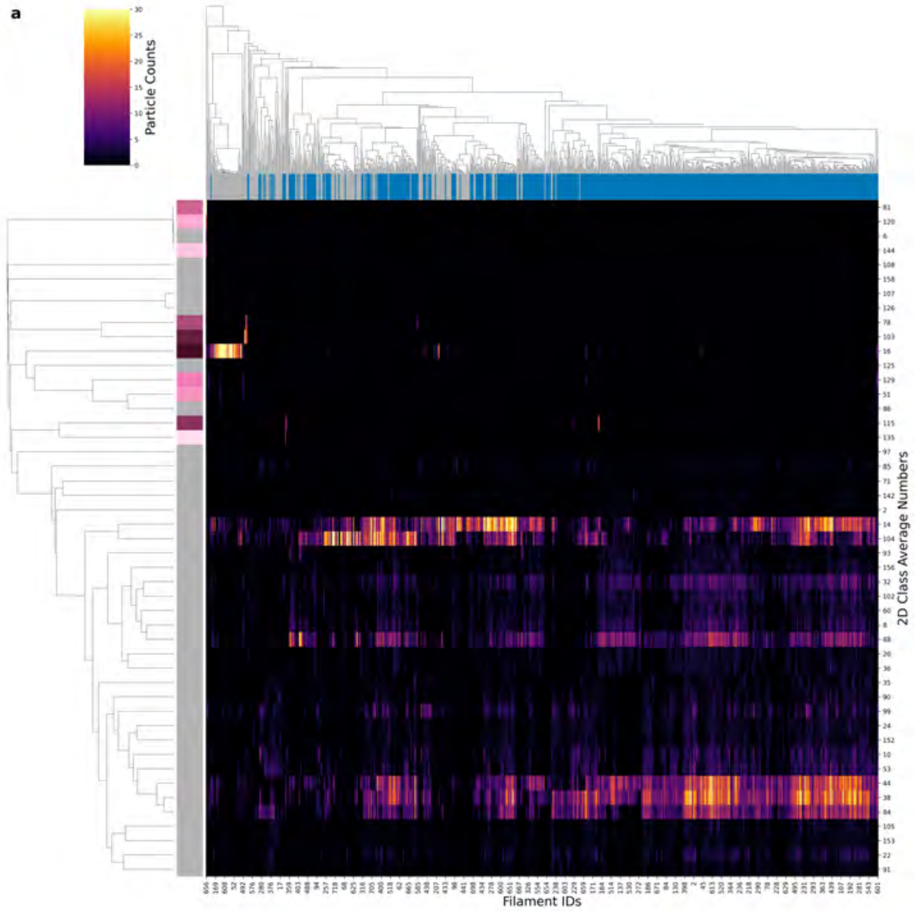

| | |
|---|---|
| **Condition** | CTE |
| **Timepoint** | 180 minutes |
| **Replicate** | 2 |
| **Picking** | Manual |
| **Micrographs** | 223 |
| **Extracted Particles** | 47641 |
| **Pixel Size** | 0.824 Å |
| **Classified Particles** | 47641 |

b

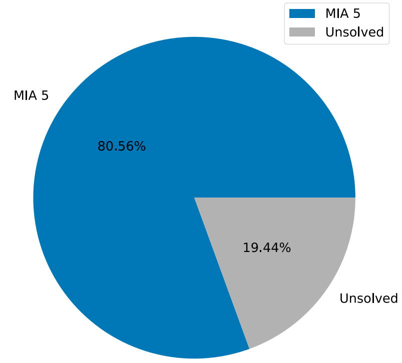

c

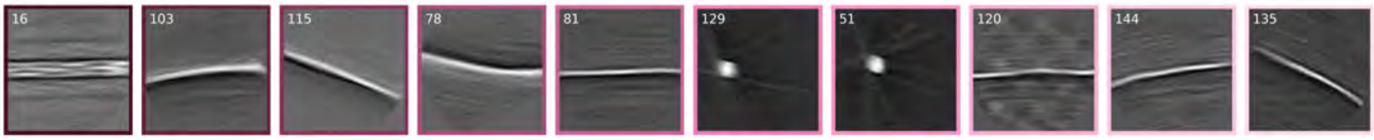

d

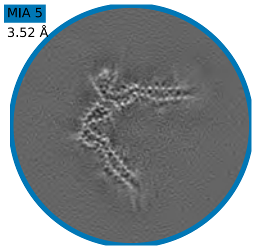

a

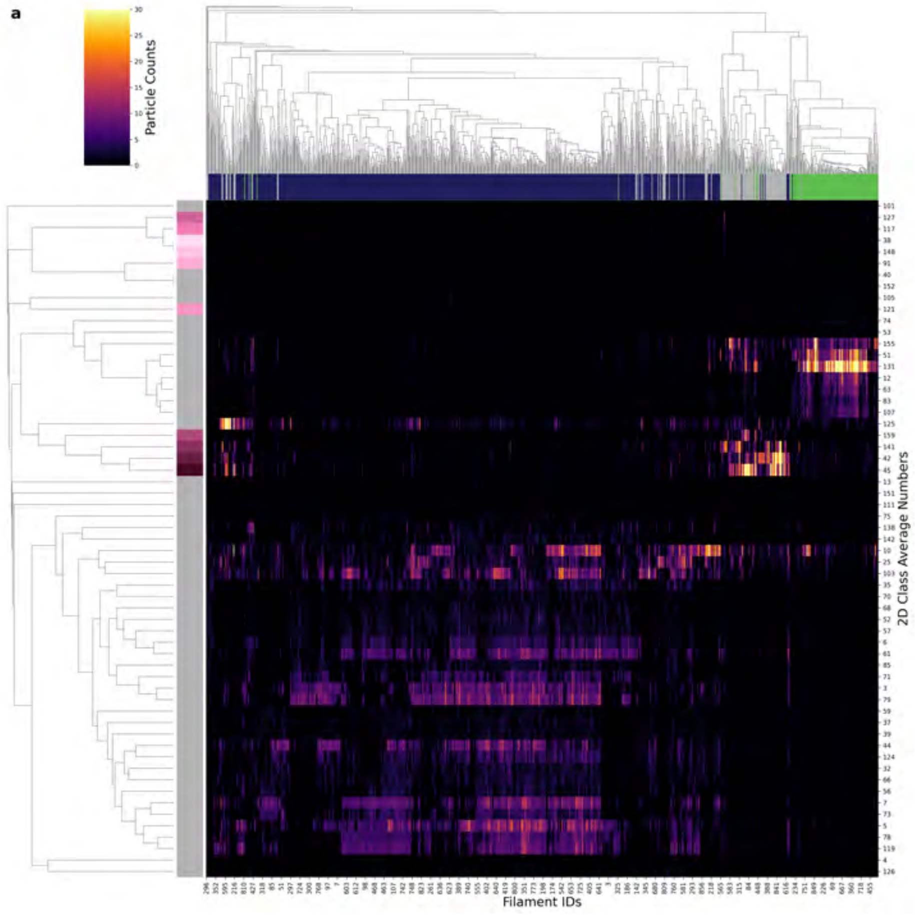

| Condition | CTE |
|---|---|
| Timepoint | 180 minutes |
| Replicate | 4 |
| Picking | Manual |
| Micrographs | 1782 |
| Extracted Particles | 58123 |
| Pixel Size | 0.824 Å |
| Classified Particles | 58123 |

b

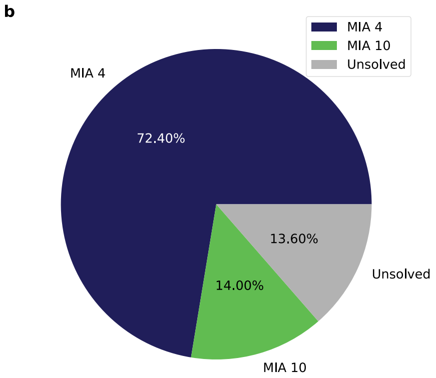

c

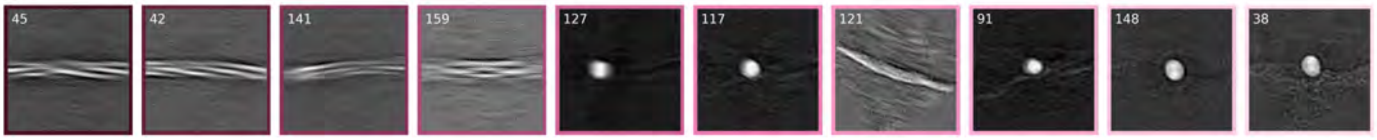

d

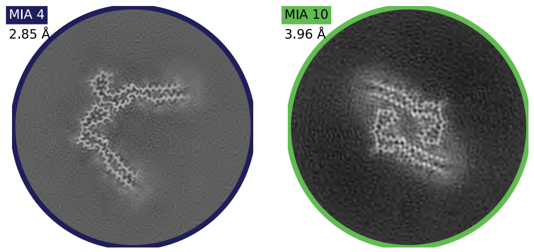

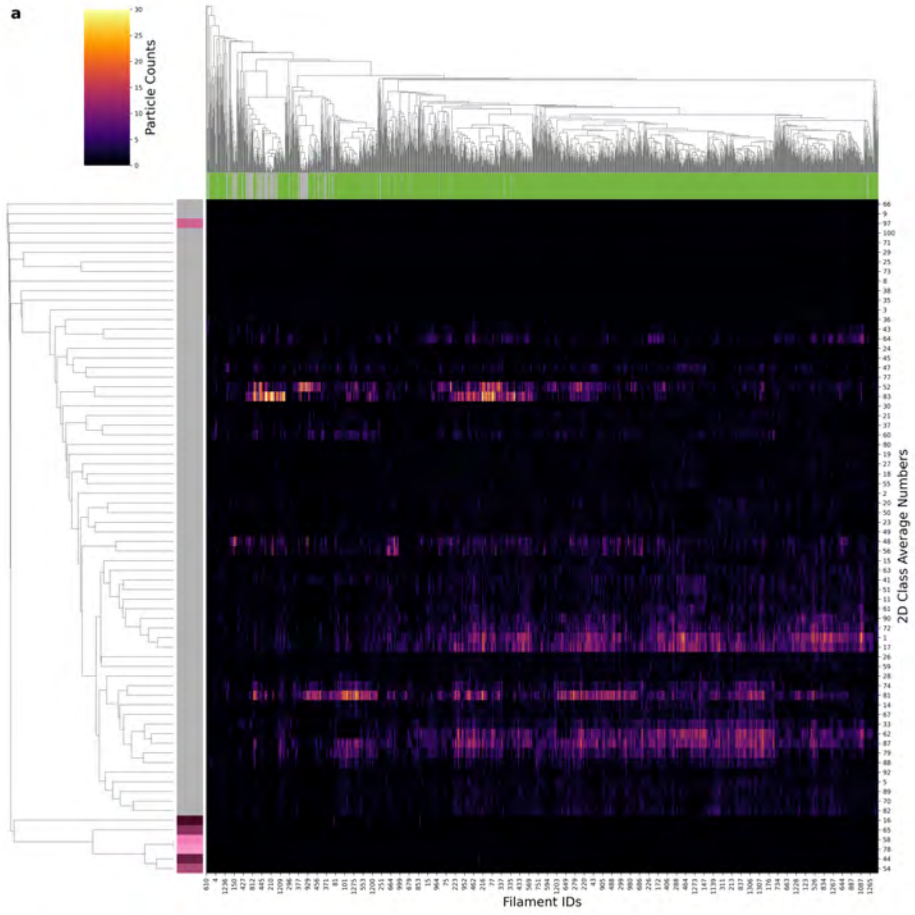

| Condition | CTE |
|---|---|
| Timepoint | 180 minutes |
| Replicate | 5 |
| Picking | Manual |
| Micrographs | 1385 |
| Extracted Particles | 99429 |
| Pixel Size | 0.824 Å |
| Classified Particles | 99429 |

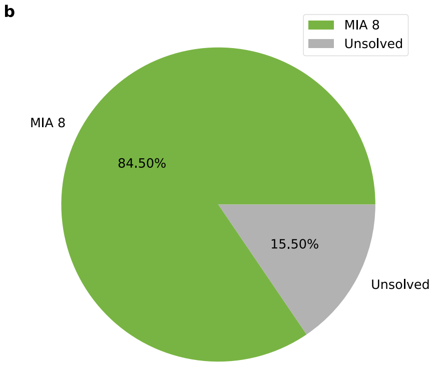

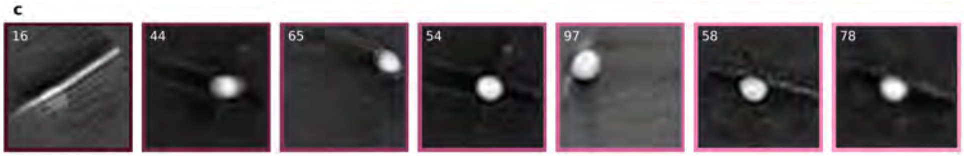

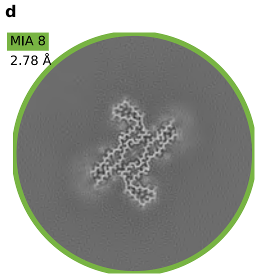

a

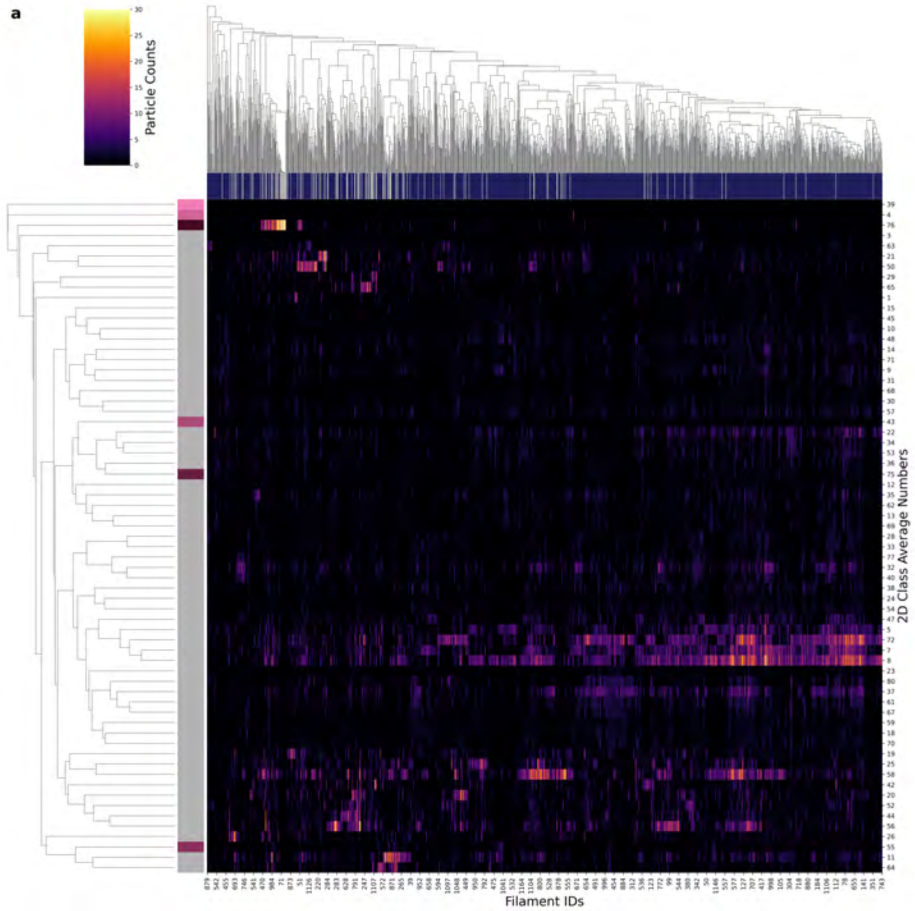

| Condition | CTE |
|---|---|
| Timepoint | 180 minutes |
| Replicate | 6 |
| Picking | Manual |
| Micrographs | 601 |
| Extracted Particles | 71046 |
| Pixel Size | 0.824 Å |
| Classified Particles | 71046 |

b

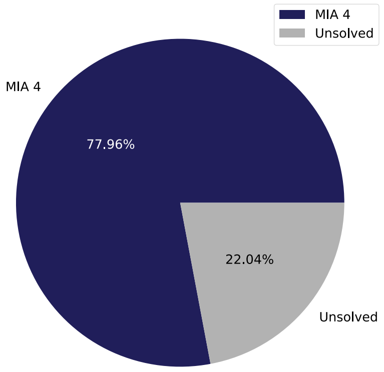

c

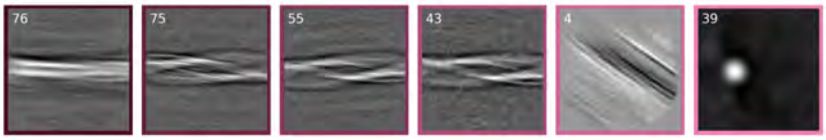

d

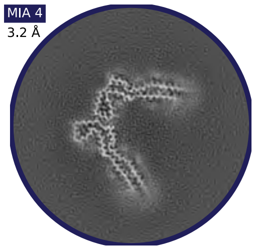

a

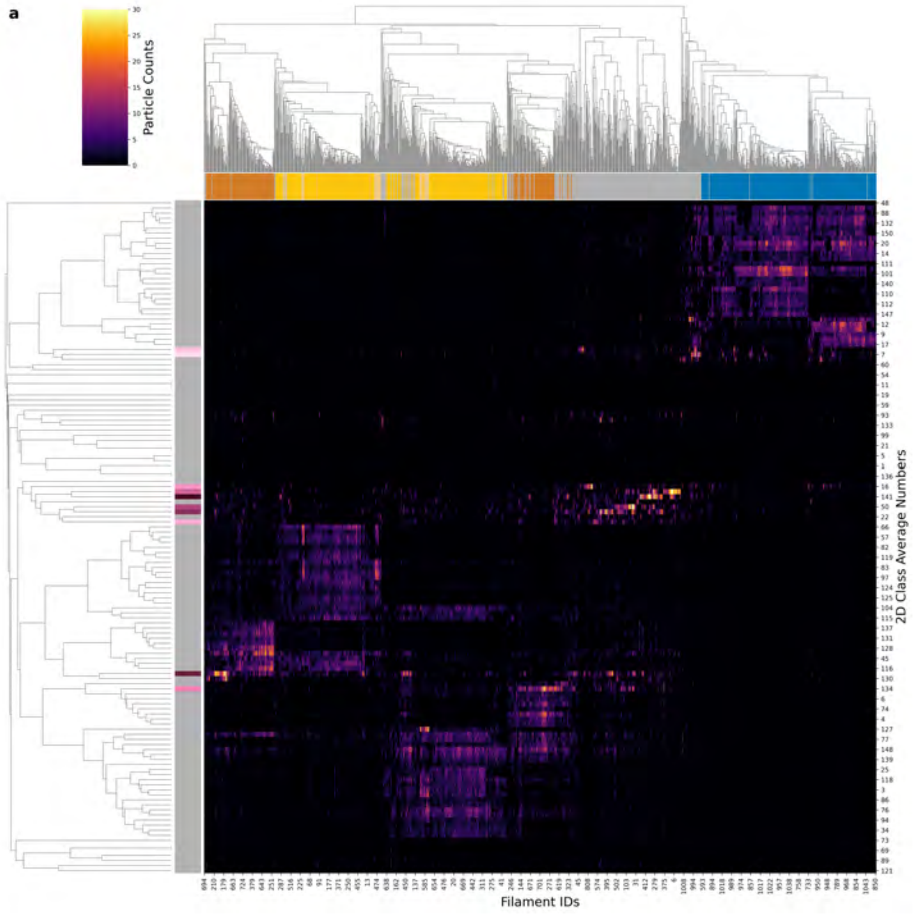

| Condition | CTE |
|---|---|
| Timepoint | 240 minutes |
| Replicate | 1 |
| Picking | Manual |
| Micrographs | 3902 |
| Extracted Particles | 95858 |
| Pixel Size | 0.727 Å |
| Classified Particles | 95858 |

b

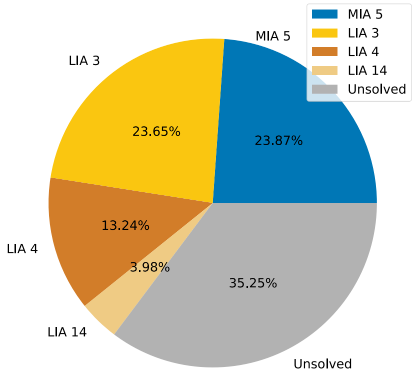

c

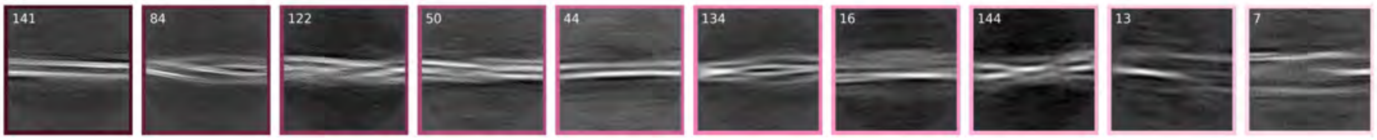

d

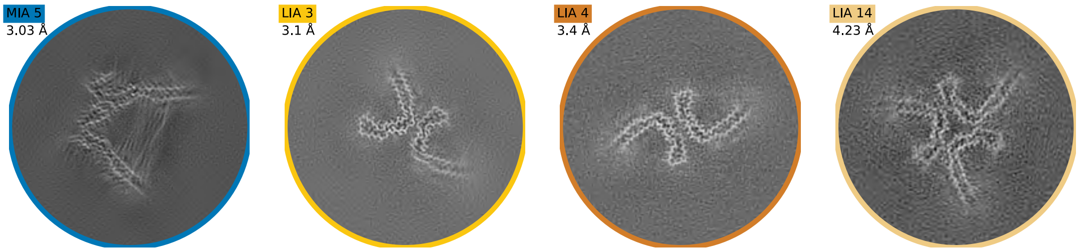

**a**

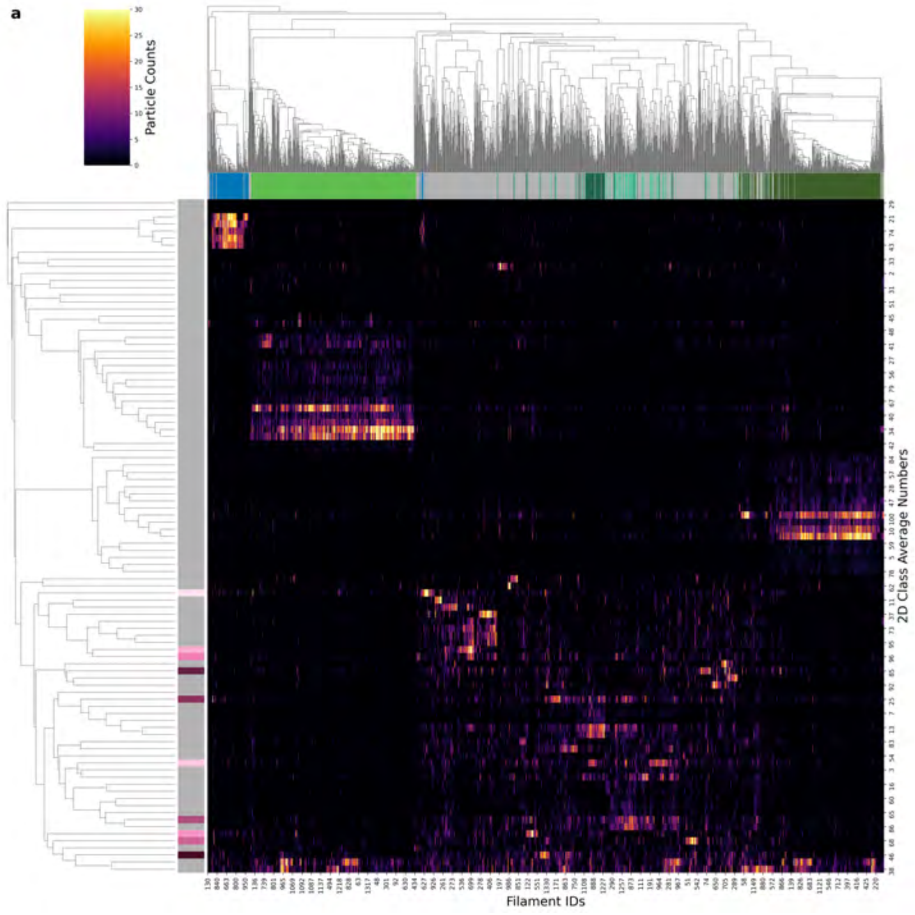

| Condition | CTE |
|---|---|
| **Timepoint** | 240 minutes |
| **Replicate** | 2 |
| **Picking** | Manual |
| **Micrographs** | 942 |
| **Extracted Particles** | 123830 |
| **Pixel Size** | 0.824 Å |
| **Classified Particles** | 123830 |

**b**

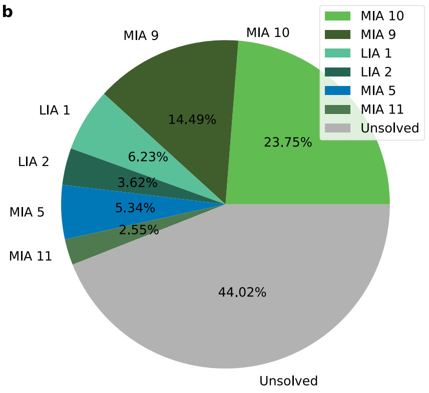

**c**

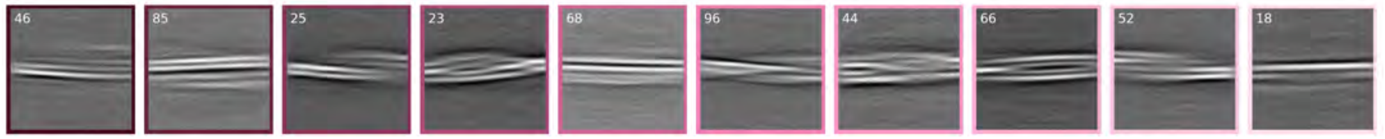

**d**

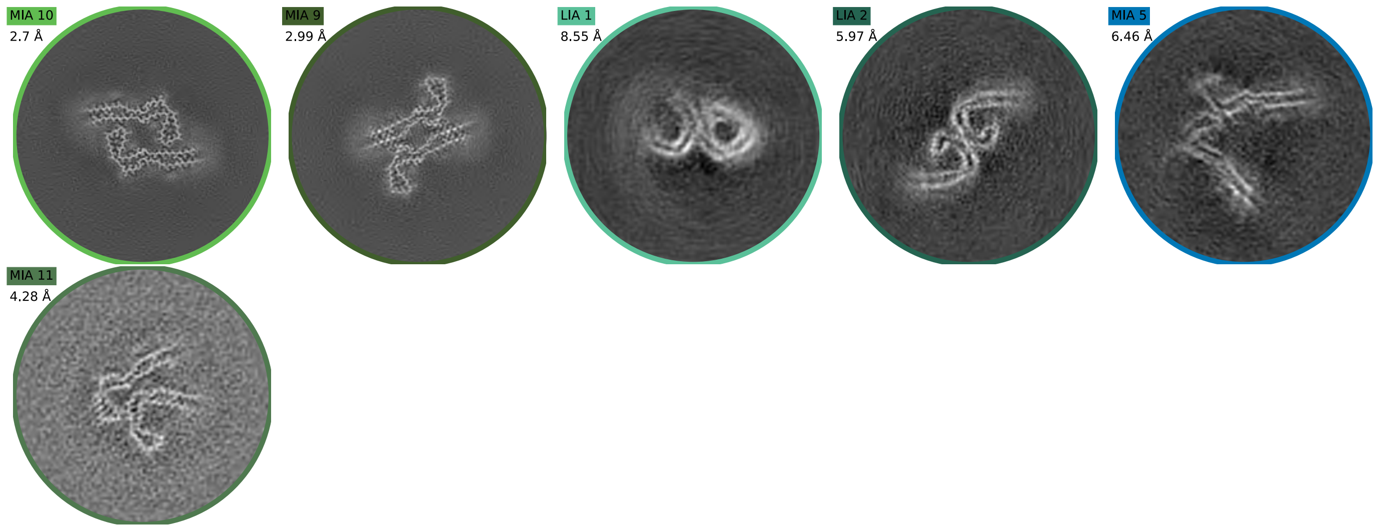

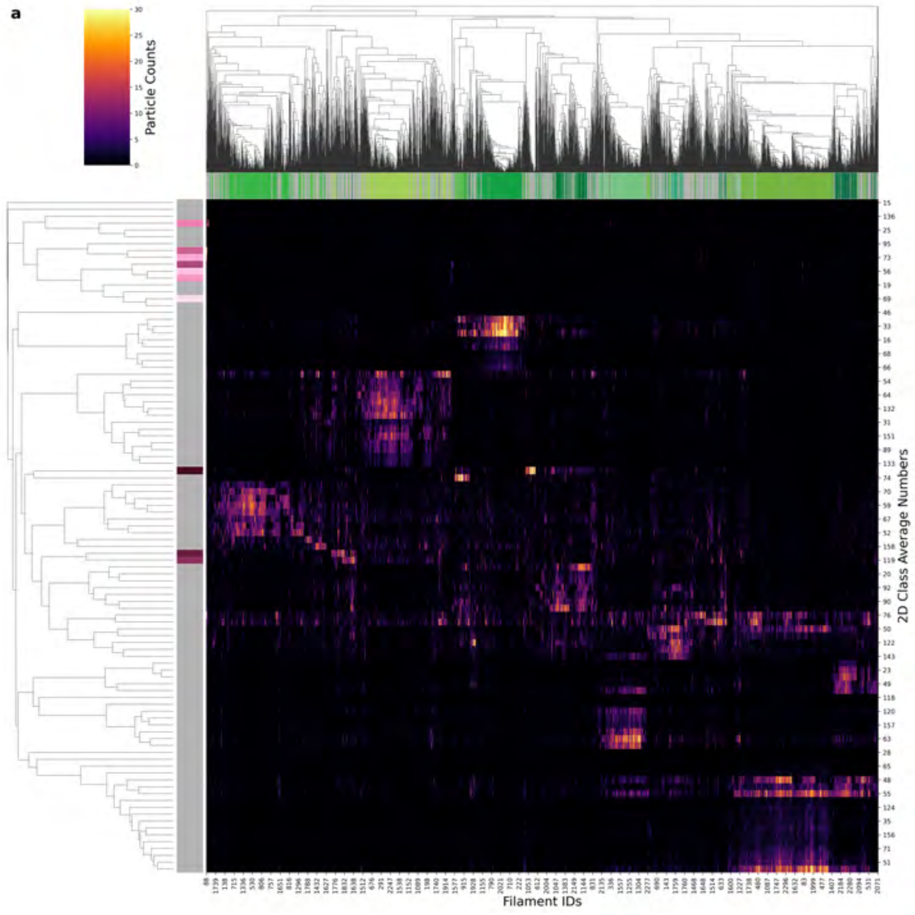

| Condition | CTE |
|---|---|
| Timepoint | 240 minutes |
| Replicate | 3 |
| Picking | Auto |
| Micrographs | 967 |
| Extracted Particles | 188025 |
| Pixel Size | 0.824 Å |
| Classified Particles | 188025 |

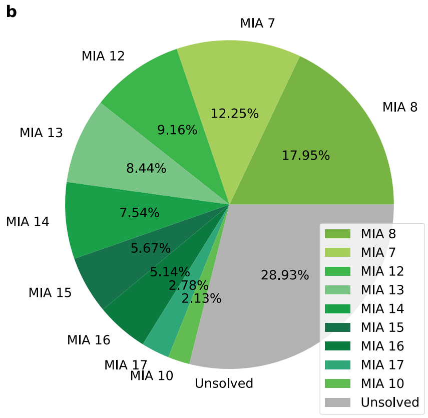

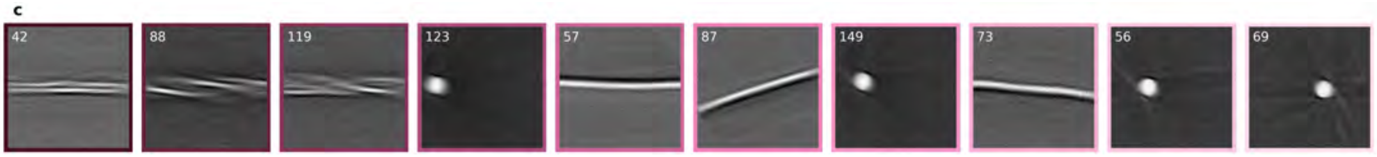

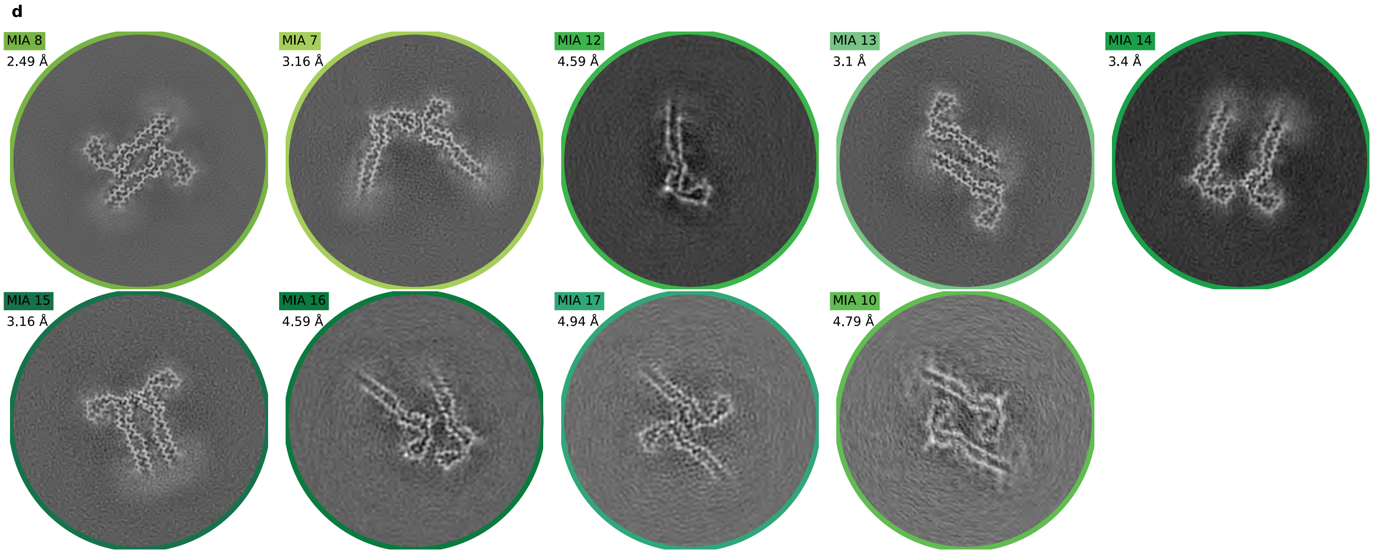

**a**

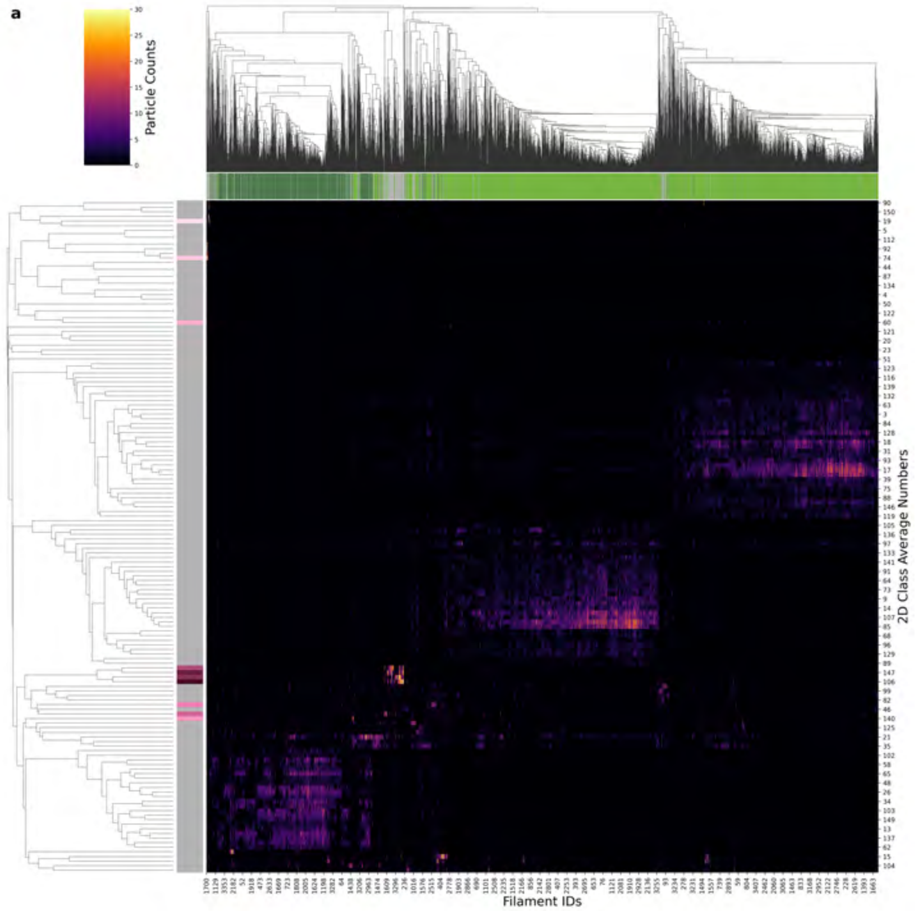

| | |
|---|---|
| **Condition** | CTE |
| **Timepoint** | 240 minutes |
| **Replicate** | 6 |
| **Picking** | Manual |
| **Micrographs** | 1341 |
| **Extracted Particles** | 234520 |
| **Pixel Size** | 0.824 Å |
| **Classified Particles** | 234520 |

**b**

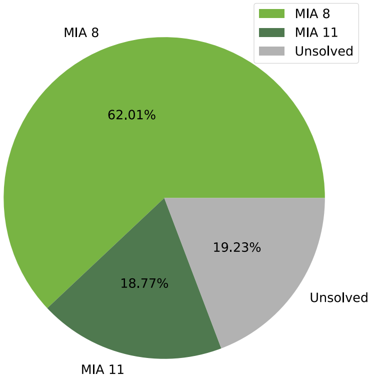

**c**

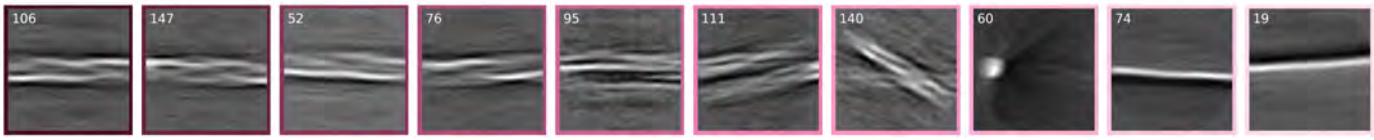

**d**

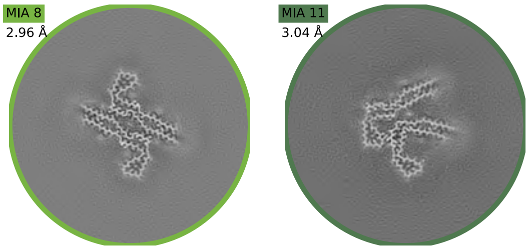

a

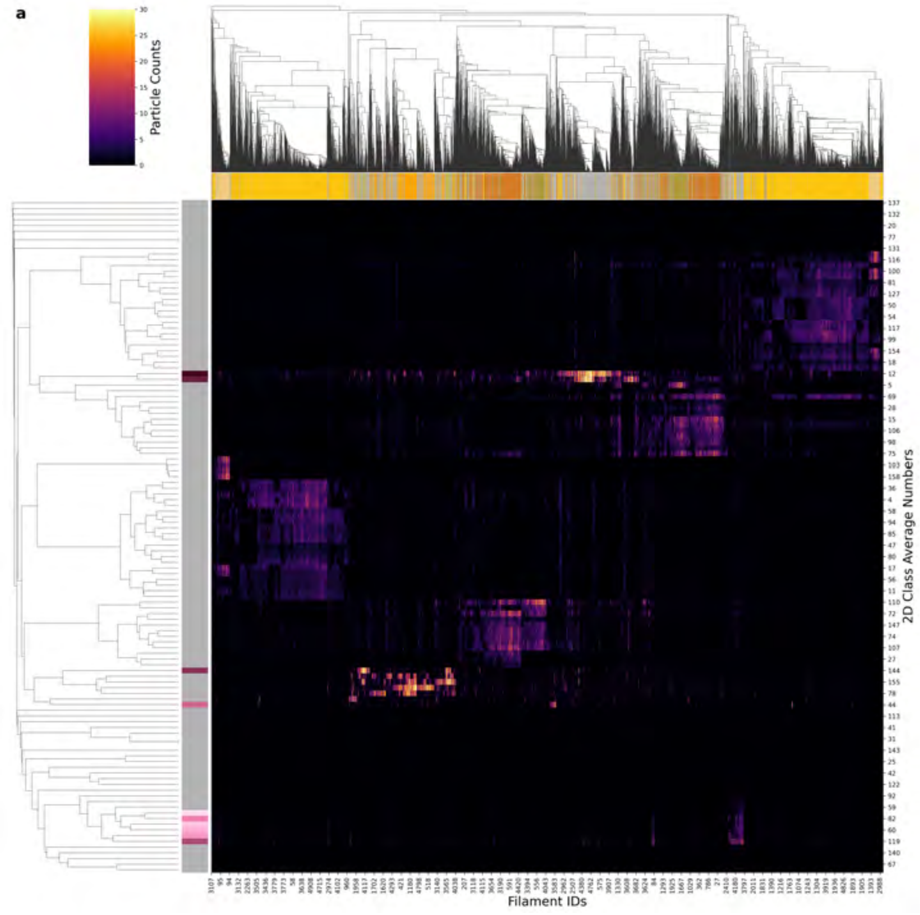

| Condition | CTE |
|---|---|
| Timepoint | 300 minutes |
| Replicate | 1 |
| Picking | Manual |
| Micrographs | 3141 |
| Extracted Particles | 317057 |
| Pixel Size | 0.727 Å |
| Classified Particles | 317057 |

b

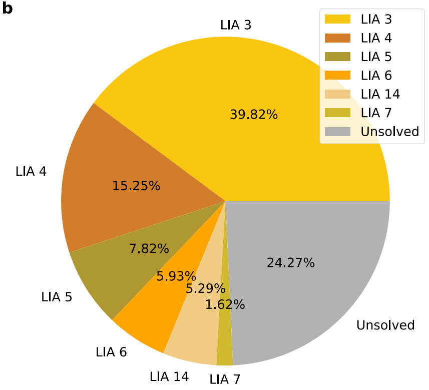

c

d

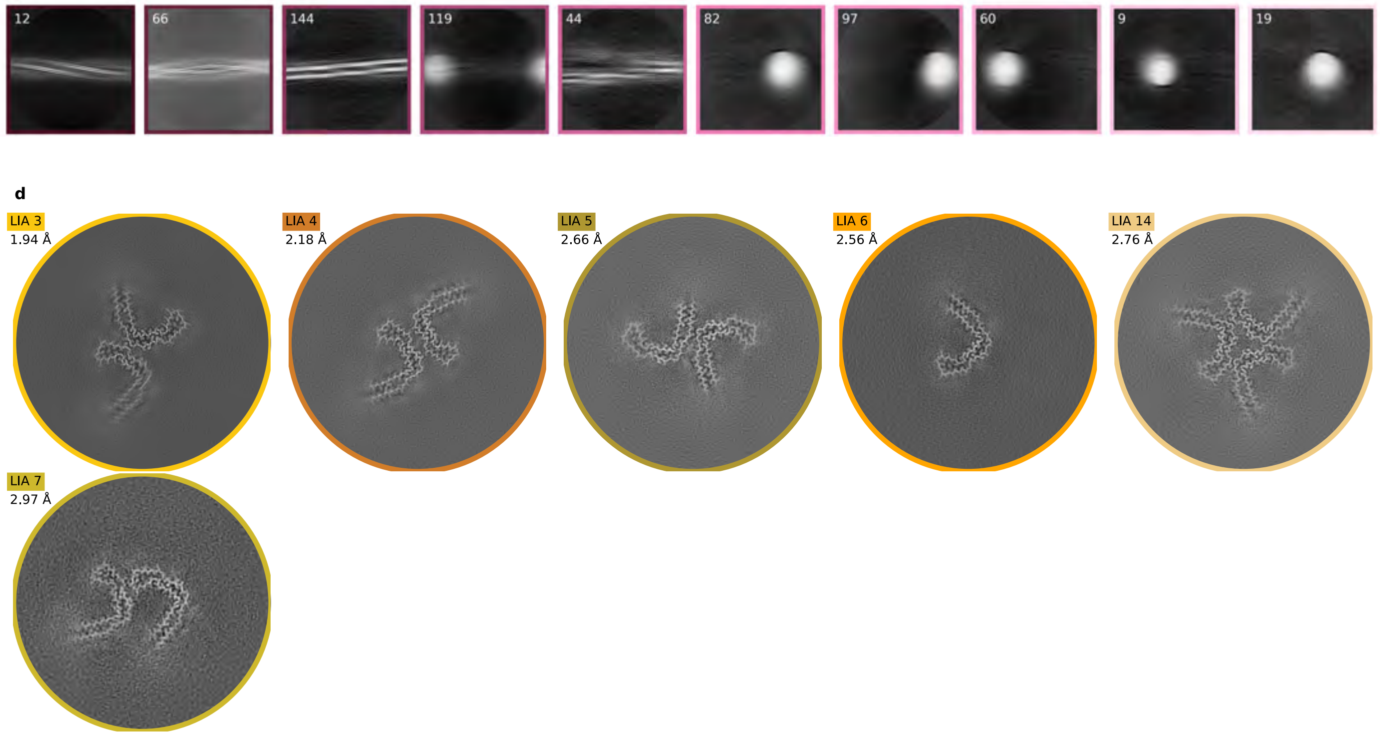

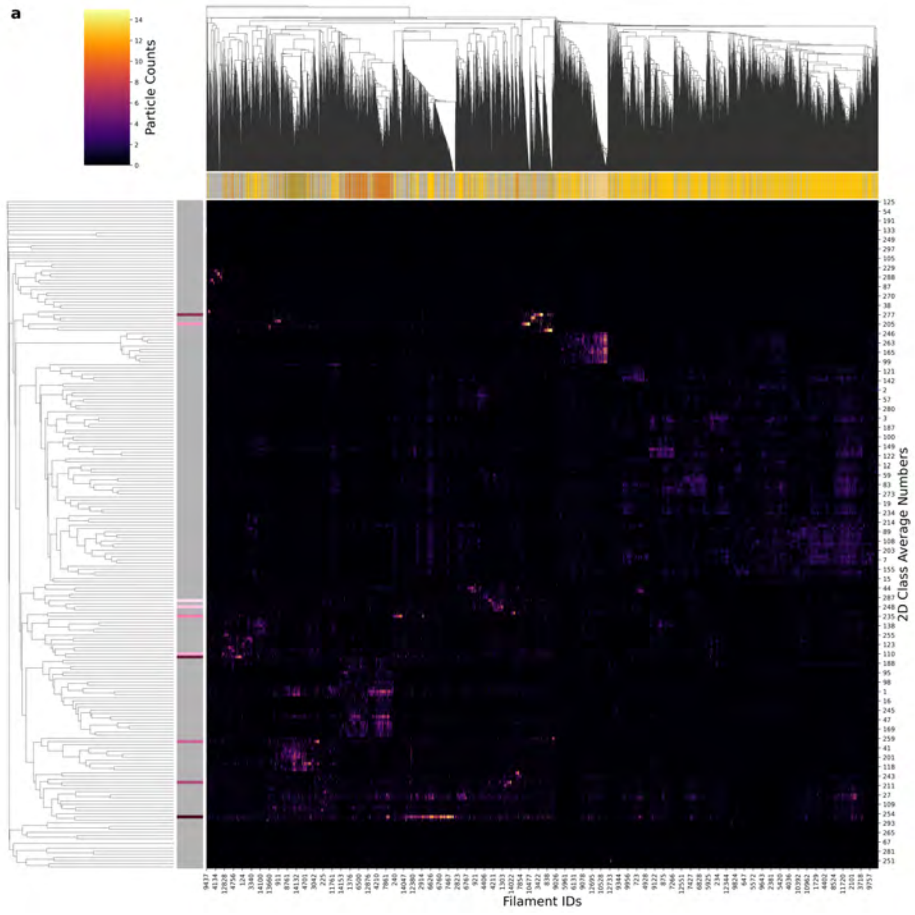

| Condition | CTE |
|---|---|
| Timepoint | 300 minutes |
| Replicate | 2 |
| Picking | Auto |
| Micrographs | 2770 |
| Extracted Particles | 2054796 |
| Pixel Size | 0.727 Å |
| Classified Particles | 540669 |

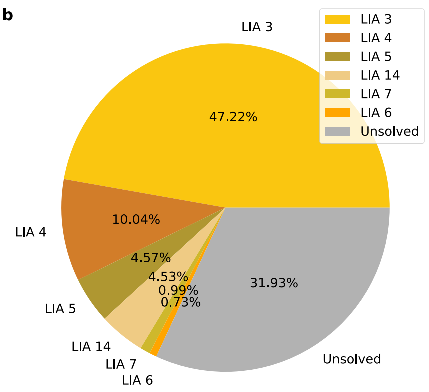

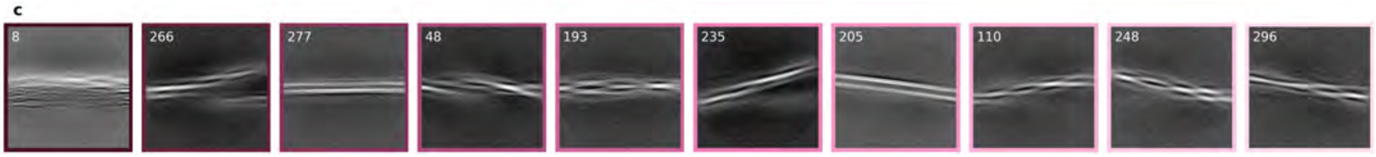

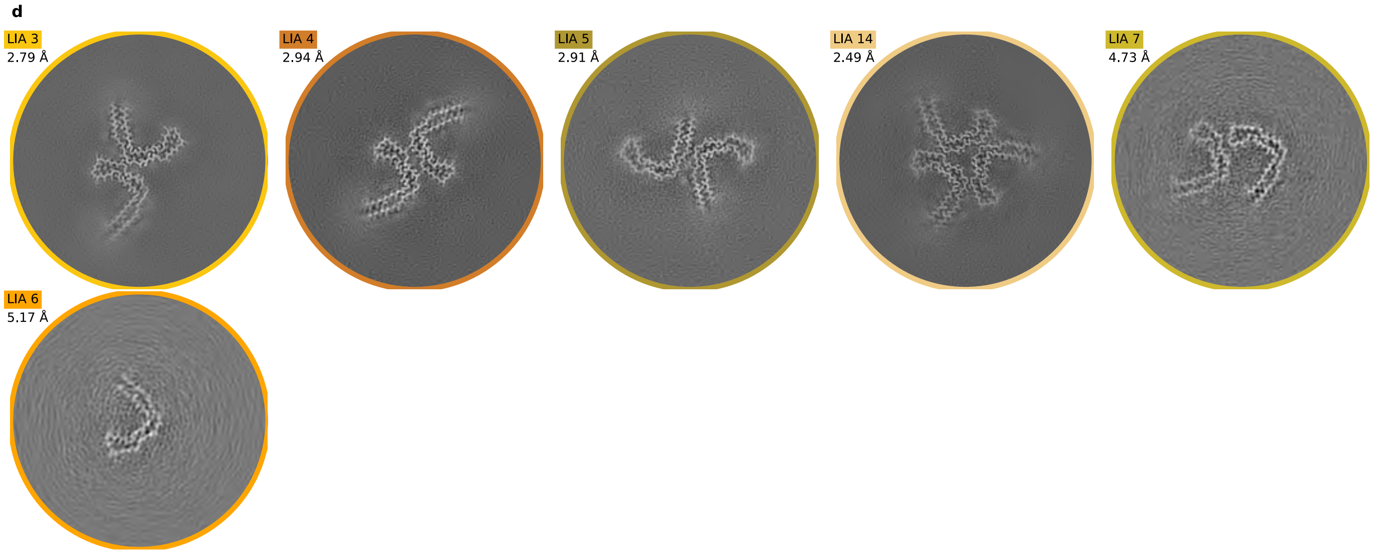

a

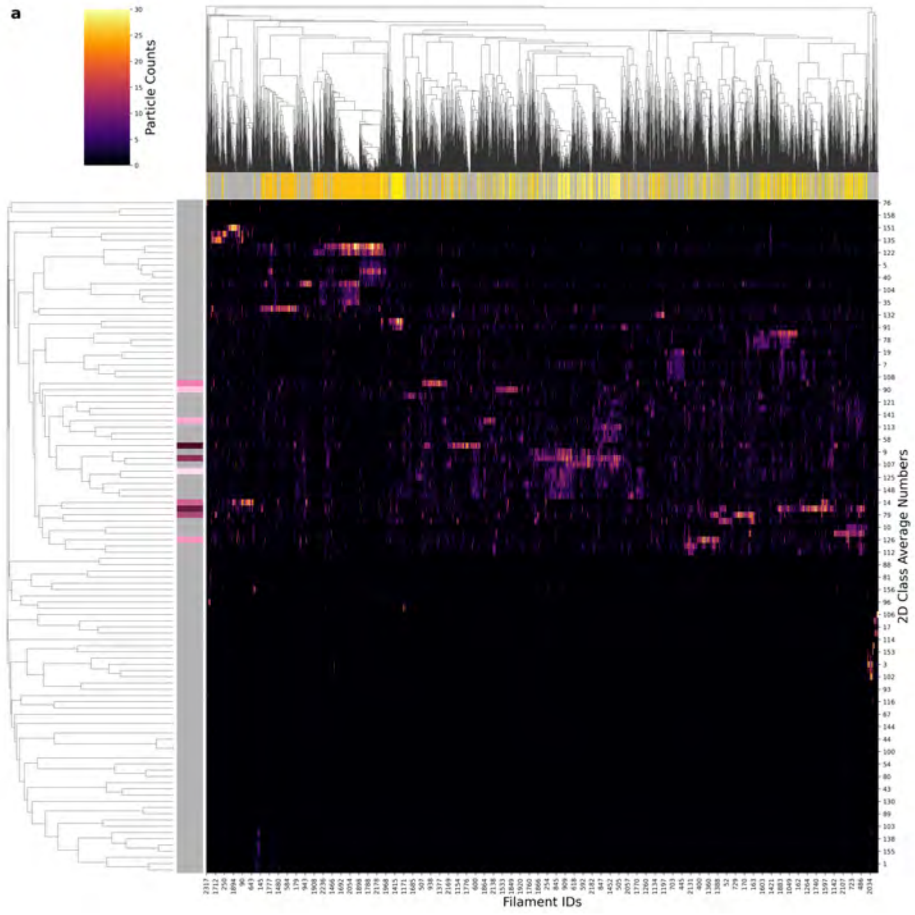

| Condition | CTE |
| --- | --- |
| **Timepoint** | 300 minutes |
| **Replicate** | 3 |
| **Picking** | Manual |
| **Micrographs** | 711 |
| **Extracted Particles** | 134013 |
| **Pixel Size** | 0.824 Å |
| **Classified Particles** | 134013 |

b

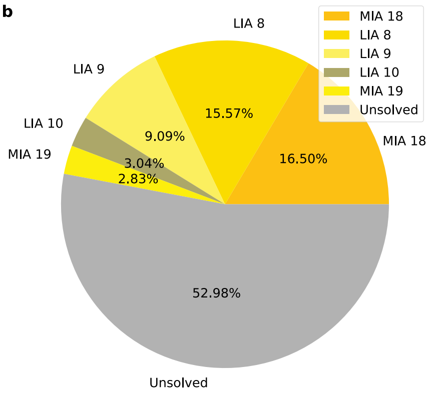

c

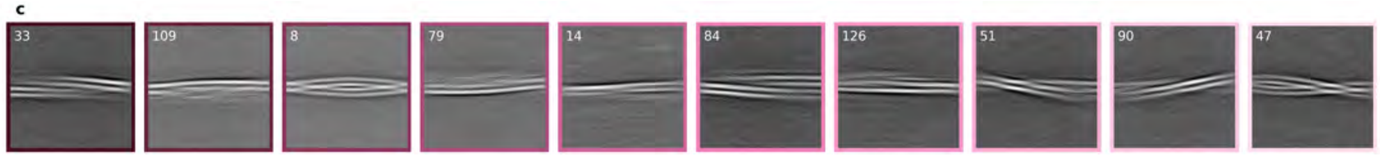

d

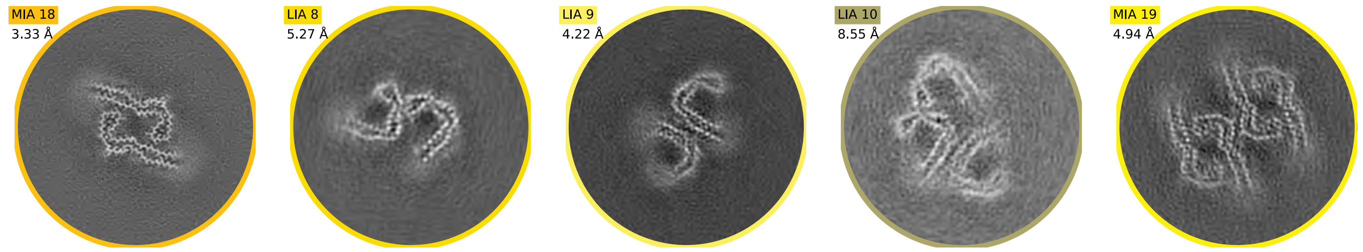

**a**

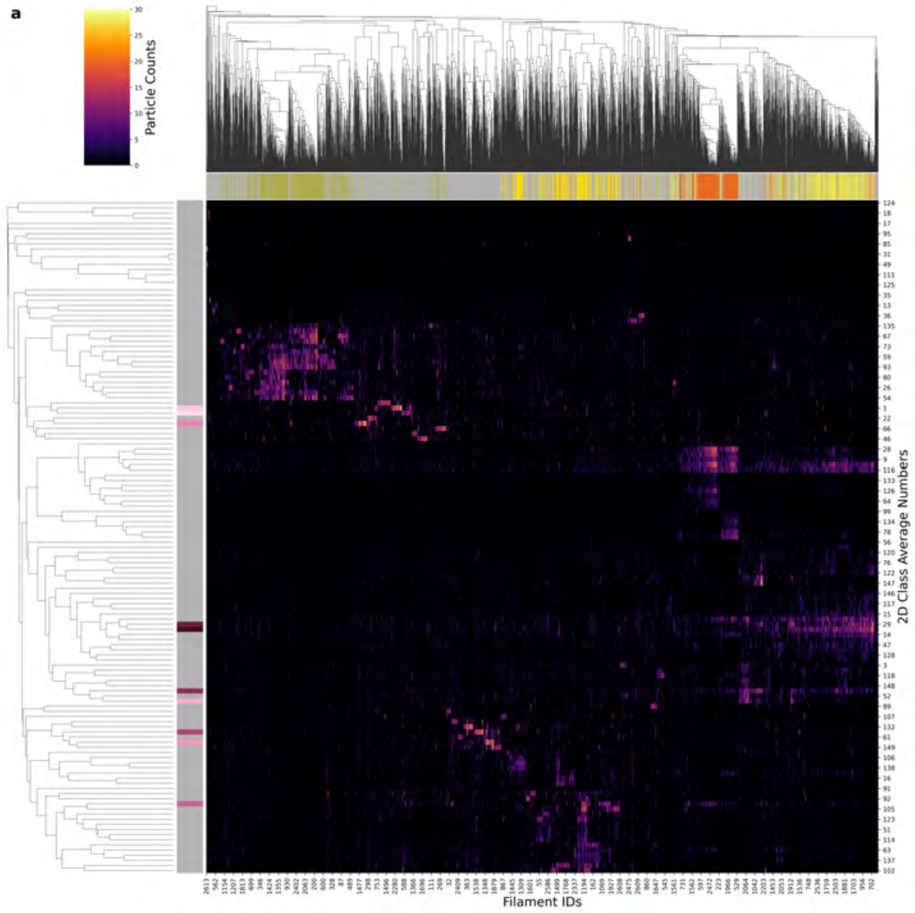

| Condition | CTE |
|---|---|
| Timepoint | 300 minutes |
| Replicate | 6 |
| Picking | Manual |
| Micrographs | 1577 |
| Extracted Particles | 181708 |
| Pixel Size | 0.824 Å |
| Classified Particles | 181708 |

**b**

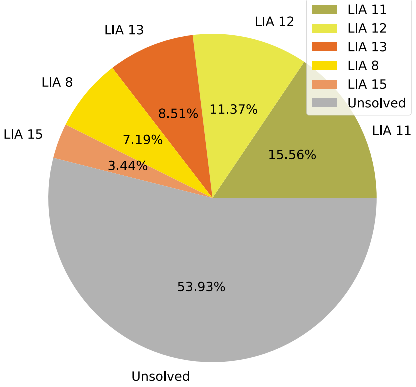

**c**

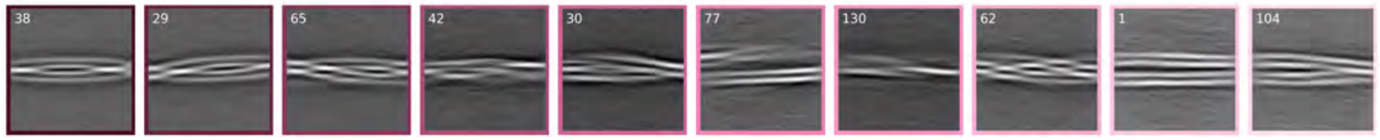

**d**

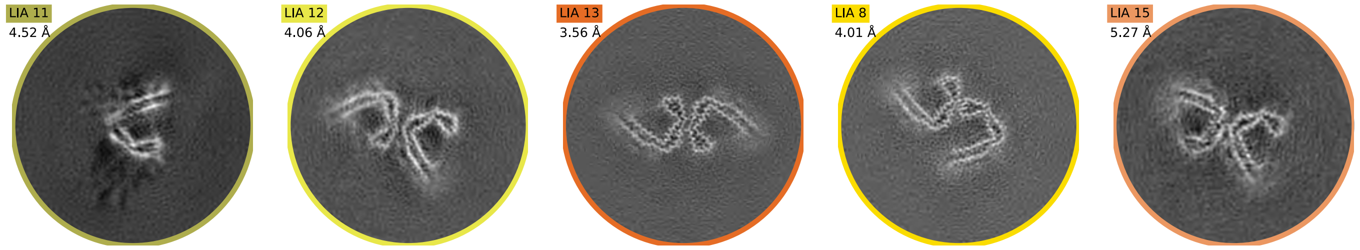

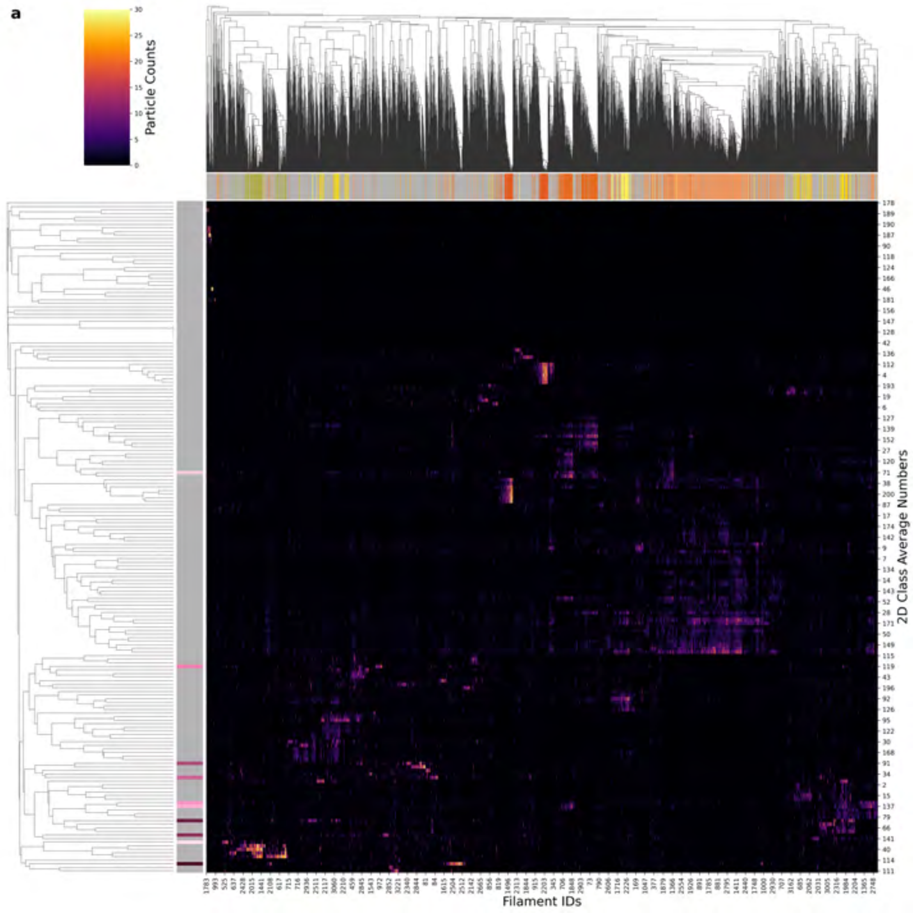

| Condition | CTE |
|---|---|
| Timepoint | 360 minutes |
| Replicate | 1 |
| Picking | Manual |
| Micrographs | 233495 |
| Extracted Particles | 2943 |
| Pixel Size | 0.824 Å |
| Classified Particles | 233495 |

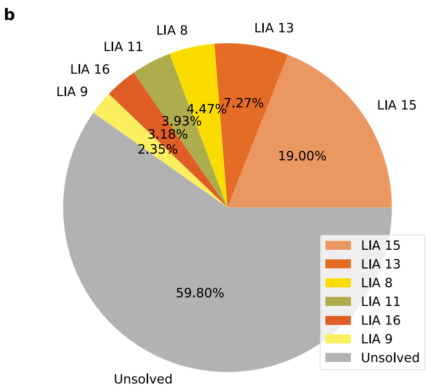

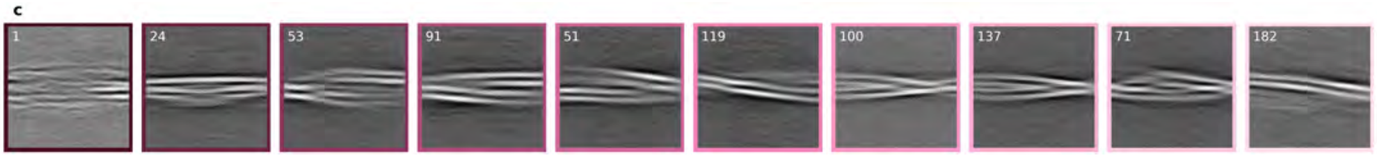

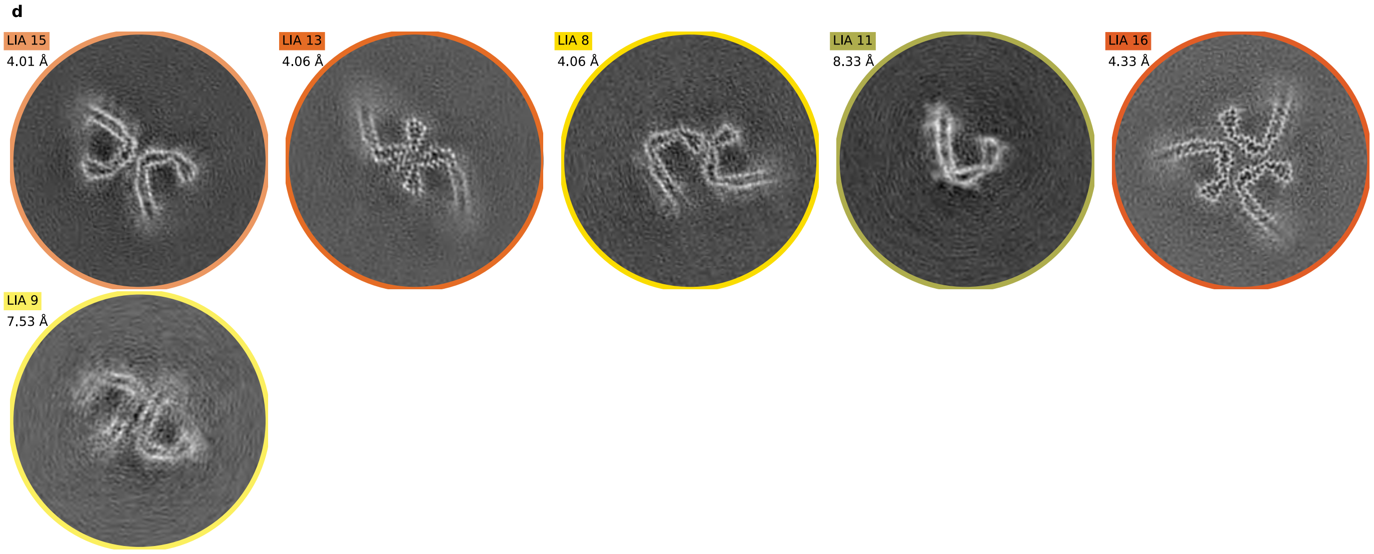

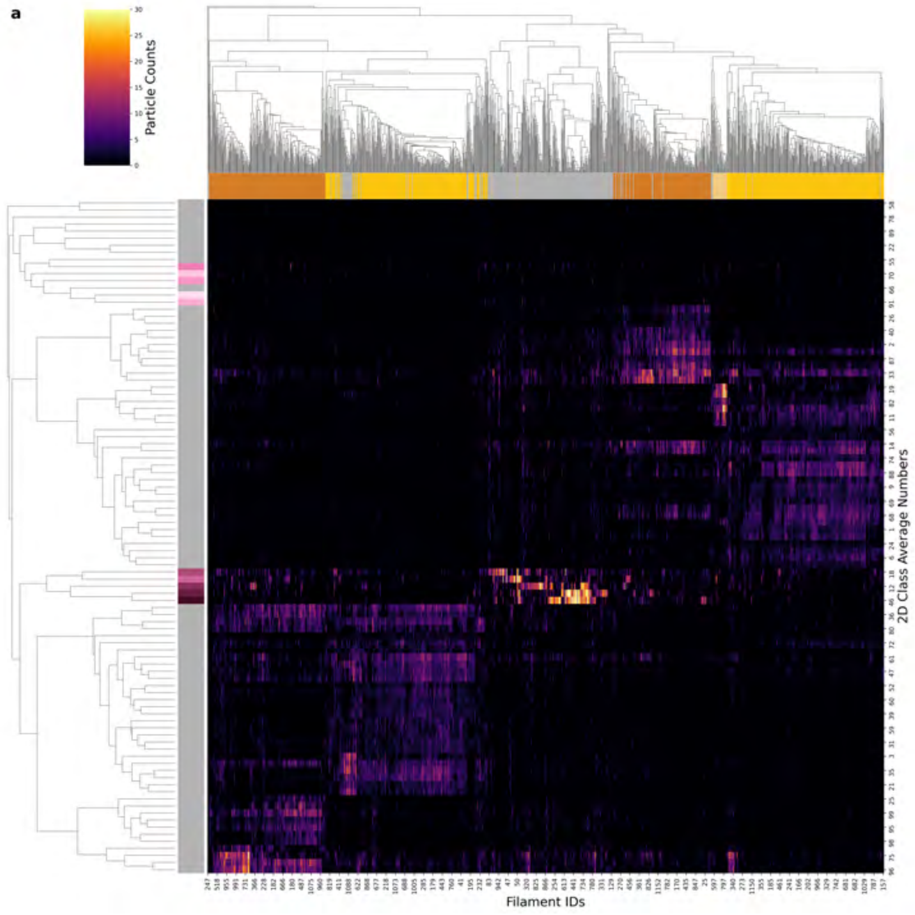

| Condition | CTE |
| --- | --- |
| Timepoint | 360 minutes |
| Replicate | 4 |
| Picking | Manual |
| Micrographs | 2183 |
| Extracted Particles | 104881 |
| Pixel Size | 0.727 Å |
| Classified Particles | 104881 |

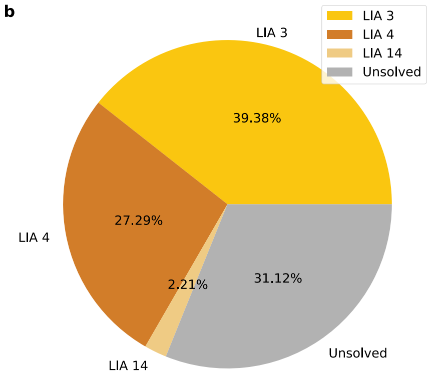

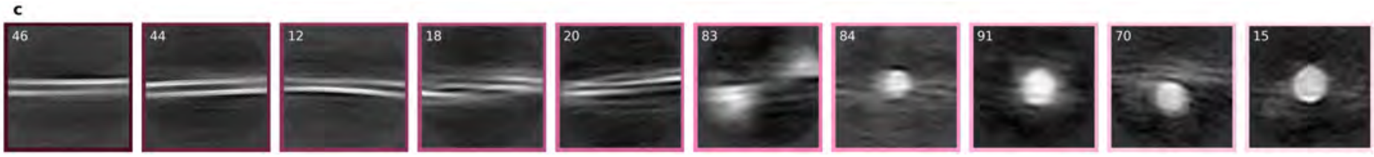

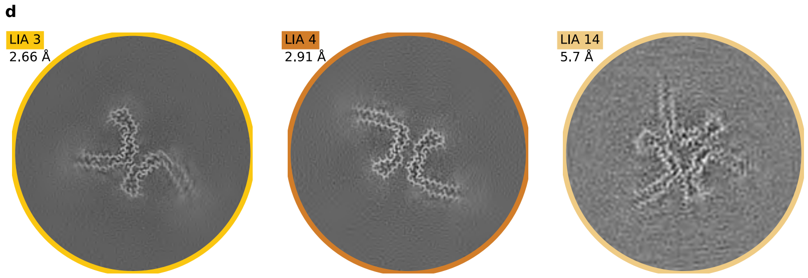

a

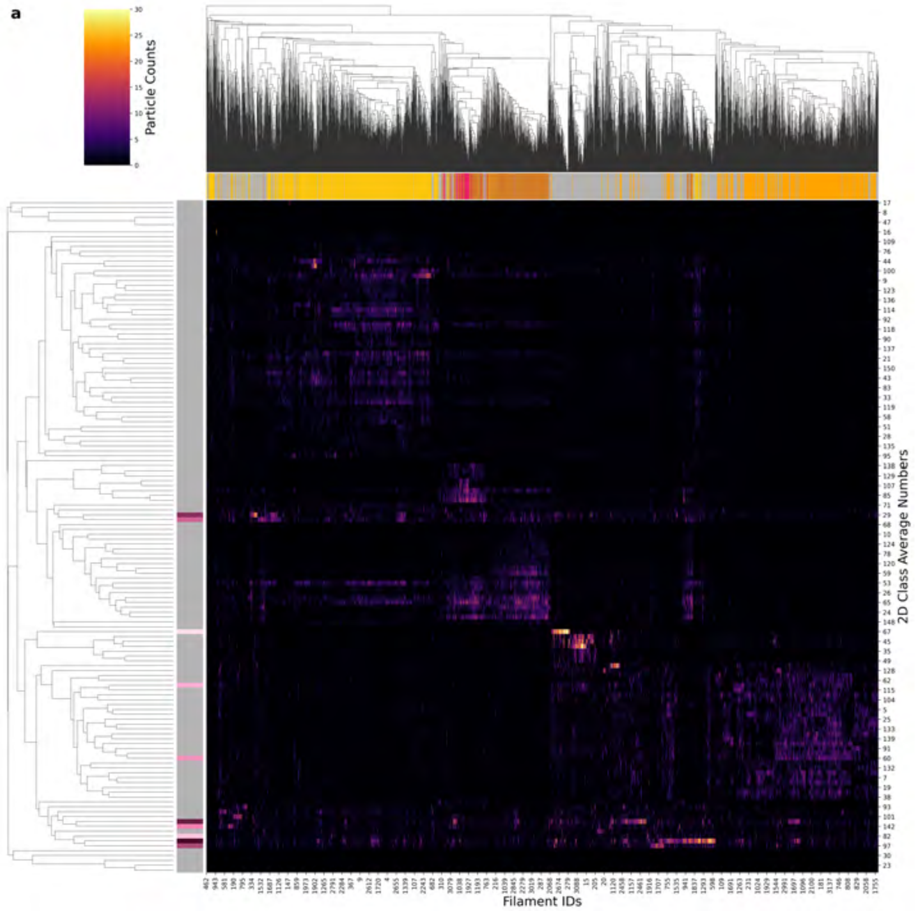

| Condition | CTE |
|---|---|
| **Timepoint** | 360 minutes |
| **Replicate** | 5 |
| **Picking** | Manual |
| **Micrographs** | 1275 |
| **Extracted Particles** | 249442 |
| **Pixel Size** | 0.824 Å |
| **Classified Particles** | 249442 |

b

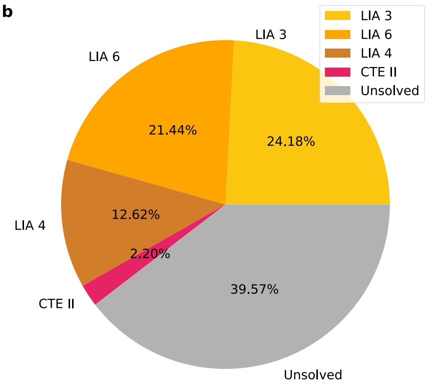

c

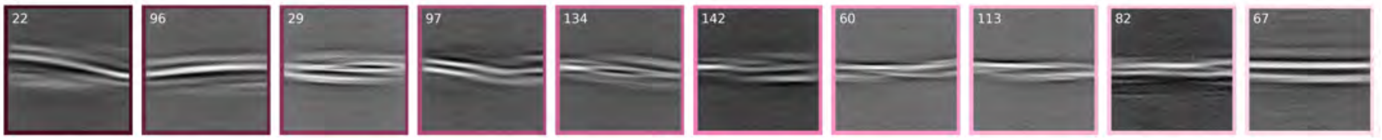

d

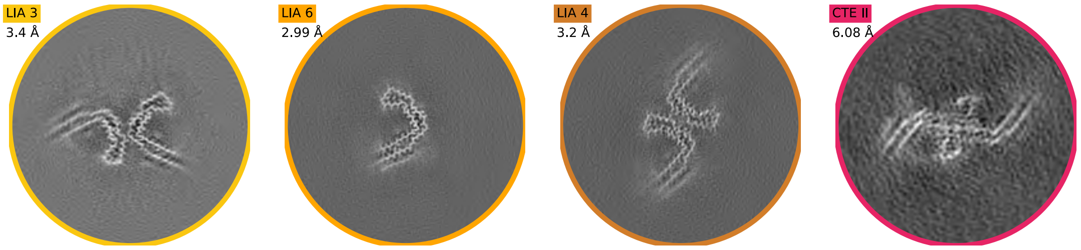

a

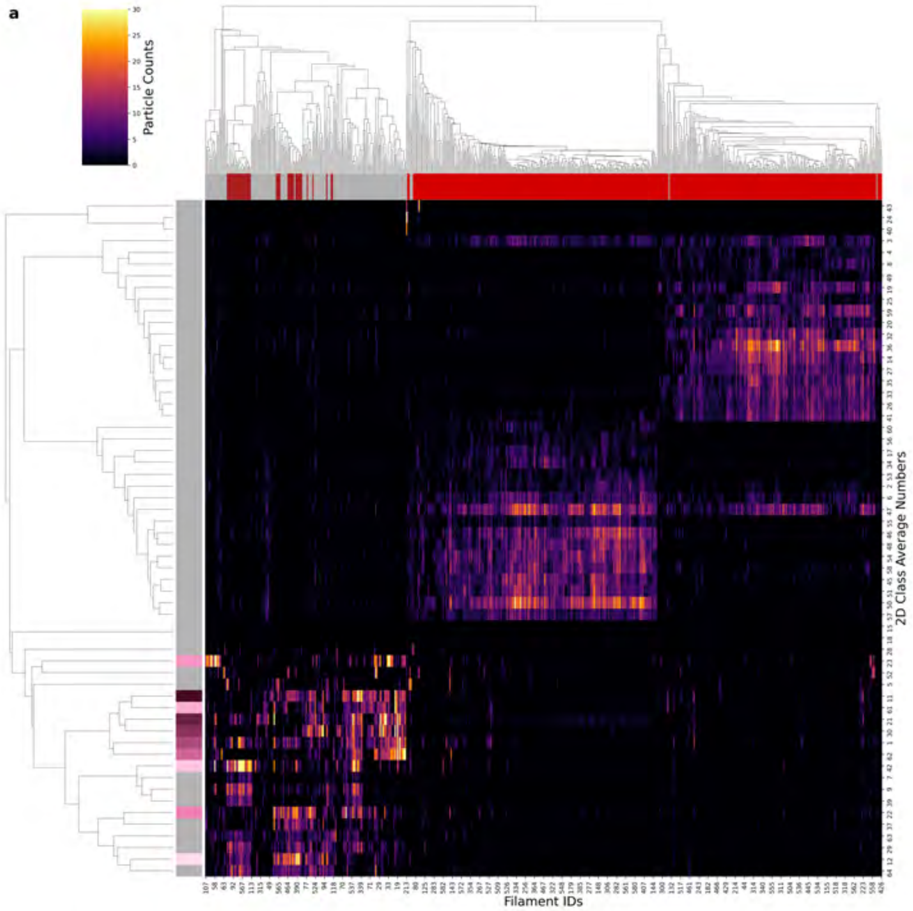

| Condition | CTE |
|---|---|
| Timepoint | 720 minutes |
| Replicate | 1 |
| Picking | Manual |
| Micrographs | 1884 |
| Extracted Particles | 53403 |
| Pixel Size | 0.824 Å |
| Classified Particles | 53403 |

b

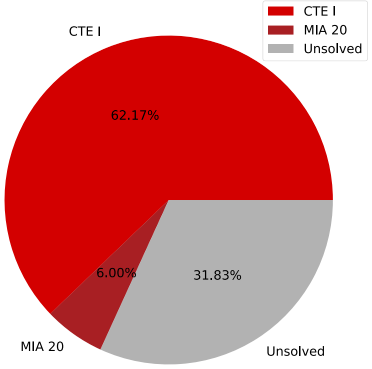

c

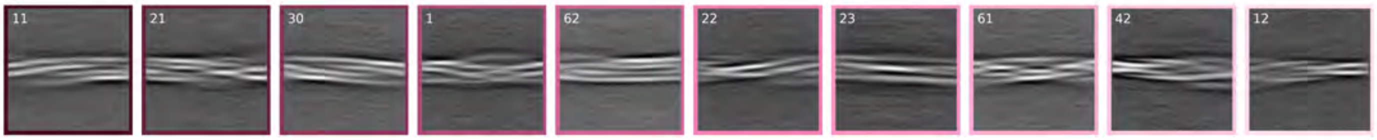

d

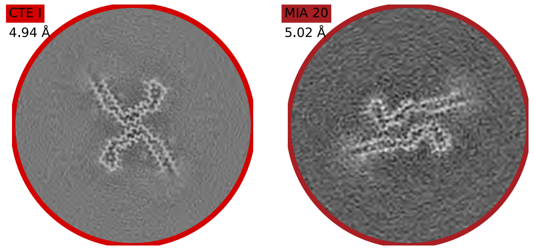

**a**

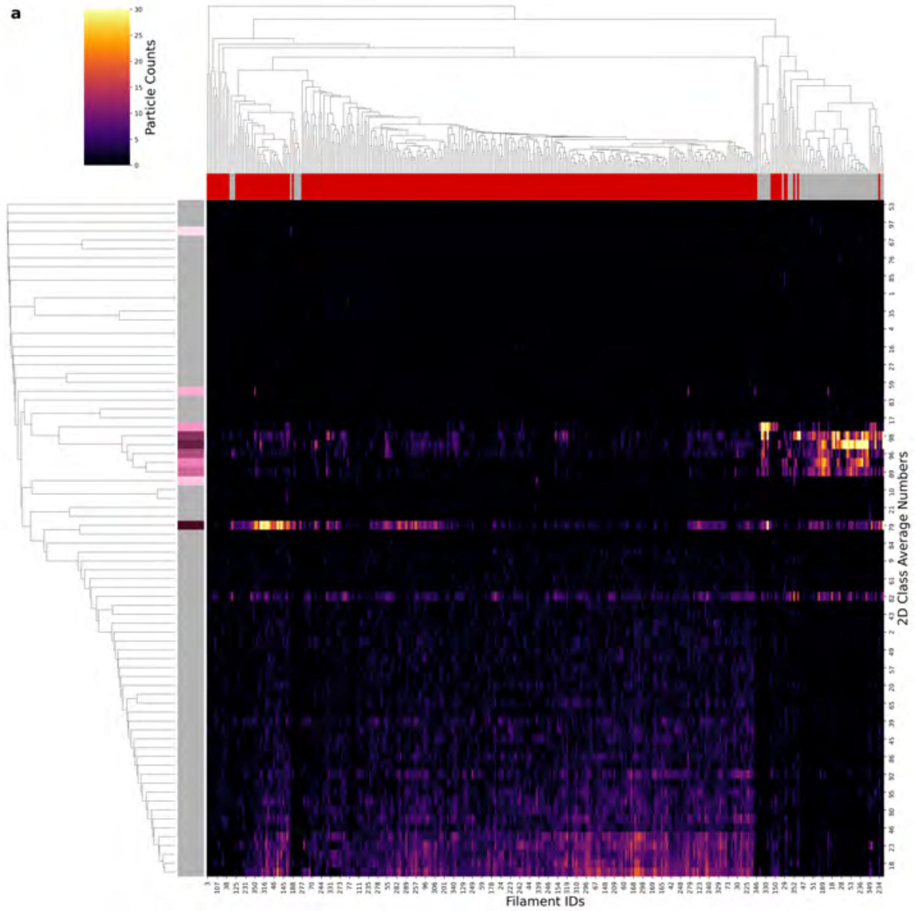

| Condition | CTE |
|---|---|
| **Timepoint** | 720 minutes |
| **Replicate** | 2 |
| **Picking** | Manual |
| **Micrographs** | 1980 |
| **Extracted Particles** | 32022 |
| **Pixel Size** | 0.824 Å |
| **Classified Particles** | 32022 |

**b**

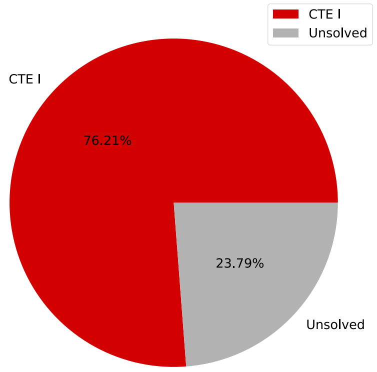

**c**

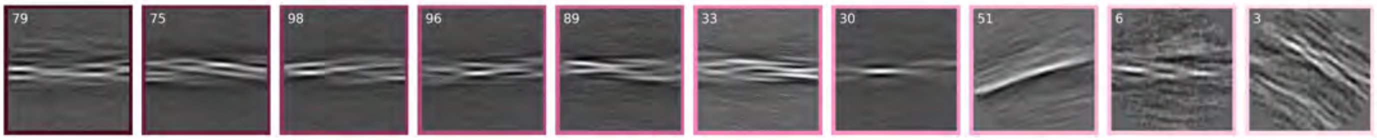

**d**

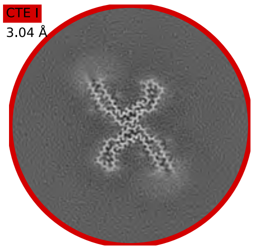

a

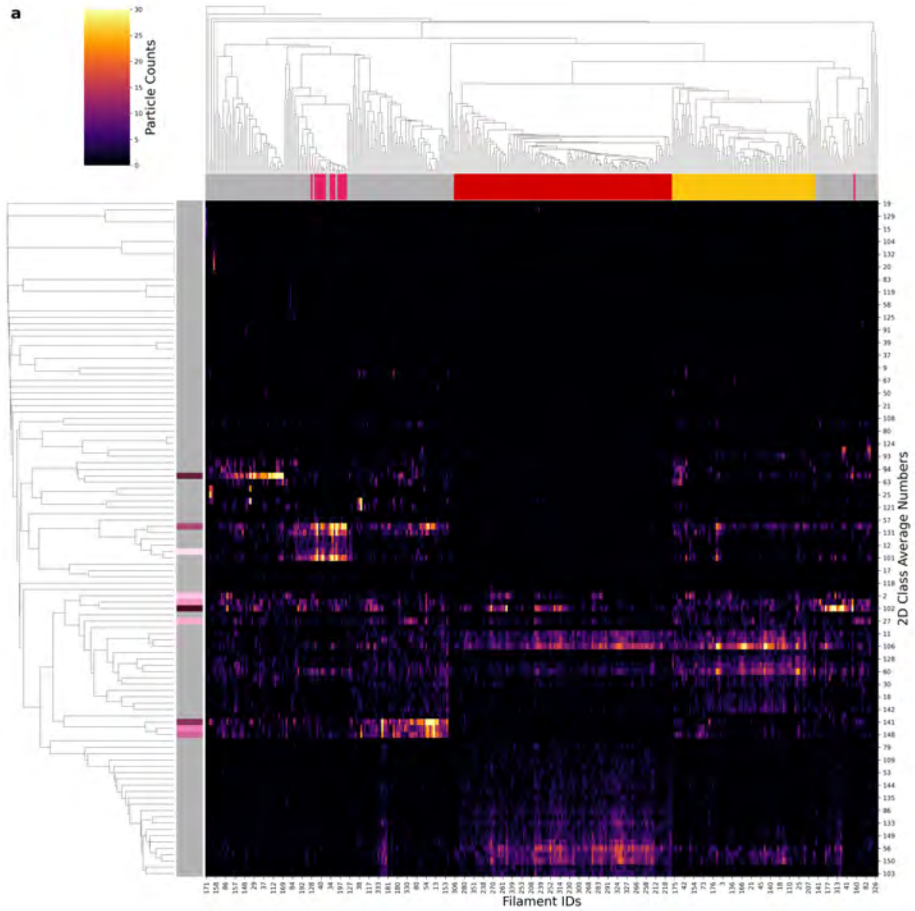

| Condition | CTE |
|---|---|
| **Timepoint** | 720 minutes |
| **Replicate** | 3 |
| **Picking** | Manual |
| **Micrographs** | 1599 |
| **Extracted Particles** | 32580 |
| **Pixel Size** | 0.824 Å |
| **Classified Particles** | 32580 |

b

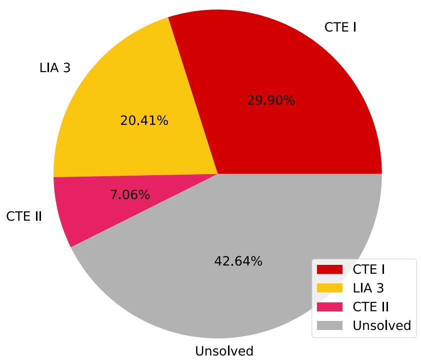

c

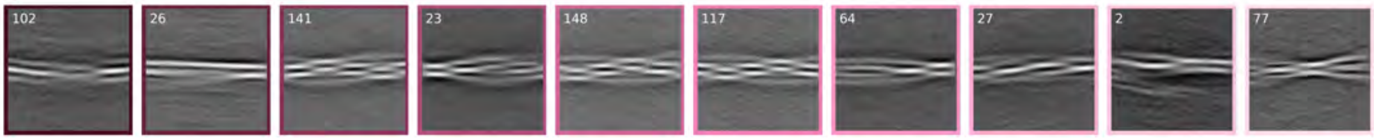

d

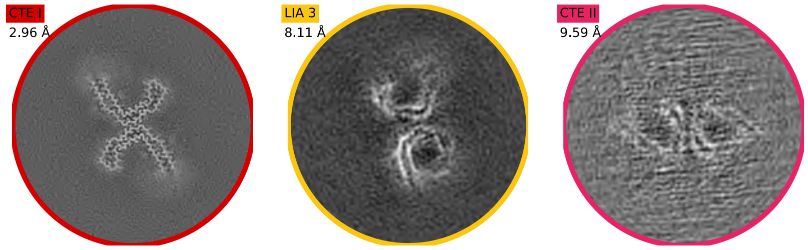

**a**

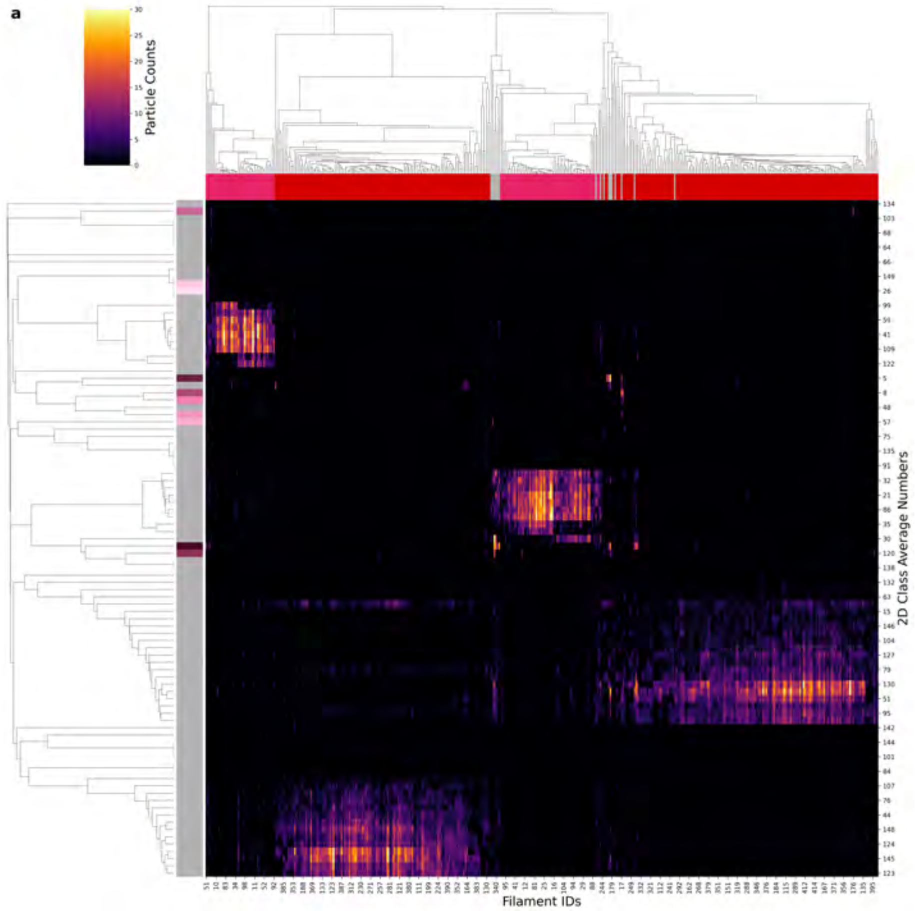

| Condition | CTE |
|---|---|
| **Timepoint** | 720 minutes |
| **Replicate** | 4 |
| **Picking** | Manual |
| **Micrographs** | 2580 |
| **Extracted Particles** | 36037 |
| **Pixel Size** | 0.824 Å |
| **Classified Particles** | 36037 |

**b**

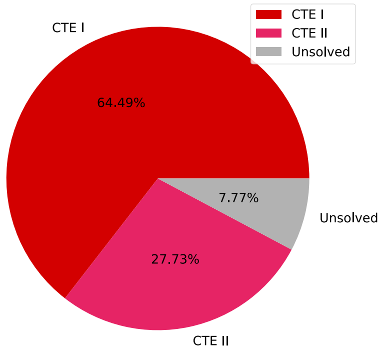

**c**

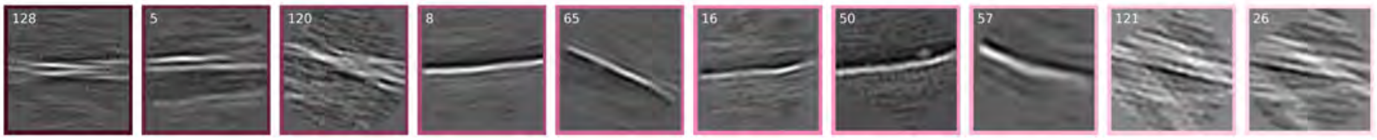

**d**

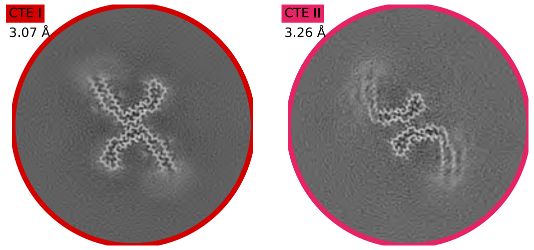

**a**

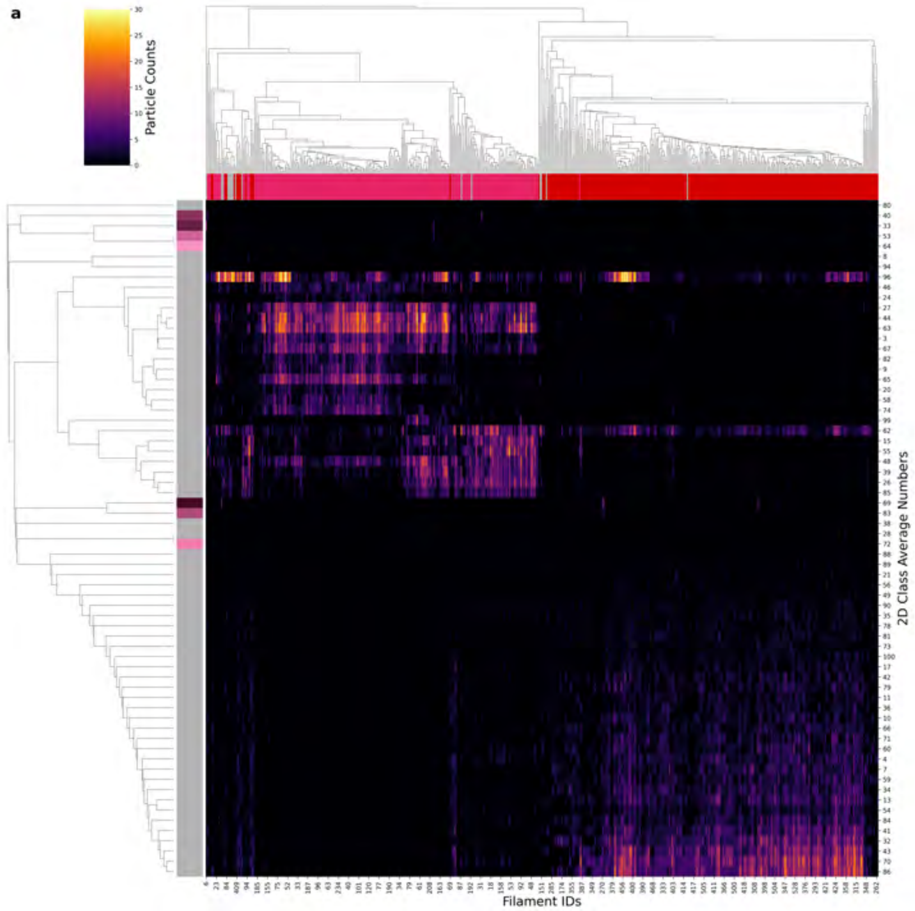

| | |
|---|---|
| **Condition** | CTE |
| **Timepoint** | 720 minutes |
| **Replicate** | 5 |
| **Picking** | Manual |
| **Micrographs** | 2580 |
| **Extracted Particles** | 46371 |
| **Pixel Size** | 0.824 Å |
| **Classified Particles** | 43926 |

**b**

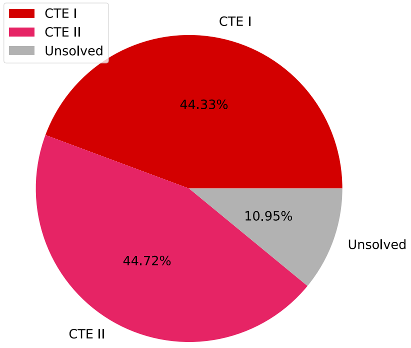

**c**

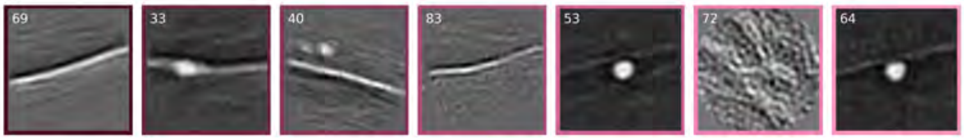

**d**

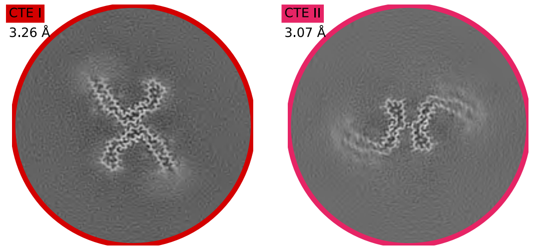

### FSC AD reactions

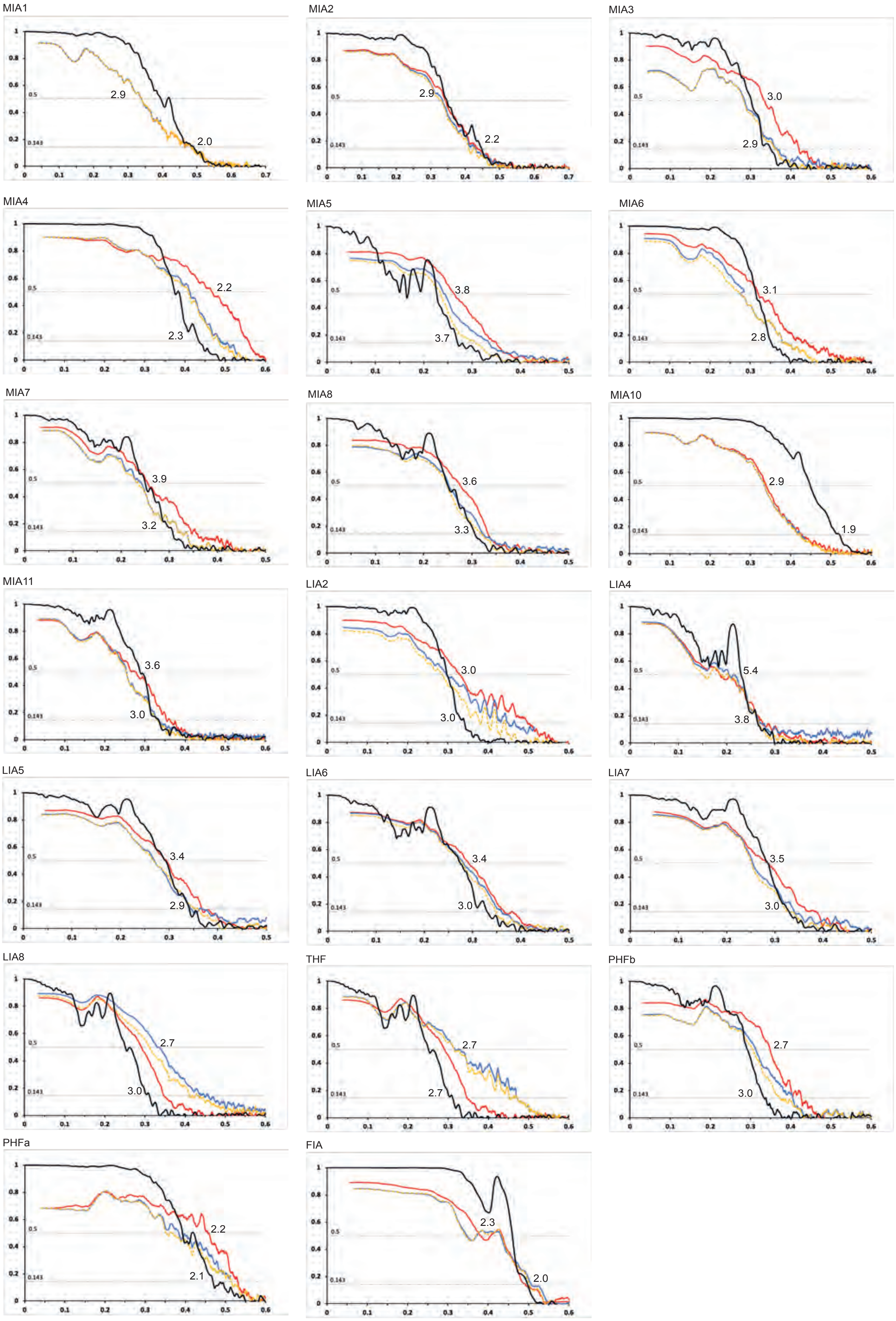

### FSC CTE reactions

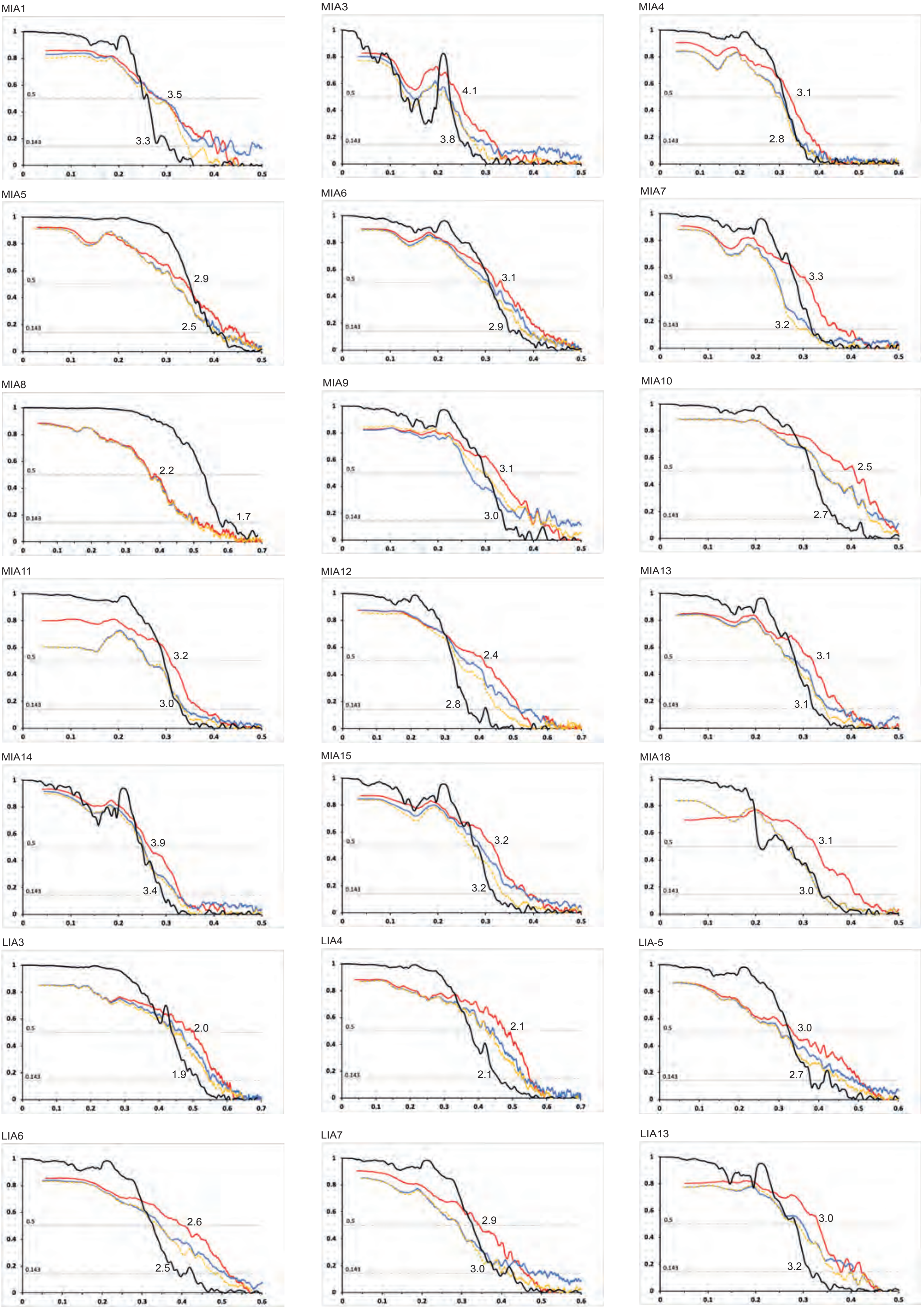

### FSC CTE reactions

LIA14

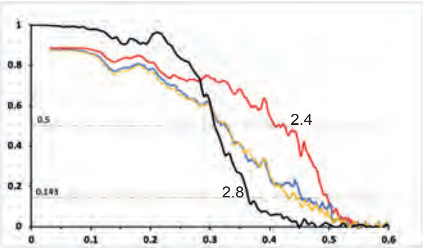

CTE I

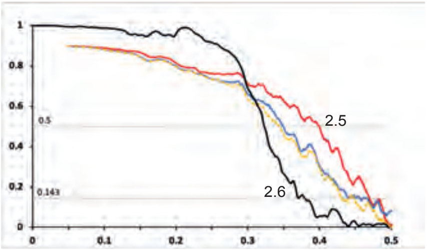

CTE I

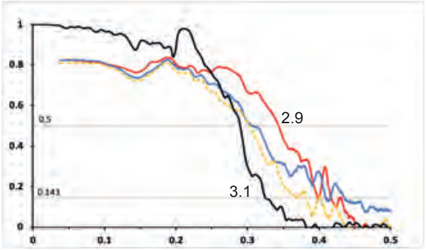

### Cryo-EM data acquisition

| | TFS Krios G4 | LMB Krios G2 |
|---|---|---|
| **Data acquisition** | | |
| Electron gun | CFEG | FEG |
| Detector | Falcon 4i | Falcon 4 |
| Energy filter slit (eV) | 10 | - |
| Magnification | 165,000 × | 105,000 × |
| Voltage (kV) | 300 | 300 |
| Electron dose (e-/Å$^2$) | 40 | 30 |
| Defocus range (µm) | 0.5 to 2.5 | 0.5 to 2.5 |
| Pixel size (Å) | 0.727 | 0.824 |

| **TFS Krios G4** | **AD-FIA**<br>(120' R1)<br>(EMDB 17806)<br>(PDB 8PPO) |
|---|---|
| **Data acquisition** | |
| Electron gun | CFEG |
| Detector | Falcon 4i |
| Energy filter slit (eV) | 10 |
| Magnification | 165,000 |
| Voltage (kV) | 300 |
| Electron dose (e-/Å$^2$) | 40 |
| Defocus range (μM) | 0.5 to 2.5 |
| Pixel size (Å) | 0.727 |
| **Data processing** | |
| Initial particle images (no.) | 1680613 |
| Final particle images (no.) | 58703 |
| Helical twist (°) | 176.886 |
| Helical rise (Å) | 2.3667 |
| Symmetry imposed | $2_1$ |
| Map resolution FSC 0.143 (Å) | 2.0 |
| **Refinement** | |
| Initial model used (PDB code) | de novo |
| Model resolution FSC 0.5 (Å) | 2.5 |
| Map sharpening *B* factor (Å$^2$) | -36.32 |
| Model composition | |
| Non-hydrogen atoms | 1696 |
| Protein residues | 240 |
| Ligands | 0 |
| *B* factors (Å$^2$) | |
| Protein | 35.13 |
| Ligand | na |
| R.m.s. deviations | |
| Bond lengths (Å) | 0.007 |
| Bond angles (°) | 1.197 |
| Validation | |
| MolProbity score | 1.05 |
| Clashscore | 0.29 |
| Poor rotamers (%) | 0 |
| Ramachandran plot | |
| Favored (%) | 93.27 |
| Allowed (%) | 6.73 |
| Disallowed (%) | 0 |

| **TFS Krios G4** | **AD-MIA1 (180’ R2)** (EMDB-xxxx) (PDB XXXX) | **AD-MIA2 (180’ R2)** (EMDB-xxxx) (PDB XXXX) |
|---|---|---|
| **Data acquisition** | | |
| Electron gun | CFEG | CFEG |
| Detector | Falcon 4i | Falcon 4i |
| Energy filter slit (eV) | 10 | 10 |
| Magnification | 165,000 | 165,000 |
| Voltage (kV) | 300 | 300 |
| Electron dose (e-/Å$^2$) | 40 | 40 |
| Defocus range (μM) | 0.5 to 2.5 | 0.5-2.5 |
| Pixel size (Å) | 0.727 | 0.727 |
| **Data processing** | | |
| Initial particle images (no.) | 138115 | 138115 |
| Final particle images (no.) | 77889 | 21130 |
| Helical twist (°) | -1.11 | -1.22 |
| Helical rise (Å) | 4.71 | 4.75 |
| Symmetry imposed | 1 | 1 |
| Map resolution FSC 0.143 (Å) | 2.02 | 2.23 |
| **Refinement** | | |
| Initial model used (PDB code) | ModelAngelo | ModelAngelo |
| Model resolution FSC 0.5 (Å) | 2.9 | 2.9 |
| Map sharpening *B* factor (Å$^2$) | -24.9 | -25.68 |
| Model composition | | |
| Non-hydrogen atoms | 3435 | 3507 |
| Protein residues | 453 | 462 |
| Ligands | 0 | 0 |
| *B* factors (Å$^2$) | | |
| Protein | 93.54 | 96.76 |
| Ligand | na | Na |
| R.m.s. deviations | | |
| Bond lengths (Å) | 0.01 | 0.01 |
| Bond angles (°) | 1.871 | 1.838 |
| Validation | | |
| MolProbity score | 0.83 | 0.56 |
| Clashscore | 0.57 | 0.14 |
| Poor rotamers (%) | 1.27 | 0 |
| Ramachandran plot | | |
| Favored (%) | 97.73 | 98.22 |
| Allowed (%) | 2.27 | 1.78 |
| Disallowed (%) | 0 | 0 |

| TFS Krios G4 | AD-MIA3 (180' R3) (EMDB-xxxx) (PDB XXXX) | AD-MIA4 (180' R3) (EMDB-xxxx) (PDB XXXX) |
|---|---|---|
| **Data acquisition** | | |
| Electron gun | CFEG | CFEG |
| Detector | Falcon 4i | Falcon 4i |
| Energy filter slit (eV) | 10 | 10 |
| Magnification | 165,000 | 165,000 |
| Voltage (kV) | 300 | 300 |
| Electron dose (e-/Å$^2$) | 40 | 40 |
| Defocus range (μM) | 0.5 to 2.5 | 0.5 to 2.5 |
| Pixel size (Å) | 0.727 | 0.727 |
| **Data processing** | | |
| Initial particle images (no.) | 159594 | 159594 |
| Final particle images (no.) | 13033 | 94975 |
| Helical twist (°) | -1.29 | -1.44 |
| Helical rise (Å) | 4.75 | 4.74 |
| Symmetry imposed | 1 | C2 |
| Map resolution FSC 0.143 (Å) | 2.88 | 2.32 |
| **Refinement** | | |
| Initial model used (PDB code) | de novo | ModelAngelo |
| Model resolution FSC 0.5 (Å) | 3.0 | 2.2 |
| Map sharpening *B* factor (Å$^2$) | -57.56 | -62.15 |
| Model composition | | |
| Non-hydrogen atoms | 3444 | 3354 |
| Protein residues | 450 | 438 |
| Ligands | 0 | 0 |
| *B* factors (Å$^2$) | | |
| Protein | 96.09 | 40.74 |
| Ligand | na | na |
| R.m.s. deviations | | |
| Bond lengths (Å) | 0.011 | 0.012 |
| Bond angles (°) | 1.884 | 2.330 |
| Validation | | |
| MolProbity score | 0.94 | 0.86 |
| Clashscore | 0.43 | 1.32 |
| Poor rotamers (%) | 0.00 | 0 |
| Ramachandran plot | | |
| Favored (%) | 95.89 | 98.12 |
| Allowed (%) | 4.11 | 1.88 |
| Disallowed (%) | 0 | 0 |

| TFS Krios G4 | AD-MIA5 (240’ R3) (EMDB-xxxx) (PDB-xxxx) | AD-MIA8 (240’ R3) (EMDB-xxxx) (PDB-xxxx) | AD-LIA8 (240’ R3) (EMDB-xxxx) (PDB-xxxx) |
|---|---|---|---|
| **Data acquisition** | | | |
| Electron gun | CFEG | CFEG | CFEG |
| Detector | Falcon 4i | Falcon 4i | Falcon 4i |
| Energy filter slit (eV) | 10 | 10 | 10 |
| Magnification | 165,000 | 165,000 | 165,000 |
| Voltage (kV) | 300 | 300 | 300 |
| Electron dose (e-/Å$^2$) | 40 | 40 | 40 |
| Defocus range (μM) | 0.5 to 2.5 | 0.5 to 2.5 | 0.5 to 2.5 |
| Pixel size (Å) | 0.727 | 0.727 | 0.727 |
| **Data processing** | | | |
| Initial particle images (no.) | 80292 | 80292 | 80292 |
| Final particle images (no.) | 2111 | 4226 | 4013 |
| Helical twist (°) | -1.32 | -1.11 | -1.24 |
| Helical rise (Å) | 4.74 | 4.77 | 4.75 |
| Symmetry imposed | 1 | 1 | 1 |
| Map resolution FSC 0.143 (Å) | 3.72 | 3.28 | 3.21 |
| **Refinement** | | | |
| Initial model used (PDB code) | de novo | de novo | de novo |
| Model resolution FSC 0.5 (Å) | 3.8 | 3.6 | 3.30 |
| Map sharpening *B* factor (Å$^2$) | -47.87 | -37.31 | -33.55 |
| Model composition | | | |
| Non-hydrogen atoms | 3402 | 3402 | 3435 |
| Protein residues | 450 | 450 | 453 |
| Ligands | 0 | 0 | 0 |
| *B* factors (Å$^2$) | | | |
| Protein | 95.31 | 95.31 | 94.92 |
| Ligand | na | na | na |
| R.m.s. deviations | | | |
| Bond lengths (Å) | 0.011 | 0.010 | 0.011 |
| Bond angles (°) | 1.967 | 1.943 | 1.929 |
| Validation | | | |
| MolProbity score | 0.50 | 0.50 | 1.02 |
| Clashscore | 0 | 0 | 0.14 |
| Poor rotamers (%) | 0.51 | 0.00 | 0 |
| Ramachandran plot | | | |
| Favored (%) | 98.86 | 98.63 | 92.52 |
| Allowed (%) | 1.14 | 1.37 | 7.48 |
| Disallowed (%) | 0 | 0 | 0.00 |

| **LMB Krios G2** | **AD-MIA6** (240’ R1) (EMDB-xxxx) (PDB XXXX) | **AD-MIA7** (240’ R1) (EMDB-xxxx) (PDB XXXX) | **AD-LIA2** (240’ R1) (EMDB-xxxx) (PDB XXXX) |
|---|---|---|---|
| **Data acquisition** | | | |
| Electron gun | FEG | FEG | FEG |
| Detector | Falcon 4 | Falcon 4 | Falcon 4 |
| Energy filter slit (eV) | na | na | na |
| Magnification | 105,000 | 105,000 | 105,000 |
| Voltage (kV) | 300 | 300 | 300 |
| Electron dose (e-/Å$^2$) | 30 | 30 | 30 |
| Defocus range (μM) | 0.5 to 2.5 | 0.5 to 2.5 | 0.5 to 2.5 |
| Pixel size (Å) | 0.824 | 0.824 | 0.824 |
| **Data processing** | | | |
| Initial particle images (no.) | 352893 | 352893 | 352893 |
| Final particle images (no.) | 125479 | 7632 | 88413 |
| Helical twist (°) | -1.06 | -0.77 | -0.82 |
| Helical rise (Å) | 4.74 | 4.77 | 4.77 |
| Symmetry imposed | na | na | na |
| Map resolution FSC 0.143 (Å) | 2.82 | 3.26 | 2.95 |
| **Refinement** | | | |
| Initial model used (PDB code) | de novo | de novo | de novo |
| Model resolution FSC 0.5 (Å) | 3.1 | 3.9 | 3.0 |
| Map sharpening *B* factor (Å$^2$) | -72.91 | -45.89 | -78.18 |
| Model composition | | | |
| Non-hydrogen atoms | 3375 | 5109 | 3468 |
| Protein residues | 441 | 669 | 456 |
| Ligands | 0 | 0 | 0 |
| *B* factors (Å$^2$) | | | |
| Protein | 93.83 | 78.97 | 50.07 |
| Ligand | na | na | na |
| R.m.s. deviations | | | |
| Bond lengths (Å) | 0.010 | 0.010 | 0.010 |
| Bond angles (°) | 1.999 | 2.036 | 1.879 |
| Validation | | | |
| MolProbity score | 1.34 | 0.85 | 1.06 |
| Clashscore | 2.47 | 0.48 | 0.85 |
| Poor rotamers (%) | 0 | 0.17 | 0.25 |
| Ramachandran plot | | | |
| Favored (%) | 95.57 | 96.93 | 95.72 |
| Allowed (%) | 4.43 | 3.07 | 4.28 |
| Disallowed (%) | 0 | 0 | 0 |

| **TFS Krios G4** | **AD-MIA10** (240' R5) (EMDB-xxxx) (PDB XXXX) | **AD-MIA11** (240' R5) (EMDB-xxxx) (PDB XXXX) |
|---|---|---|
| **Data acquisition** | | |
| Electron gun | CFEG | CFEG |
| Detector | Falcon 4i | Falcon 4i |
| Energy filter slit (eV) | 10 | 10 |
| Magnification | 165,000 | 165,000 |
| Voltage (kV) | 300 | 300 |
| Electron dose (e-/$Å^2$) | 40 | 40 |
| Defocus range (μM) | 0.5 to 2.5 | 0.5 to 2.5 |
| Pixel size (Å) | 0.727 | 0.727 |
| **Data processing** | | |
| Initial particle images (no.) | 1178114 | 1178114 |
| Final particle images (no.) | 744002 | 17410 |
| Helical twist (°) | -1.08 | -0.99 |
| Helical rise (Å) | 4.74 | 4.76 |
| Symmetry imposed | 1 | 1 |
| Map resolution FSC 0.143 (Å) | 1.92 | 3.00 |
| **Refinement** | | |
| Initial model used (PDB code) | de novo | de novo |
| Model resolution FSC 0.5 (Å) | 2.9 | 3.6 |
| Map sharpening *B* factor ($Å^2$) | -32.0 | -54.81 |
| Model composition | | |
| Non-hydrogen atoms | 3246 | 4947 |
| Protein residues | 429 | 654 |
| Ligands | 0 | 0 |
| *B* factors ($Å^2$) | | |
| Protein | 96.32 | 95.98 |
| Ligand | na | na |
| R.m.s. deviations | | |
| Bond lengths (Å) | 0.011 | 0.010 |
| Bond angles (°) | 1.790 | 1.895 |
| Validation | | |
| MolProbity score | 0.78 | 0.80 |
| Clashscore | 0.91 | 0.69 |
| Poor rotamers (%) | 0 | 0 |
| Ramachandran plot | | |
| Favored (%) | 98.80 | 97.64 |
| Allowed (%) | 1.20 | 2.36 |
| Disallowed (%) | 0 | 0 |

| **LMB Krios G2** | **AD-LIA4** (300’ R4) (EMDB-xxxx) (PDB XXXX) |
|---|---|
| **Data acquisition** | |
| Electron gun | FEG |
| Detector | Falcon 4 |
| Energy filter slit (eV) | na |
| Magnification | 105,000 |
| Voltage (kV) | 300 |
| Electron dose (e-/Å$^2$) | 30 |
| Defocus range (μM) | 0.5 to 2.5 |
| Pixel size (Å) | 0.824 |
| **Data processing** | |
| Initial particle images (no.) | 100371 |
| Final particle images (no.) | 6032 |
| Helical twist (°) | 179.61 |
| Helical rise (Å) | 2.38 |
| Symmetry imposed | $2_1$ |
| Map resolution FSC 0.143 (Å) | 3.81 |
| **Refinement** | |
| Initial model used (PDB code) | de novo |
| Model resolution FSC 0.5 (Å) | 5.4 |
| Map sharpening *B* factor (Å$^2$) | -48.66 |
| Model composition | |
| Non-hydrogen atoms | 6810 |
| Protein residues | 894 |
| Ligands | 0 |
| *B* factors (Å$^2$) | |
| Protein | 93.55 |
| Ligand | na |
| R.m.s. deviations | |
| Bond lengths (Å) | 0.011 |
| Bond angles (°) | 2.082 |
| Validation | |
| MolProbity score | 1.81 |
| Clashscore | 5.47 |
| Poor rotamers (%) | 0 |
| Ramachandran plot | |
| Favored (%) | 91.03 |
| Allowed (%) | 9.97 |
| Disallowed (%) | 0 |

| LMB Krios G2 | AD-LIA6 (300’ R2) (EMDB-xxxx) (PDB XXXX) | AD-LIA5 (300’ R2) (EMDB-xxxx) (PDB XXXX) |
|---|---|---|
| **Data acquisition** | | |
| Electron gun | FEG | FEG |
| Detector | Falcon 4 | Falcon 4 |
| Energy filter slit (eV) | na | na |
| Magnification | 105,000 | 105,000 |
| Voltage (kV) | 300 | 300 |
| Electron dose (e-/Å$^2$) | 30 | 30 |
| Defocus range (μM) | 1.0 to 2.5 | 1.0 to 2.5 |
| Pixel size (Å) | 0.824 | 0.824 |
| **Data processing** | | |
| Initial particle images (no.) | 119480 | 119480 |
| Final particle images (no.) | 4959 | 9294 |
| Helical twist (°) | -1.07 | -1.07 |
| Helical rise (Å) | 4.766 | 4.766 |
| Symmetry imposed | na | na |
| Map resolution FSC 0.143 (Å) | 3.04 | 2.93 |
| **Refinement** | | |
| Initial model used (PDB code) | ModelAngelo | ModelAngelo |
| Model resolution FSC 0.5 (Å) | 3.4 | 3.4 |
| Map sharpening *B* factor (Å$^2$) | -27.03 | -30.40 |
| Model composition | | |
| Non-hydrogen atoms | 3468 | 3468 |
| Protein residues | 456 | 456 |
| Ligands | 0 | 0 |
| *B* factors (Å$^2$) | | |
| Protein | 92.45 | 92.45 |
| Ligand | na | na |
| R.m.s. deviations | | |
| Bond lengths (Å) | 0.01 | 0.01 |
| Bond angles (°) | 1.946 | 1.913 |
| Validation | | |
| MolProbity score | 0.66 | 0.93 |
| Clashscore | 0.00 | 0.14 |
| Poor rotamers (%) | 0 | 0.76 |
| Ramachandran plot | | |
| Favored (%) | 97.07 | 94.59 |
| Allowed (%) | 2.93 | 5.49 |
| Disallowed (%) | 0 | 0 |

| **LMB Krios G2** | **AD-LIA7**<br>(300’ R3)<br>(EMDB-xxxx)<br>(PDB XXXX) | **AD-PHFb**<br>(300’ R3)<br>(EMDB-xxxx)<br>(PDB XXXX) |
|---|---|---|
| **Data acquisition** | | |
| Electron gun | FEG | FEG |
| Detector | Falcon 4 | Falcon 4 |
| Energy filter slit (eV) | na | na |
| Magnification | 105,000 | 105,000 |
| Voltage (kV) | 300 | 300 |
| Electron dose (e-/Å$^2$) | 30 | 30 |
| Defocus range (μM) | 0.5 to 2.5 | 0.5 to 2.5 |
| Pixel size (Å) | 0.824 | 0.824 |
| **Data processing** | | |
| Initial particle images (no.) | 495698 | 495698 |
| Final particle images (no.) | 24999 | 109455 |
| Helical twist (°) | -1.05 | 179.86 |
| Helical rise (Å) | 4.77 | 2.394 |
| Symmetry imposed | 1 | $2_1$ |
| Map resolution FSC 0.143 (Å) | 3.04 | 2.95 |
| **Refinement** | | |
| Initial model used (PDB code) | de novo | de novo |
| Model resolution FSC 0.5 (Å) | 3.51 | 2.7 |
| Map sharpening *B* factor (Å$^2$) | -46.92 | -31.86 |
| Model composition | | |
| Non-hydrogen atoms | 3405 | 3468 |
| Protein residues | 447 | 456 |
| Ligands | 0 | 0 |
| *B* factors (Å$^2$) | | |
| Protein | 93.55 | 50.07 |
| Ligand | na | na |
| R.m.s. deviations | | |
| Bond lengths (Å) | 0.011 | 0.010 |
| Bond angles (°) | 2.141 | 2.029 |
| Validation | | |
| MolProbity score | 1.07 | 0.97 |
| Clashscore | 0.58 | 0.00 |
| Poor rotamers (%) | 0 | 0 |
| Ramachandran plot | | |
| Favored (%) | 94.48 | 92.57 |
| Allowed (%) | 5.29 | 7.43 |
| Disallowed (%) | 0.23 | 0 |

| **LMB Krios G2** | **AD-THF**<br>(360’ R1)<br>(EMDB-xxxx)<br>(PDB XXXX) |
|---|---|
| **Data acquisition** | |
| Electron gun | CFEG |
| Detector | Falcon 4i |
| Energy filter slit (eV) | 10 |
| Magnification | 165,000 |
| Voltage (kV) | 300 |
| Electron dose (e-/Å$^2$) | 40 |
| Defocus range (μM) | 0.5 to 2.5 |
| Pixel size (Å) | 0.727 |
| **Data processing** | |
| Initial particle images (no.) | 382080 |
| Final particle images (no.) | 60376 |
| Helical twist (°) | -0.86 |
| Helical rise (Å) | 4.75 |
| Symmetry imposed | 1 |
| Map resolution FSC 0.143 (Å) | 2.68 |
| **Refinement** | |
| Initial model used (PDB code) | de novo |
| Model resolution FSC 0.5 (Å) | 2.7 |
| Map sharpening *B* factor (Å$^2$) | -39.98 |
| Model composition | |
| Non-hydrogen atoms | 5202 |
| Protein residues | 684 |
| Ligands | 0 |
| *B* factors (Å$^2$) | |
| Protein | 50.03 |
| Ligand | na |
| R.m.s. deviations | |
| Bond lengths (Å) | 0.011 |
| Bond angles (°) | 2.224 |
| Validation | |
| MolProbity score | 1.22 |
| Clashscore | 0.85 |
| Poor rotamers (%) | 0 |
| Ramachandran plot | |
| Favored (%) | 92.79 |
| Allowed (%) | 7.21 |
| Disallowed (%) | 0.00 |

| **TFS Krios G4** | **AD-PHFa** (EMDB-xxxx) (PDB xxxx) |
|---|---|
| **Data acquisition** | |
| Electron gun | CFEG |
| Detector | Falcon 4i |
| Energy filter slit (eV) | 10 |
| Magnification | 165,000 |
| Voltage (kV) | 300 |
| Electron dose (e-/Å$^2$) | 40 |
| Defocus range (μM) | 0.5 to 2.5 |
| Pixel size (Å) | 0.727 |
| **Data processing** | |
| Initial particle images (no.) | 125865 |
| Final particle images (no.) | 72387 |
| Helical twist (°) | 179.494 |
| Helical rise (Å) | 2.38 |
| Symmetry imposed | $2_1$ |
| Map resolution FSC 0.143 (Å) | 2.1 |
| **Refinement** | |
| Initial model used (PDB code) | de novo |
| Model resolution FSC 0.5 (Å) | 2.2 |
| Map sharpening *B* factor (Å$^2$) | -50.2 |
| Model composition | |
| Non-hydrogen atoms | 3468 |
| Protein residues | 456 |
| Ligands | 0 |
| *B* factors (Å$^2$) | |
| Protein | 50.01 |
| Ligand | na |
| R.m.s. deviations | |
| Bond lengths (Å) | 0.011 |
| Bond angles (°) | 1.911 |
| Validation | |
| MolProbity score | 1.08 |
| Clashscore | 0.29 |
| Poor rotamers (%) | 0 |
| Ramachandran plot | |
| Favored (%) | 88.96 |
| Allowed (%) | 11.04 |
| Disallowed (%) | 0 |

| **LMB Krios G2** | **CTE-MIA1**<br>(160’ R4)<br>(EMDB-xxxx)<br>(PDB xxxx) |
|---|---|
| **Data acquisition** | |
| Electron gun | FEG |
| Detector | Falcon 4 |
| Energy filter slit (eV) | na |
| Magnification | 105,000 |
| Voltage (kV) | 300 |
| Electron dose (e-/Å$^2$) | 30 |
| Defocus range (μM) | 0.5 to 2.5 |
| Pixel size (Å) | 0.824 |
| **Data processing** | |
| Initial particle images (no.) | 59766 |
| Final particle images (no.) | 20639 |
| Helical twist (°) | -1.23 |
| Helical rise (Å) | 4.77 |
| Symmetry imposed | C2 |
| Map resolution FSC 0.143 (Å) | 3.3 |
| **Refinement** | |
| Initial model used (PDB code) | de novo |
| Model resolution FSC 0.5 (Å) | 3.5 |
| Map sharpening *B* factor (Å$^2$) | -52.74 |
| Model composition | |
| Non-hydrogen atoms | 3246 |
| Protein residues | 426 |
| Ligands | 0 |
| *B* factors (Å$^2$) | |
| Protein | 39.48 |
| Ligand | na |
| R.m.s. deviations | |
| Bond lengths (Å) | 0.011 |
| Bond angles (°) | 2.006 |
| Validation | |
| MolProbity score | 0.92 |
| Clashscore | 0.00 |
| Poor rotamers (%) | 0 |
| Ramachandran plot | |
| Favored (%) | 93.73 |
| Allowed (%) | 6.28 |
| Disallowed (%) | 0 |

| LMB Krios G2 | CTE-MIA3 (160’ R5) (EMDB-xxxx) (PDB xxxx) | CTE-MIA12 (160’ R5) (EMDB-xxxx) (PDB XXXX) |
|---|---|---|
| **Data acquisition** | | |
| Electron gun | FEG | CFEG |
| Detector | Falcon 4 | Falcon 4 |
| Energy filter slit (eV) | na | na |
| Magnification | 105,000 | 105,000 |
| Voltage (kV) | 300 | 300 |
| Electron dose (e-/Å$^2$) | 30 | 30 |
| Defocus range (μM) | 0.5 to 2.5 | 0.5 to 2.5 |
| Pixel size (Å) | 0.824 | 0.824 |
| **Data processing** | | |
| Initial particle images (no.) | 158080 | 158080 |
| Final particle images (no.) | 4243 | 44288 |
| Helical twist (°) | -1.13 | -1.39 |
| Helical rise (Å) | 4.77 | 4.77 |
| Symmetry imposed | 1 | C2 |
| Map resolution FSC 0.143 (Å) | 3.8 | 2.75 |
| **Refinement** | | |
| Initial model used (PDB code) | de novo | de novo |
| Model resolution FSC 0.5 (Å) | 4.1 | 2.4 |
| Map sharpening *B* factor (Å$^2$) | -25.39 | -34.96 |
| Model composition | | |
| Non-hydrogen atoms | 3444 | 1545 |
| Protein residues | 450 | 204 |
| Ligands | 0 | 0 |
| *B* factors (Å$^2$) | | |
| Protein | 94.85 | 97.44 |
| Ligand | na | na |
| R.m.s. deviations | | |
| Bond lengths (Å) | 0.011 | 0.01 |
| Bond angles (°) | 2.018 | 1.856 |
| Validation | | |
| MolProbity score | 1.39 | 0.89 |
| Clashscore | 1.99 | 0.63 |
| Poor rotamers (%) | 0 | 0 |
| Ramachandran plot | | |
| Favored (%) | 93.61 | 96.97 |
| Allowed (%) | 6.39 | 3.03 |
| Disallowed (%) | 0 | 0 |

| LMB Krios G2 | CTE-MIA4 (180’ R4) (EMDB-xxxx) (PDB XXXX) |
|---|---|
| **Data acquisition** | |
| Electron gun | FEG |
| Detector | Falcon 4 |
| Energy filter slit (eV) | na |
| Magnification | 105,000 |
| Voltage (kV) | 300 |
| Electron dose (e-/Å$^2$) | 30 |
| Defocus range (μM) | 1.0 to 2.5 |
| Pixel size (Å) | 0.824 |
| **Data processing** | |
| Initial particle images (no.) | 66426 |
| Final particle images (no.) | 42082 |
| Helical twist (°) | -1.09 |
| Helical rise (Å) | 4.73 |
| Symmetry imposed | 1 |
| Map resolution FSC 0.143 (Å) | 2.85 |
| **Refinement** | |
| Initial model used (PDB code) | de novo |
| Model resolution FSC 0.5 (Å) | 3.1 |
| Map sharpening *B* factor (Å$^2$) | -50.89 |
| Model composition | |
| Non-hydrogen atoms | 3246 |
| Protein residues | 429 |
| Ligands | 0 |
| *B* factors (Å$^2$) | |
| Protein | 96.32 |
| Ligand | na |
| R.m.s. deviations | |
| Bond lengths (Å) | 0.010 |
| Bond angles (°) | 1.863 |
| Validation | |
| MolProbity score | 0.82 |
| Clashscore | 0.45 |
| Poor rotamers (%) | 0.00 |
| Ramachandran plot | |
| Favored (%) | 97.12 |
| Allowed (%) | 2.88 |
| Disallowed (%) | 0 |

| TFS Krios G4 | CTE-MIA5 (180’ R1) (EMDB-xxxx) (PDB XXXX) | CTE-MIA6 (180’ R1) (EMDB-xxxx) (PDB XXXX) | CTE-MIA8 (180’ R1) (EMDB-xxxx) (PDB XXXX) | CTE-MIA18 (180’ R1) (EMDB-xxxx) (PDB XXXX) |
|---|---|---|---|---|
| **Data acquisition** | | | | |
| Electron gun | CFEG | CFEG | CFEG | CFEG |
| Detector | Falcon 4i | Falcon 4i | Falcon 4i | Falcon 4i |
| Energy filter slit (eV) | 10 | 10 | 10 | 10 |
| Magnification | 165,000 | 165,000 | 165,000 | 165,000 |
| Voltage (kV) | 300 | 300 | 300 | 300 |
| Electron dose (e-/Å$^2$) | 40 | 40 | 40 | 40 |
| Defocus range (μM) | 1.0 to 2.5 | 1.0 to 2.5 | 0.5 to 2.5 | 0.5 to 2.5 |
| Pixel size (Å) | 0.727 | 0.727 | 0.727 | 0.727 |
| **Data processing** | | | | |
| Initial particle images (no.) | 960020 | 960020 | 960020 | 960020 |
| Final particle images (no.) | 177452 | 12971 | 37333 | 27343 |
| Helical twist (°) | -1.08 | -1.12 | -1.31 | -1.65 |
| Helical rise (Å) | 4.74 | 4.71 | 4.75 | 4.74 |
| Symmetry imposed | 1 | 1 | C2 | 1 |
| Map resolution FSC 0.143 (Å) | 2.54 | 2.88 | 1.75 | 2.97 |
| **Refinement** | | | | |
| Initial model used (PDB code) | de novo | de novo | ModelAngelo | ModelAngelo |
| Model resolution FSC 0.5 (Å) | 2.9 | 3.1 | 2.7 | 3.1 |
| Map sharpening *B* factor (Å$^2$) | -65.69 | -50.06 | -19.86 | -65.20 |
| Model composition | | | | |
| Non-hydrogen atoms | 3435 | 3375 | 3444 | 3312 |
| Protein residues | 453 | 441 | 450 | 432 |
| Ligands | 0 | 0 | 0 | 0 |
| *B* factors (Å$^2$) | | | | |
| Protein | 93.54 | 93.83 | 96.09 | 94.68 |
| Ligand | na | na | na | na |
| R.m.s. deviations | | | | |
| Bond lengths (Å) | 0.011 | 0.011 | 0.01 | 0.01 |
| Bond angles (°) | 1.871 | 1.989 | 1.822 | 1.960 |
| Validation | | | | |
| MolProbity score | 0.87 | 0.68 | 0.50 | 0.92 |
| Clashscore | 0.57 | 0.29 | 0.00 | 1.18 |
| Poor rotamers (%) | 1.27 | 0 | 0 | 1.32 |
| Ramachandran plot | | | | |
| Favored (%) | 97.51 | 97.67 | 99.54 | 98.10 |
| Allowed (%) | 2.49 | 2.33 | 0.46 | 1.90 |
| Disallowed (%) | 0 | 0 | 0 | 0 |

| LMB Krios G2 | CTE-MIA7 (240’ R3) (EMDB-xxxx) (PDB XXXX) | CTE-MIA13 (240’ R3) (EMDB-xxxx) (PDB XXXX) | CTE-MIA14 (240’ R3) (EMDB-xxxx) (PDB XXXX) | CTE-MIA15 (240’ R3) (EMDB-xxxx) (PDB XXXX) |
|---|---|---|---|---|
| **Data acquisition** | | | | |
| Electron gun | FEG | FEG | FEG | FEG |
| Detector | Falcon 4 | Falcon 4 | Falcon 4 | Falcon 4 |
| Energy filter slit (eV) | na | na | na | na |
| Magnification | 105,000 | 105,000 | 105,000 | 105,000 |
| Voltage (kV) | 300 | 300 | 300 | 300 |
| Electron dose (e-/Å$^2$) | 30 | 30 | 30 | 30 |
| Defocus range (µM) | 1.0 to 2.5 | 1.0 to 2.5 | 1.0 to 2.5 | 1.0 to 2.5 |
| Pixel size (Å) | 0.824 | 0.824 | 0.824 | 0.824 |
| **Data processing** | | | | |
| Initial particle images (no.) | 208952 | 208952 | 208952 | 208952 |
| Final particle images (no.) | 23040 | 15874 | 14186 | 10660 |
| Helical twist (º) | -1.05 | 179.5 | -1.24 | -1.25 |
| Helical rise (Å) | 4.76 | 2.38 | 4.74 | 4.76 |
| Symmetry imposed | 1 | $2_1$ | 1 | 1 |
| Map resolution FSC 0.143 (Å) | 3.16 | 3.10 | 3.40 | 3.16 |
| **Refinement** | | | | |
| Initial model used (PDB code) | de novo | de novo | de novo | de novo |
| Model resolution FSC 0.5 (Å) | 3.3 | 3.1 | 3.8 | 3.2 |
| Map sharpening *B* factor (Å$^2$) | -53.84 | -48.26 | -33.30 | -45.23 |
| Model composition | | | | |
| Non-hydrogen atoms | 3444 | 3444 | 3444 | 3444 |
| Protein residues | 450 | 450 | 450 | 450 |
| Ligands | 0 | 0 | 0 | 0 |
| *B* factors (Å$^2$) | | | | |
| Protein | 94.85 | 94.85 | 94.85 | 94.85 |
| Ligand | na | na | na | na |
| R.m.s. deviations | | | | |
| Bond lengths (Å) | 0.010 | 0.010 | 0.010 | 0.010 |
| Bond angles (°) | 1.982 | 1.891 | 2.056 | 1.900 |
| Validation | | | | |
| MolProbity score | 0.82 | 0.78 | 0.90 | 1.11 |
| Clashscore | 0.00 | 0.00 | 0.00 | 0.85 |
| Poor rotamers (%) | 0.00 | 0.00 | 0.00 | 0.00 |
| Ramachandran plot | | | | |
| Favored (%) | 95.43 | 95.89 | 94.06 | 94.98 |
| Allowed (%) | 4.57 | 4.11 | 5.94 | 5.02 |
| Disallowed (%) | 0 | 0 | 0 | 0 |

| **LMB Krios G2** | **CTE-MIA9** (240’ R2) (EMDB-xxxx) (PDB XXXX) | **CTE-MIA10** (240’ R2) (EMDB-xxxx) (PDB XXXX) |
|---|---|---|
| **Data acquisition** | | |
| Electron gun | FEG | FEG |
| Detector | Falcon 4 | Falcon 4 |
| Energy filter slit (eV) | na | na |
| Magnification | 105,000 | 105,000 |
| Voltage (kV) | 300 | 300 |
| Electron dose (e-/Å$^2$) | 30 | 30 |
| Defocus range (μM) | 1.0 to 2.5 | 1.0 to 2.5 |
| Pixel size (Å) | 0.824 | 0.824 |
| **Data processing** | | |
| Initial particle images (no.) | 123830 | 123830 |
| Final particle images (no.) | 18265 | 38067 |
| Helical twist (°) | -1.14 | -1.30 |
| Helical rise (Å) | 4.77 | 4.77 |
| Symmetry imposed | C2 | 1 |
| Map resolution FSC 0.143 (Å) | 2.99 | 2.70 |
| **Refinement** | | |
| Initial model used (PDB code) | de novo | de novo |
| Model resolution FSC 0.5 (Å) | 3.1 | 2.5 |
| Map sharpening *B* factor (Å$^2$) | -44.91 | -42.43 |
| Model composition | | |
| Non-hydrogen atoms | 3246 | 3246 |
| Protein residues | 426 | 426 |
| Ligands | 0 | 0 |
| *B* factors (Å$^2$) | | |
| Protein | 39.48 | 39.48 |
| Ligand | na | na |
| R.m.s. deviations | | |
| Bond lengths (Å) | 0.011 | 0.010 |
| Bond angles (°) | 2.084 | 1.911 |
| Validation | | |
| MolProbity score | 0.50 | 0.63 |
| Clashscore | 0.00 | 0.00 |
| Poor rotamers (%) | 0.00 | 0.00 |
| Ramachandran plot | | |
| Favored (%) | 99.28 | 97.34 |
| Allowed (%) | 0.72 | 2.66 |
| Disallowed (%) | 0 | 0 |

| | |
|---|---|
| **LMB Krios G2** | **CTE-MIA11**<br>(240’ R6)<br>(EMDB-xxxx)<br>(PDB XXXX) |
| **Data acquisition** | |
| Electron gun | FEG |
| Detector | Falcon 4 |
| Energy filter slit (eV) | na |
| Magnification | 105,000 |
| Voltage (kV) | 300 |
| Electron dose (e-/$Å^2$) | 30 |
| Defocus range (µM) | 1.0 to 2.5 |
| Pixel size (Å) | 0.824 |
| **Data processing** | |
| Initial particle images (no.) | 234520 |
| Final particle images (no.) | 44009 |
| Helical twist (°) | -1.24 |
| Helical rise (Å) | 4.76 |
| Symmetry imposed | 1 |
| Map resolution FSC 0.143 (Å) | 3.04 |
| **Refinement** | |
| Initial model used (PDB code) | de novo |
| Model resolution FSC 0.5 (Å) | 3.2 |
| Map sharpening *B* factor ($Å^2$) | -71.45 |
| Model composition | |
| Non-hydrogen atoms | 3507 |
| Protein residues | 462 |
| Ligands | 0 |
| *B* factors ($Å^2$) | |
| Protein | 96.76 |
| Ligand | na |
| R.m.s. deviations | |
| Bond lengths (Å) | 0.010 |
| Bond angles (°) | 1.876 |
| Validation | |
| MolProbity score | 0.81 |
| Clashscore | 0.42 |
| Poor rotamers (%) | 0.00 |
| Ramachandran plot | |
| Favored (%) | 97.11 |
| Allowed (%) | 2.89 |
| Disallowed (%) | 0 |

| **TFS Krios G4** | **CTE-LIA3** (300’ R1) (EMDB-xxxx) (PDB XXXX) | **CTE-LIA4** (300’ R1) (EMDB-xxxx) (PDB XXXX) | **CTE-LIA5** (300’ R1) (EMDB-xxxx) (PDB XXXX) | **CTE-LIA6** (300’ R1) (EMDB-xxxx) (PDB XXXX) |
|---|---|---|---|---|
| **Data acquisition** | | | | |
| Electron gun | CFEG | CFEG | CFEG | CFEG |
| Detector | Falcon 4i | Falcon 4i | Falcon 4i | Falcon 4i |
| Energy filter slit (eV) | 10 | 10 | 10 | 10 |
| Magnification | 165,000 | 165,000 | 165,000 | 165,000 |
| Voltage (kV) | 300 | 300 | 300 | 300 |
| Electron dose (e-/Å$^2$) | 40 | 40 | 40 | 40 |
| Defocus range (μM) | 1.0 to 2.5 | 1.0 to 2.5 | 1.0 to 2.5 | 1.0 to 2.5 |
| Pixel size (Å) | 0.727 | 0.727 | 0.727 | 0.727 |
| **Data processing** | | | | |
| Initial particle images (no.) | 361062 | 361062 | 361062 | 361062 |
| Final particle images (no.) | 126250 | 48346 | 24805 | 18914 |
| Helical twist (°) | -1.45 | 179.389 | -1.2 | -1.48 |
| Helical rise (Å) | 4.74 | 2.374 | 4.76 | 4.76 |
| Symmetry imposed | 1 | $2_1$ | 1 | 1 |
| Map resolution FSC 0.143 (Å) | 1.93 | 2.18 | 2.65 | 2.56 |
| **Refinement** | | | | |
| Initial model used (PDB code) | de novo | de novo | de novo | de novo |
| Model resolution FSC 0.5 (Å) | 2.0 | 2.1 | 3.0 | 2.5 |
| Map sharpening *B* factor (Å$^2$) | -30.35 | -27.78 | -34.15 | -27.70 |
| Model composition | | | | |
| Non-hydrogen atoms | 3132 | 3342 | 3342 | 1671 |
| Protein residues | 414 | 438 | 438 | 219 |
| Ligands | 0 | 0 | 0 | 0 |
| *B* factors (Å$^2$) | | | | |
| Protein | 101 | 110 | 110 | 110 |
| Ligand | na | na | na | na |
| R.m.s. deviations | | | | |
| Bond lengths (Å) | 0.011 | 0.010 | 0.011 | 0.010 |
| Bond angles (°) | 1.872 | 1.835 | 2.049 | 1.860 |
| Validation | | | | |
| MolProbity score | 0.62 | 0.65 | 0.50 | 0.75 |
| Clashscore | 0.31 | 0.00 | 0.00 | 0.00 |
| Poor rotamers (%) | 0.00 | 0.00 | 0.00 | 0.00 |
| Ramachandran plot | | | | |
| Favored (%) | 99.75 | 97.18 | 98.36 | 96.24 |
| Allowed (%) | 0.25 | 2.82 | 1.64 | 3.76 |
| Disallowed (%) | 0 | 0 | 0 | 0 |

| **TFS Krios G4** | **CTE-LIA7** (300’ R1) (EMDB-xxxx) (PDB XXXX) | **CTE-LIA14** (300’ R1) (EMDB-xxxx) (PDB XXXX) |
|---|---|---|
| **Data acquisition** | | |
| Electron gun | CFEG | CFEG |
| Detector | Falcon 4i | Falcon 4i |
| Energy filter slit (eV) | 10 | 10 |
| Magnification | 165,000 | 165,000 |
| Voltage (kV) | 300 | 300 |
| Electron dose (e-/Å$^2$) | 40 | 40 |
| Defocus range (μM) | 1.0 to 2.5 | 1.0 to 2.5 |
| Pixel size (Å) | 0.727 | 0.727 |
| **Data processing** | | |
| Initial particle images (no.) | 361062 | 361062 |
| Final particle images (no.) | 5127 | 16784 |
| Helical twist (°) | -1.32 | 119.496 |
| Helical rise (Å) | 4.74 | 1.58 |
| Symmetry imposed | 1 | $3_1$ |
| Map resolution FSC 0.143 (Å) | 2.96 | 2.76 |
| **Refinement** | | |
| Initial model used (PDB code) | de novo | de novo |
| Model resolution FSC 0.5 (Å) | 2.9 | 2.4 |
| Map sharpening *B* factor (Å$^2$) | -30.73 | -43.56 |
| Model composition | | |
| Non-hydrogen atoms | 3342 | 5013 |
| Protein residues | 438 | 657 |
| Ligands | 0 | 0 |
| *B* factors (Å$^2$) | | |
| Protein | 110.13 | 110.93 |
| Ligand | na | na |
| R.m.s. deviations | | |
| Bond lengths (Å) | 0.011 | 0.010 |
| Bond angles (°) | 1.928 | 1.864 |
| Validation | | |
| MolProbity score | 0.50 | 0.50 |
| Clashscore | 0.00 | 0.00 |
| Poor rotamers (%) | 0.00 | 0.00 |
| Ramachandran plot | | |
| Favored (%) | 98.83 | 98.44 |
| Allowed (%) | 1.17 | 1.56 |
| Disallowed (%) | 0 | 0 |

| **LMB Krios G2** | **CTE-LIA13** (300’ R6) (EMDB-xxxx) (PDB XXXX) |
|---|---|
| **Data acquisition** | |
| Electron gun | FEG |
| Detector | Falcon 4 |
| Energy filter slit (eV) | na |
| Magnification | 105,000 |
| Voltage (kV) | 300 |
| Electron dose (e-/Å$^2$) | 30 |
| Defocus range (μM) | 1.0 to 2.5 |
| Pixel size (Å) | 0.824 |
| **Data processing** | |
| Initial particle images (no.) | 181708 |
| Final particle images (no.) | 15464 |
| Helical twist (°) | 179.353 |
| Helical rise (Å) | 2.39 |
| Symmetry imposed | $2_1$ |
| Map resolution FSC 0.143 (Å) | 3.20 |
| **Refinement** | |
| Initial model used (PDB code) | de novo |
| Model resolution FSC 0.5 (Å) | 3.0 |
| Map sharpening *B* factor (Å$^2$) | -37.66 |
| Model composition | |
| Non-hydrogen atoms | 3090 |
| Protein residues | 408 |
| Ligands | 0 |
| *B* factors (Å$^2$) | |
| Protein | 96.76 |
| Ligand | na |
| R.m.s. deviations | |
| Bond lengths (Å) | 0.010 |
| Bond angles (°) | 2.063 |
| Validation | |
| MolProbity score | 1.28 |
| Clashscore | 1.75 |
| Poor rotamers (%) | 0.00 |
| Ramachandran plot | |
| Favored (%) | 94.95 |
| Allowed (%) | 5.05 |
| Disallowed (%) | 0 |

| **LMB Krios G2** | **CTE-type I**<br>(720’ R1)<br>(EMDB-xxxx)<br>(PDB XXXX) |
|---|---|
| **Data acquisition** | |
| Electron gun | FEG |
| Detector | Falcon 4 |
| Energy filter slit (eV) | na |
| Magnification | 105,000 |
| Voltage (kV) | 300 |
| Electron dose (e-/Å$^2$) | 30 |
| Defocus range (μM) | 1.0 to 2.5 |
| Pixel size (Å) | 0.824 |
| **Data processing** | |
| Initial particle images (no.) | 36037 |
| Final particle images (no.) | 25924 |
| Helical twist (°) | 179.43 |
| Helical rise (Å) | 2.376 |
| Symmetry imposed | $2_1$ |
| Map resolution FSC 0.143 (Å) | 2.65 |
| **Refinement** | |
| Initial model used (PDB code) | de novo |
| Model resolution FSC 0.5 (Å) | 2.5 |
| Map sharpening *B* factor (Å$^2$) | -43.39 |
| Model composition | |
| Non-hydrogen atoms | 3444 |
| Protein residues | 450 |
| Ligands | 0 |
| *B* factors (Å$^2$) | |
| Protein | 115 |
| Ligand | na |
| R.m.s. deviations | |
| Bond lengths (Å) | 0.01 |
| Bond angles (°) | 1.859 |
| Validation | |
| MolProbity score | 0.82 |
| Clashscore | 1.14 |
| Poor rotamers (%) | 0.00 |
| Ramachandran plot | |
| Favored (%) | 98.17 |
| Allowed (%) | 1.83 |
| Disallowed (%) | 0 |

| **LMB Krios G2** | **CTE-type II**<br>(720’ R5)<br>(EMDB-xxxx)<br>(PDB XXXX) |
|---|---|
| **Data acquisition** | |
| Electron gun | FEG |
| Detector | Falcon 4 |
| Energy filter slit (eV) | na |
| Magnification | 105,000 |
| Voltage (kV) | 300 |
| Electron dose (e-/Å$^2$) | 30 |
| Defocus range (μM) | 1.0 to 2.5 |
| Pixel size (Å) | 0.824 |
| **Data processing** | |
| Initial particle images (no.) | 43926 |
| Final particle images (no.) | 19988 |
| Helical twist (°) | 179.262 |
| Helical rise (Å) | 2.39 |
| Symmetry imposed | $2_1$ |
| Map resolution FSC 0.143 (Å) | 3.1 |
| **Refinement** | |
| Initial model used (PDB code) | de novo |
| Model resolution FSC 0.5 (Å) | 2.9 |
| Map sharpening *B* factor (Å$^2$) | -49.83 |
| Model composition | |
| Non-hydrogen atoms | 3444 |
| Protein residues | 450 |
| Ligands | 0 |
| *B* factors (Å$^2$) | |
| Protein | 115 |
| Ligand | na |
| R.m.s. deviations | |
| Bond lengths (Å) | 0.011 |
| Bond angles (°) | 1.941 |
| Validation | |
| MolProbity score | 1.36 |
| Clashscore | 1.14 |
| Poor rotamers (%) | 3.03 |
| Ramachandran plot | |
| Favored (%) | 97.03 |
| Allowed (%) | 2.97 |
| Disallowed (%) | 0 |

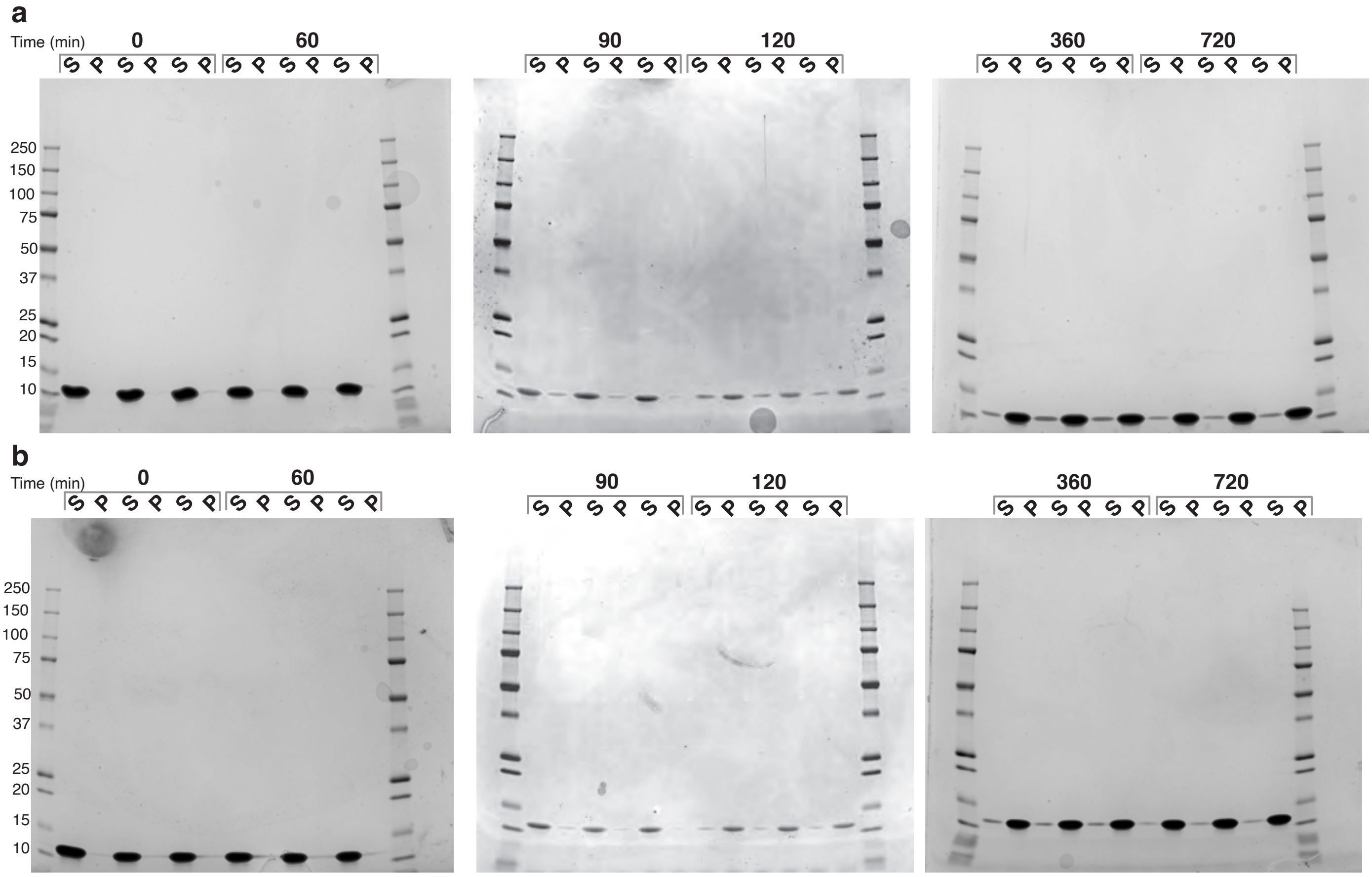
a
Time (min)
0
60
90
120
360
720
S P S P S P
250
150
100
75
50
37
25
20
15
10
b
Time (min)
0
60
90
120
360
720
S P S P S P
250
150
100
75
50
37
25
20
15
10
